# In vitro antimicrobial resistance in clinical isolates of bacterial pathogens causing bovine respiratory disease: a systematic review and meta-analysis

**DOI:** 10.64898/2026.09.28.754992

**Authors:** Qamer Mahmood, Philip Rasmussen, Peter Damborg, Kristine Berg Hansen, Camilla Bro Gregersen, Karolina Scahill, Luis Pedro Carmo, Clair L Firth, Bart Pardon, Maria Stokstad, Lise Marie Ånestad, Luca Guardabassi, Beate Conrady

## Abstract

**Background:** Bovine respiratory disease (BRD) is a major cause of antimicrobial use in cattle, raising concerns about antimicrobial resistance (AMR). This systematic review estimated pooled AMR proportions in BRD pathogens: *Mannheimia haemolytica, Pasteurella multocida*, and *Histophilus somni*, and assessed temporal and geographic trends in resistance.

**Methods:** Five antimicrobials commonly used for BRD treatment were evaluated: penicillin G (PEN), tetracycline (TET), florfenicol (FLO), tulathromycin (TUL), and enrofloxacin (ENR). Following PRISMA guidelines, 1,097 records were screened and 24 studies met inclusion criteria. Study quality and risk of bias were assessed, and random-effects meta-analyses estimated pooled proportions of resistant isolates by antimicrobial, continent and period.

**Results:** A total of 13,949 bacterial isolates were analysed including *P. multocida* (5,615), *M. haemolytica* (5,585), and *H. somni* (2,749). TET resistance was highest across all pathogens (12-30%), followed by TUL (6-11%), PEN (4-12%), FLO (0.5-1.6%) and ENR (0.2-1.1%). TET resistance in *M. haemolytica* and *P. multocida*, and FLO resistance in *P. multocida* were significantly more frequent in North America than in Europe (*p*≤0.05). TUL resistance increased significantly in *P. multocida* after 2013 (*p*=0.038), with similar non-significant trends in *M. haemolytica* and *H. somni*. FLO resistance decreased significantly in *H. somni* after 2013 (p=0.013). Egger’s test identified significant funnel-plot asymmetry for ENR resistance in *H. somni* and PEN resistance in *M. haemolytica*, suggesting possible small-study effects.

**Conclusion:** These results offer insights into distribution of phenotypic resistance across pathogens and regions, informing BRD antimicrobial treatment guidelines and One-Health antimicrobial stewardship aimed at preserving the effectiveness of medically important antimicrobials.

## 1. Introduction

Bovine respiratory disease (BRD) is a major cause of production losses, reduced animal welfare and increased antimicrobial consumption worldwide ^1-3^. This polymicrobial disease results from the complex interaction between viruses, bacteria, host and environmental factors affecting the respiratory tract of cattle ^4,5^. The major opportunistic pathogens involved are *Mannheimia haemolytica (M. haemolytica), Pasteurella multocida (P. multocida)*, and *Histophilus somni (H. somni)* ^6^. BRD is the primary reason for antimicrobial use in calves and youngstock ^7, 8^ including WHO-classified ‘highest priority critically important antimicrobials’ (HPCIA) classes such as fluoroquinolones and, to a lesser extent, extended-spectrum cephalosporins ^1, 9^. The use of antimicrobials for this indication is particularly high in feedlot and veal calf production systems worldwide ^1, 8^.

Although many calves appear to clinically recover following antimicrobial treatment, complete resolution of pulmonary lesions is often not achieved, resulting in persistent subclinical disease, chronicity, production losses, and compromised animal welfare ^10^. This has been demonstrated in the literature, where thoracic lung ultrasonography revealed that approximately 69% of male dairy calves remained subclinically affected after treatment, indicating a high rate of treatment failure ^11^. In the feedlot industry, based on clinical signs, treatment failure usually affects a small percentage of calves within a production system resulting in chronic illness and subsequent repeated treatments, salvage slaughter, euthanasia, or causes death, leading to overall negative impacts on costs and animal welfare ^12, 13^.

The extent to which treatment failure can be attributed to antimicrobial resistance (AMR) among bacterial respiratory pathogens remains unclear. Evidence from the literature suggests a decline in the susceptibility of BRD pathogens to commonly used antimicrobials. For instance, a review of 16 studies involving bacterial isolates from both clinically healthy and diseased feedlot cattle in North America reported a significant reduction in susceptibility to antimicrobial agents routinely used for

BRD treatment ^14^. Similarly, an assessment by the European Food Safety Authority (EFSA) identified moderate levels of resistance among BRD pathogens, while noting that clinical isolates generally remained susceptible to at least one therapeutic option commonly used for BRD treatment, including florfenicol, tulathromycin, and tilmicosin ^15^. Nevertheless, neither study quantitatively synthesized the available evidence through a formal meta-analysis.

The objective of this systematic review and meta-analysis was to estimate the pooled proportions of AMR in the BRD pathogens *M. haemolytica, P. multocida*, and *H. somni* from cattle (dairy, veal and beef). Furthermore, the review aimed to investigate geographical and temporal variation in resistance to five antimicrobial agents commonly used for the treatment of BRD: enrofloxacin (ENR), florfenicol (FLO), penicillin G (PEN), tetracycline (TET), and tulathromycin (TUL).

## 2. Methods

This systematic review was conducted according to the Preferred Reporting Items for Systematic Reviews and Meta-Analyses (PRISMA) 2020 guidelines provided in Supplementary Materials I ^16^. The review protocol was not published beforehand and is included in Supplementary Materials II.

### 2.1 Information sources and search strategy

The literature search was conducted in two online databases (PubMed and Embase), as previous research has shown that combining these databases increases the coverage of relevant publications by approximately 6.8%, thereby improving the comprehensiveness of systematic reviews ^17^.

An initial search was performed on 31 August 2023 and included studies published in English from 2010 up to the search date. To ensure that the review incorporated the most recent evidence, the search strategy was updated on 16 April 2026. This updated search was performed using the same search terms, eligibility criteria, and databases (PubMed and Embase) as the initial search.

For both searches, the search strategy was structured into three conceptual components: antimicrobial agents, host species, and bacterial species. The full search strings for both databases (PubMed and Embase) are provided in the review protocol in Supplementary Materials II.

### 2.2 Study selection

The study selection process followed a two-stage approach consisting of title and abstract screening followed by full text assessment. Three reviewers (and) conducted the screening, with CBG, KBH independently screening studies during the first phase and QM screening studies during the second phase. Each study was screened once by a single reviewer. Prior to screening, a calibration exercise was done on 15 randomly selected studies to increase standardization between reviewers. The inclusion criteria were comprised of original research articles (i) reporting bacterial isolates obtained from clinical respiratory disease cases in cattle, (ii) providing minimum inhibitory concentration data interpreted using Clinical and Laboratory Standards Institute (CLSI) clinical breakpoints ^18^, and (iii) published in English. Full eligibility criteria applied during screening are provided in the review protocol in Supplementary Materials II.

### 2.3 Data extraction

For each eligible study, data were extracted into Microsoft Excel, including the first author and year of publication, study design, continent or geographical location of the study, sample collection period, temporal classification period, production type (beef, dairy, both), bacterial pathogen investigated (*M. haemolytica, P. multocida*, and/or *H. somni*), antimicrobial agent tested (ENR, FLO, PEN, TET, and/or TUL), and antimicrobial class. In addition, the total number of isolates tested for antimicrobial susceptibility (N) and the corresponding susceptibility outcomes were recorded, including the proportion and number of resistant isolates (R% and N(R)), intermediate isolates (I% and N(I)), and susceptible isolates (S% and N(S)). Where reported, combined resistant and intermediate proportions (R+I%) and combined susceptible and intermediate proportions (S+I%) were also extracted. When susceptibility results were reported as percentages only, the corresponding numbers of isolates were calculated using the reported sample size whenever possible. Similarly, when only isolate counts were reported, percentages were calculated based on the total number of isolates tested for the respective antimicrobial agent. The data extraction tool is provided in Supplementary Materials III.

### 2.4 Quality appraisal and risk of bias assessment

Methodological quality and risk of bias within included studies were assessed using a 20-point appraisal tool developed by Downes et al. ^19^. The evaluation focused on clarity of objectives, appropriateness of study design, representativeness of the sample frame, and statistical validity. Criteria were scored as ‘Yes’, ‘No’, or ‘Do not know’ based on the presence and transparency of information reported in the full texts. As the appraisal tool does not generate a composite study-level risk-of-bias score, no risk-of-bias-based meta-analyses or sensitivity analyses were conducted.

Additionally, funnel-plot asymmetry and potential small-study effects were evaluated for analyses involving at least 10 studies. This was performed via visual inspection of funnel plots and Egger’s linear regression test ^20, 21^, where a *p*-value < 0.05 indicated statistically significant asymmetry.

### 2.5 Meta analysis

Meta-analyses were performed to estimate the pooled proportion of isolates resistant to each of the five antimicrobials for each pathogen. Resistant (R) isolates were treated as events; intermediate (I) and susceptible (S) isolates as non-events. Multiple records from the same study for a given pathogen–antimicrobial combination were combined so that each study contributed a single estimate per analysis.

Proportions were pooled on the logit scale using a random-effects, random-intercept logistic regression model (generalized linear mixed model), with between-study variance (τ^2^) estimated by maximum likelihood. Confidence intervals for the pooled proportion were derived using the Hartung-Knapp adjustment, and a 95% prediction interval was calculated for each estimate. Heterogeneity was assessed using Cochran’s Q test (P ≤ 0.05 indicating heterogeneity), the I^2^ statistic, and τ^2^, with I^2^ > 70% interpreted as substantial heterogeneity ^22^. Pooled estimates were back transformed to proportions and are reported with 95% confidence intervals (CI).

Analyses were performed overall and, where data permitted, stratified by continent and by sampling period; differences between subgroups were evaluated using the random-effects test for between-subgroup heterogeneity. The sampling period cutoff (2013) was derived from the data rather than chosen arbitrarily, corresponding to the median collection period midpoint across the 24 included studies. Studies whose reported collection period straddled this cutoff were excluded from the period-stratified analysis only and were retained in the overall and continent-stratified analyses. At least three independent studies were required to pool an overall estimate and three per group for a subgroup estimate ^21, 23^. Analyses were conducted in R (Version 4.4.2; R Core Team, 2026) using the metaprop, forest, funnel, and metabias functions of the meta package version [8.2.0] ^24^.

To address differences in resistance across study designs, a sensitivity analysis was conducted. Overall pooled estimates were re-estimated restricted to cross-sectional studies only (the overwhelmingly dominant study design; 19 of 24), for pathogen-antimicrobial combinations with at least three eligible cross-sectional studies, to assess whether the inclusion of controlled trial and *in vitro* laboratory studies influenced the pooled resistance estimates. Because BRD treatment is commonly selected empirically and is expected to provide coverage across the main bacterial pathogens, pooled estimates were also summarised at the antimicrobial level across *H. somni, M. haemolytica* and *P. multocida*.

The clinical question of the present study was: for each antimicrobial commonly used for BRD treatment, what is the phenotypic resistance profile across the main BRD-associated bacterial pathogens isolated from cattle with clinical respiratory disease?

### 2.6 Certainty assessment

No formal GRADE guidance exists specifically for proportional meta-analysis. Therefore, certainty in the antimicrobial-level resistance profile was assessed using a GRADE-informed approach, applying standard GRADE principles while considering methodological guidance for proportional meta-analysis ^25, 26^. Certainty was initially considered high for the descriptive question of phenotypic resistance proportions and was downgraded based on risk of bias, inconsistency, imprecision, indirectness and publication bias. Risk of bias judgements were informed by the quality appraisal described above, with particular emphasis on representativeness of isolate selection, clarity of the sampling frame and validity of outcome measurement.

Prespecified pragmatic resistance thresholds were used to support the interpretation of both inconsistency and imprecision, defining the following interpretive ranges: very low <10%, low 10– <20%, moderate 20–<30% and high resistance ≥30%. These thresholds were not intended as validated thresholds for treatment decisions, but as interpretive ranges for judging certainty in pooled AMR proportions. Inconsistency was assessed by visual inspection of forest plots, considering whether study-specific point estimates fell within the same or different prespecified resistance ranges. I^2^ was considered a descriptive indicator of heterogeneity but was not used as the sole basis for downgrading, because I^2^ was developed for comparative meta-analysis and can be difficult to interpret in proportional meta-analysis ^25, 27^. Imprecision was assessed using the 95% confidence intervals around the pooled resistance estimates, considering how many prespecified resistance thresholds were crossed. Indirectness was assessed in relation to the predefined in vitro resistance question, considering whether the population, pathogens, antimicrobial susceptibility testing approach and resistance outcome aligned with the question of interest. Publication bias was assessed qualitatively, considering review methods, characteristics of the evidence base and funnel plot asymmetry where available. Funnel plot asymmetry and Egger’s tests were interpreted cautiously because these methods are not well established for proportional meta-analysis ^25^. Where inconsistency and imprecision overlapped, particularly because random-effects confidence intervals incorporate between study heterogeneity, final certainty judgements avoided double-counting the same underlying uncertainty.

## 3. Results

### 3.1 Study selection

The database search, conducted in two combined literature search rounds, identified 1,097 records, including 681 records from Embase and 416 from PubMed. After removal of 323 duplicates, 774 records underwent title and abstract screening, resulting in the exclusion of 595 records. The full texts of the remaining 179 articles were assessed for eligibility, of which 155 were excluded based on the predefined inclusion criteria. Consequently, 24 studies were included in the final qualitative synthesis and meta-analysis (Figure 1).

**Figure 1.**
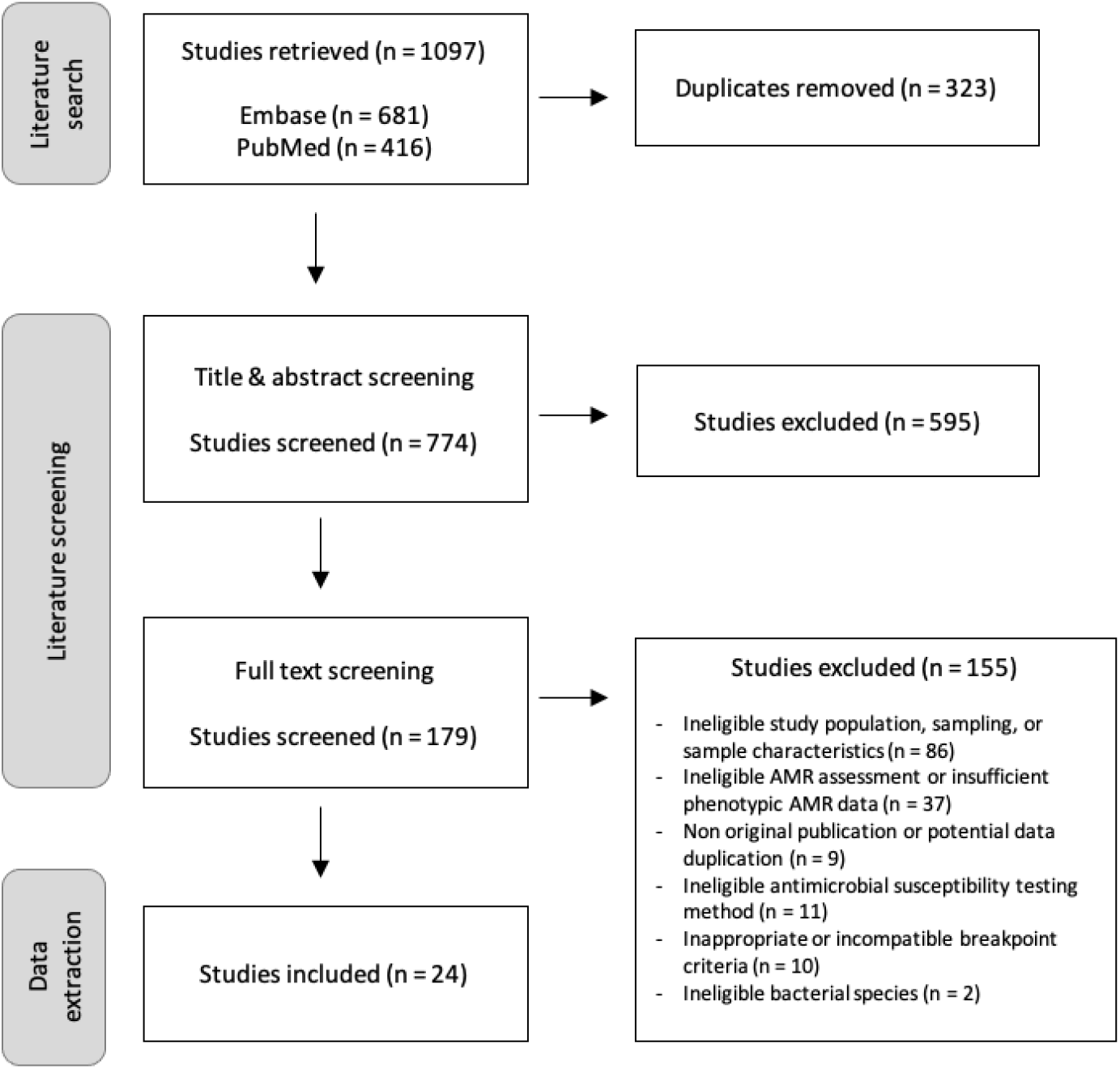
PRISMA flow diagram illustrating the selection process of included studies in the systematic review ^16^. A detailed exclusion criteria applied at full text screening is provided in Supplementary Materials II.

### 3.2 Study characteristics

From the 24 included studies, a total of 13,949 bacterial isolates were analysed across the three bovine respiratory pathogens: *P. multocida* (5,615 isolates), *M. haemolytica* (5,585 isolates), and *H. somni* (2,749 isolates). Research focus varied among production systems, with beef only systems being the most frequently investigated (n=12 studies; comprising 8 feedlot operations, 3 other beef producing systems, and 1 mixed beef operation), followed by multi-system studies evaluating combined beef and dairy sectors (n = 8 studies, of which 2 also included veal calves). Traditional dairy-only systems were targeted in 3 studies, while veal calf-only systems were evaluated in 2 studies. Study-level collection periods were classified relative to 2013 (the median collection period midpoint across included studies): 11 studies were classified as after 2013, 8 studies as before 2013, and 5 studies whose collection period straddled the cutoff were excluded from the period-stratified analysis (retained in the overall and continent-stratified analyses). Most of the studies were conducted in North America (11 studies) and Europe (9 studies), with additional representation from Asia (3 studies) and Australia (1 study). Regarding study design, 19 studies were cross-sectional while 5 studies were experimental (3 controlled trial, 2 in vitro laboratory studies) (Table 1).

**Table 1.**
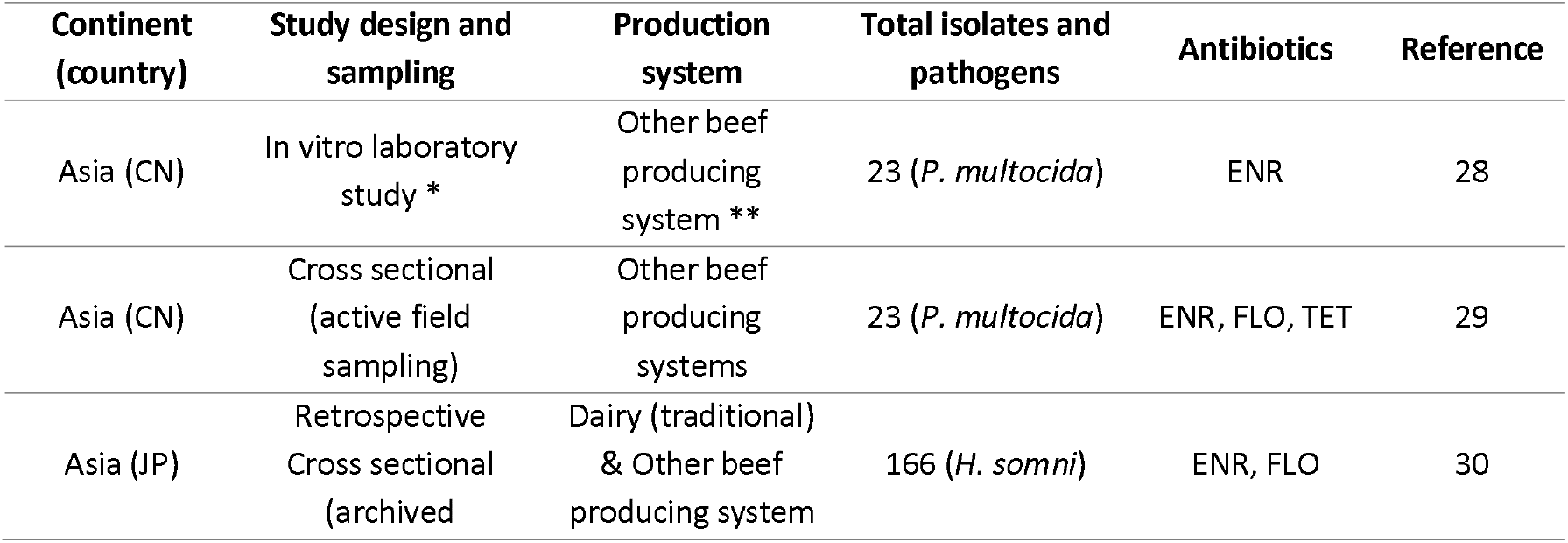
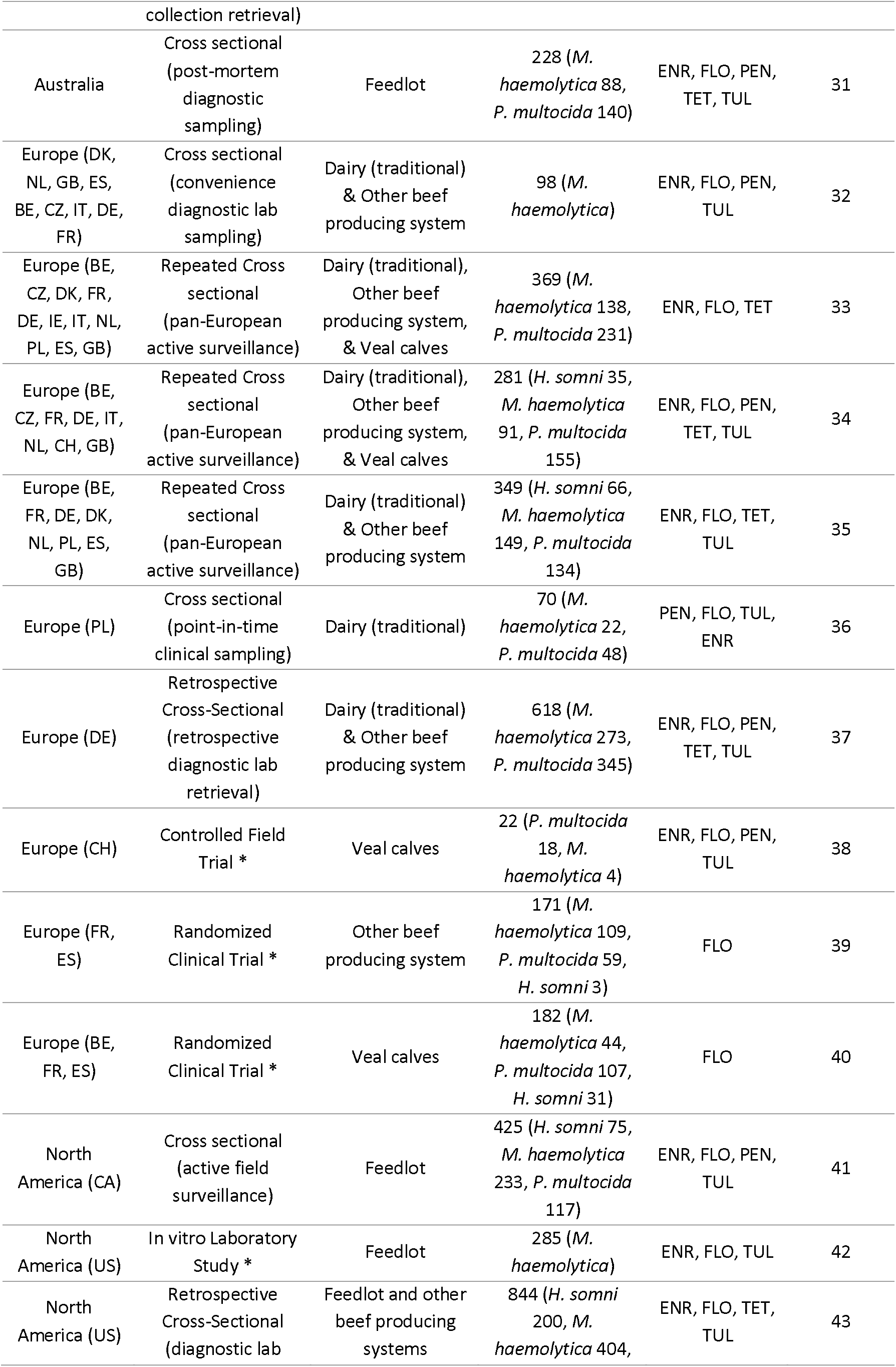
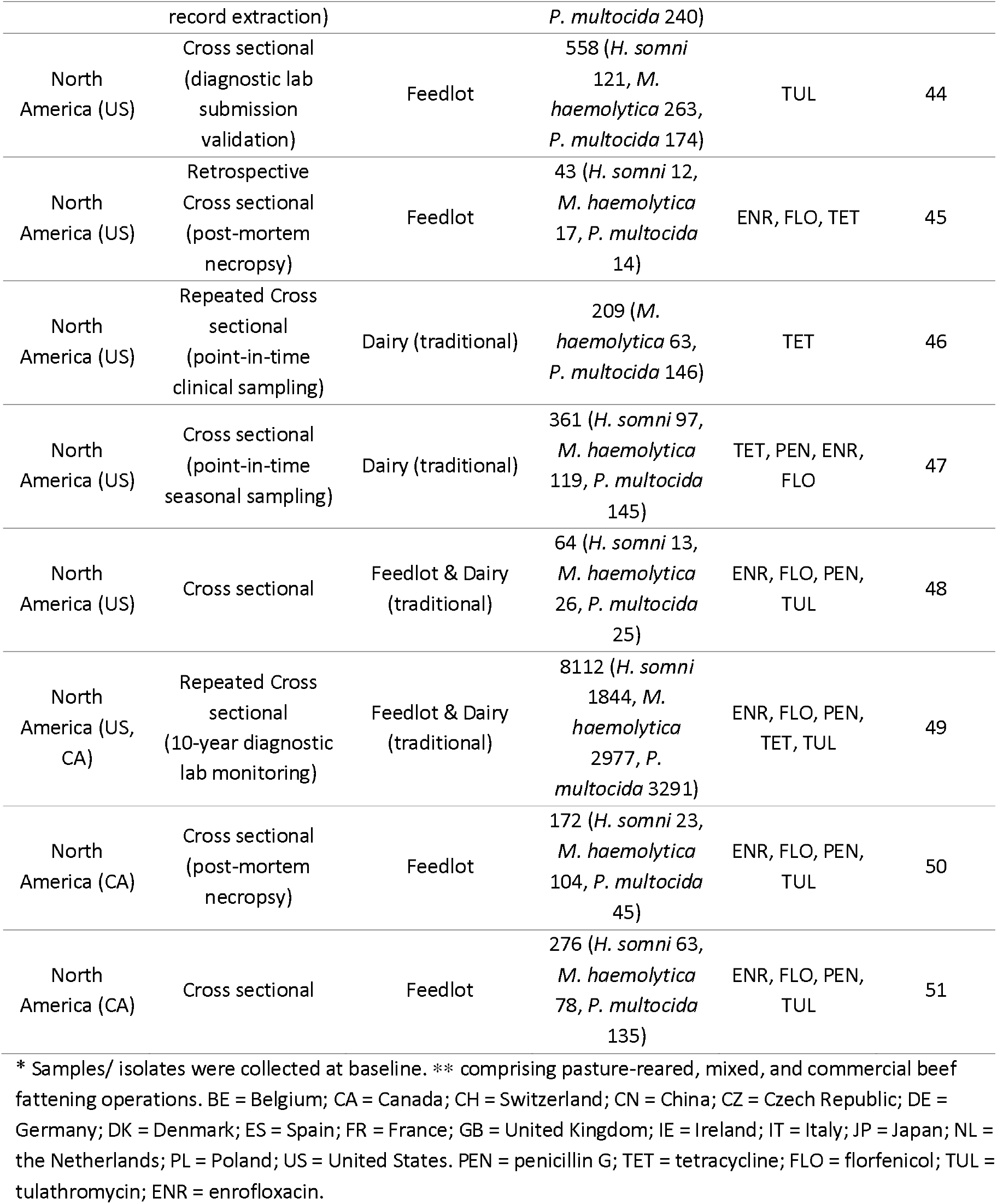
Summary of key characteristics of the studies included in systematic review.

### 3.3 Quality appraisal and risk of bias assessment

The quality appraisal and publication bias was accessed for the included studies and are reported below.

#### 3.3.1 Quality appraisal across included studies

The methodological quality of the 24 included studies varied across appraisal domains. Objectives and study designs were generally clearly reported and appropriate, but recurrent limitations concerned sample representativeness, sample-size justification and reporting of potential selection or non-response bias. In addition, a considerable risk of non-response bias was identified in retrospective surveillance and diagnostic laboratory-based studies ^37, 43^, as these studies relied on passive laboratory submissions and provided limited information on the underlying eligible population and samples or isolates that were not represented in the analysed dataset. While most studies (92%) adequately discussed their limitations, explicit reporting of ethical approval or participant consent was missing in 40% (n=10) of the set, particularly in pre-2013 studies utilising secondary diagnostic data, as ethics approval may not have been sought in the past. Detailed evaluations for each criterion across all included studies are documented in Supplementary Materials II. Concerns regarding isolate selection and representativeness informed the risk-of-bias judgements in the GRADE-informed certainty assessment below.

#### 3.3.2 Publication bias

Publication bias was assessed for the 12 of 15 pathogen-antimicrobial combinations represented by data from at least 10 studies (all except PEN, TET, and TUL in *H. somni*). Egger’s test indicated significant funnel-plot asymmetry for ENR resistance in *H. somni* (*p*=0.002) and PEN resistance in *M. haemolytica* (*p*=0.003) and in *P. multocida* (*p*<0.001 and *p*=0.005) (Figure 2), while no significant asymmetry was detected for the remaining combinations. Funnel plots and test statistics for all evaluable combinations are provided in Supplementary Materials IV. These findings were interpreted cautiously because funnel plot asymmetry and Egger’s tests are difficult to interpret in proportional meta-analysis and may reflect heterogeneity, sparse events or sampling differences rather than publication bias. The implications of these findings for the certainty of evidence are further considered in the GRADE-informed assessment below.

**Figure 2.**
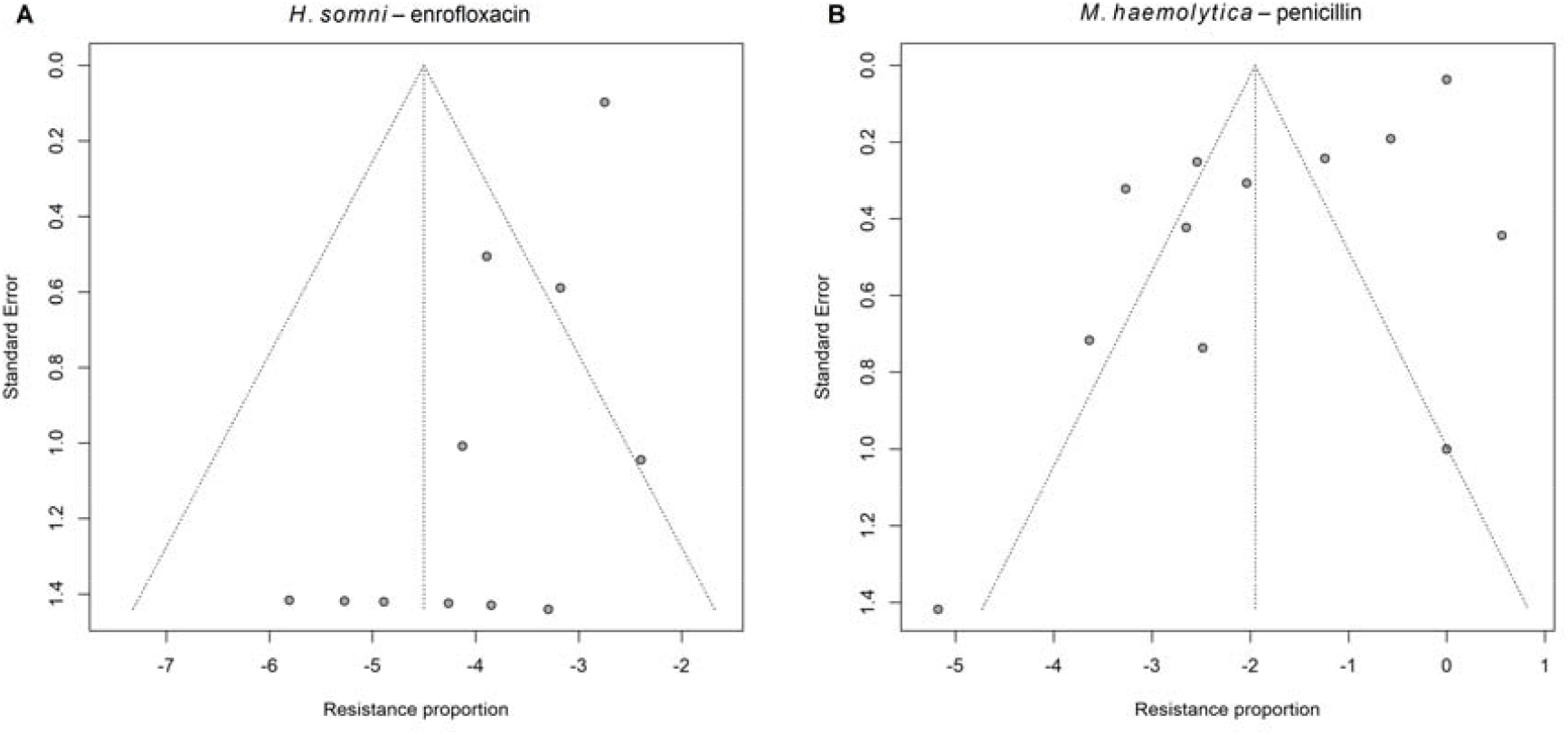
Funnel plot symmetry by linear regression test and Egger’s test results. **(A)** Funnel plot for enrofloxacin (ENR) resistance in *H. somni* where t = -4.23, df = 9, p-value = 0.002, Bias estimate: -1.273 (SE=0.301), multiplicative residual heterogeneity variance - tau^2 = 0.683, predictor: standard error, weight: inverse variance. **(B)** Funnel plot for penicillin (PEN) resistance in *M. haemolytica* where t = -3.93, df = 10, *p*-value = 0.003, Bias estimate: -5.044 (SE = 1.284), multiplicative residual heterogeneity variance - tau^2 = 13.613, predictor: standard error, weight: inverse variance.

### 3.4 Meta-analysis of pooled proportion of resistant isolates and antimicrobial resistance profiles

Across the three bacterial pathogens, TET resistance was the most prevalent with pooled mean proportions ranging from 12% to 30%, while lower resistance percentages were observed for TUL (6– 11%), PEN (4-12%), ENR (0.2-1.1%), and FLO (0.6-1.6%). The AMR landscape for the three BRD pathogens across the five antimicrobial classes are visually summarized in the pooled resistance heatmap (Figure 3).

**Figure 23.**
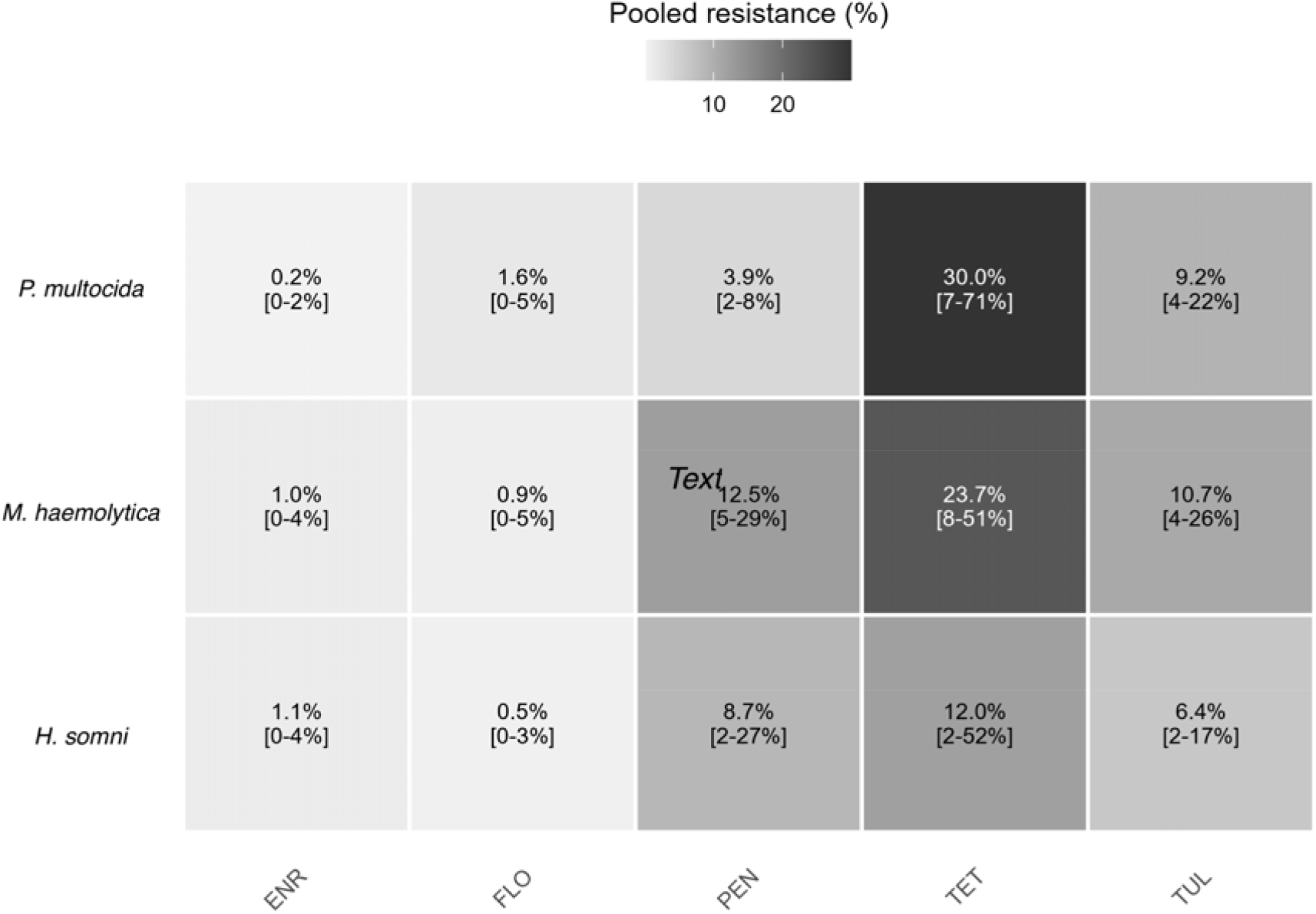
Heatmap of pooled percentage of resistant isolates for bovine respiratory pathogens. Each title shows the pooled point estimate and 95% confidence interval. The intensity of the shading represents the magnitude of resistance, with the highest percentage observed for tetracycline in *P. multocida* (30.0%) and *M. haemolytica* (23.7%). PEN = penicillin G; TET = tetracycline; FLO = florfenicol; TUL = tulathromycin; ENR = enrofloxacin.

The detailed results of the meta-analyses for AMR in *H. somni, M. haemolytica*, and *P. multocida*, stratified by period and continent, are presented in Table 2. Due to the limited number of studies from Asia (n = 3) and Australia (n = 1), the continent-specific subgroup analyses were restricted to Europe and North America. Resistance to TET in *M. haemolytica* (North America 48.1% vs. Europe 16.6%, *p*=0.008), TET in *P. multocida* (North America 73.6% vs. Europe 13.7%, *p*=0.003), and to FLO in *P. multocida* (North America 6.3% vs. Europe 0.9%, *p*=0.024) was significantly higher in North America than in Europe. Point estimates were also higher in North America for several other combinations, including FLO and TUL in *M. haemolytica* and TUL in *P. multocida*, but these between-continent differences did not reach statistical significance. When stratified by sampling period, TUL resistance in *P. multocida* was significantly higher after 2013 than before (17.0% vs. 3.1%, p=0.038), while FLO resistance *H. somni* was significantly lower after 2013 than before (0.6% vs. 2.8%, *p*=0.013). Point estimates were also higher after 2013 for TUL resistance in *M. haemolytica* (2.6% to 17.5%) and *H. somni* (0.9% to 7.4%), but these differences were not statistically significant.

**Table 2.** Meta-analysis of pooled mean proportions of antimicrobial resistance in *H. somni, M. haemolytica*, and *P. multocida* including 95% confidence intervals (CI), stratified by sampling period and continent. ENR= enrofloxacin; FLO= florfenicol; PEN= penicillin; TET= tetracycline; TUL= tulathromycin. Bold text indicates statistically significant differences (*p*<0.05) between subgroups. *I*^2^(%) represents the index of statistical heterogeneity, quantifying the proportion of total variation across studies that is due to heterogeneity rather than chance. Dashes (–) indicate that data were insufficient for that specific subgroup analysis.

| | Overall<br>Mean<br>proportion<br>%; 95% CI in<br>bracket | Before 2013<br>Mean<br>proportion %;<br>95% CI in<br>bracket | After 2013<br>Mean<br>proportion %;<br>95% CI in<br>bracket | North<br>America<br>Mean<br>proportion %;<br>95% CI in<br>bracket | Europe<br>Mean<br>proportion %;<br>95% CI in<br>bracket | $I^2(\%)$ |
| --- | --- | --- | --- | --- | --- | --- |
| ***H. somni*** |  |  |  |  |  |  |
| FLO | 0.5<br>[0.1;2.6] | 2.8<br>[2.0;4.1] | 0.6<br>[0.1;2.7] | 1.0<br>[0.2;4.4] | 0.0<br>[0.0;100.0] | 0 |
| ENR | 1.1 [0.3;4.4] | 3.0 [0.2;34.3] | 1.6 [0.7;4.0] | 2.4 [0.9;6.5] | - | 0 |
| PEN | 8.7 [2.4;26.8] | - | 8.9 [1.1;46.5] | 11.6<br>[3.5;32.0] | - | 94 |
| TUL | 6.4 [2.3;16.6] | 0.9 [0.0;99.7] | 7.4 [2.4;20.6] | 11.2<br>[6.4;18.8] | - | 63 |
| TET | 12.0<br>[1.7;51.9] | 11.4<br>[0.3;85.2] | 11.3<br>[0.0;99.0] | 29.7<br>[6.9;70.7] | - | 94 |
| ***M. haemolytica*** |  |  |  |  |  |  |
| FLO | 0.9 [0.2;4.6] | 0.1 [0.0;39.0] | 2.6 [0.7;9.1] | 3.5 [0.5;20.0] | 0.4 [0.0;4.6] | 88 |
| ENR | 1.0 [0.3;3.8] | 0.2 [0.0;11.5] | 2.0 [0.4;10.0] | 1.5 [0.1;13.3] | 0.9 [0.4;2.3] | 91 |
| PEN | 12.5<br>[4.7;29.0] | 30.2<br>[2.0;90.1] | 8.1 [2.0;27.6] | 13.8<br>[4.0;38.4] | 19.0<br>[3.4;60.8] | 97 |
| TUL | 10.7<br>[3.9;26.1] | 2.6 [0.1;35.5] | 17.5<br>[4.5;49.1] | 17.8<br>[4.1;52.2] | 7.5 [1.6;28.6] | 96 |
| TET | 23.7<br>[8.5;51.1] | 30.3<br>[7.0;71.5] | 18.4<br>[2.3;68.2] | **48.1**<br>**[16.1;81.7]** | **16.6**<br>**[11.0;24.1]** | 98 |
| ***P. multocida*** |  |  |  |  |  |  |
| FLO | 1.6 [0.4;5.4] | 0.8 [0.0;19.0] | 2.4 [0.5;10.2] | **6.3 [1.5;23.3]** | **0.9 [0.2;4.0]** | 94 |
| ENR | 0.2 [0.0;2.4] | 0.9 [0.1;13.6] | 0.2 [0.0;6.5] | 0.4 [0.0;18.7] | 0.4 [0.0;4.1] | 96 |
| PEN | 3.9 [1.8;8.4] | 7.0 [5.3;9.1] | 3.1 [1.0;9.4] | 3.4 [0.9;12.3] | 4.5 [0.6;28.8] | 73 |
| TUL | 9.2 [3.5;22.0] | **3.1 [0.3;23.6]** | **17.0**<br>**[5.4;42.4]** | 14.1<br>[3.2;45.2] | 4.6 [0.7;24.0] | 97 |
| TET | 30.0<br>[7.0;70.8] | 21.7<br>[3.6;67.2] | 52.4<br>[4.7;96.1] | **73.6**<br>**[7.3;99.0]** | **13.7**<br>**[3.7;39.2]** | 97 |

As a sensitivity analysis restricted to cross-sectional studies, pooled estimates were re-estimated for the nine pathogen-antimicrobial combinations with at least three eligible cross-sectional studies (FLO in *H. somni*; ENR, FLO, PEN, and TUL in *M. haemolytica*; and ENR, FLO, PEN, and TUL in *P. multocida*). This restriction to cross-sectional studies produced pooled estimates comparable to the overall analyses, with confidence intervals overlapping in all cases. The largest shift in point estimate was observed for TUL resistance in *M. haemolytica* (10.7% overall vs 15.3% when restricted to cross-sectional), while the remaining combinations shifted by less than 2 percentage points. Detailed results of the study design sensitivity analyses are provided in Supplementary Materials V.

Additionally, high levels of statistical heterogeneity were observed for most pooled resistance estimates (typically I^2^ > 70%), likely reflecting differences in production systems, geographic regions, study populations, and methodological approaches, although I^2^ can be difficult to interpret reliably in proportional meta-analyses. Random-effects models were used *a priori*, as described in the Section 2.5 of the Methods, to account for this between-study variability and to generate more conservative and robust pooled resistance estimates with improved generalizability.

To visualize the distribution of resistance proportions across individual studies and their relative weights in the meta-analysis, forest plots were generated for each pathogen-antimicrobial combination. Figures 4, 5 and 6 present the forest plot for the proportion of TET resistance in *H. somni*, TET resistance in *M. haemolytica* and FLO resistance in *P. multocida*, respectively. Forest plots for the remaining pathogen-antimicrobial combinations are provided in Supplementary Materials VI. Forest plots stratified by geographical region and study period were also generated for each pathogen-antimicrobial combination and are provided in Supplementary Materials VII.

**Figure 4.**
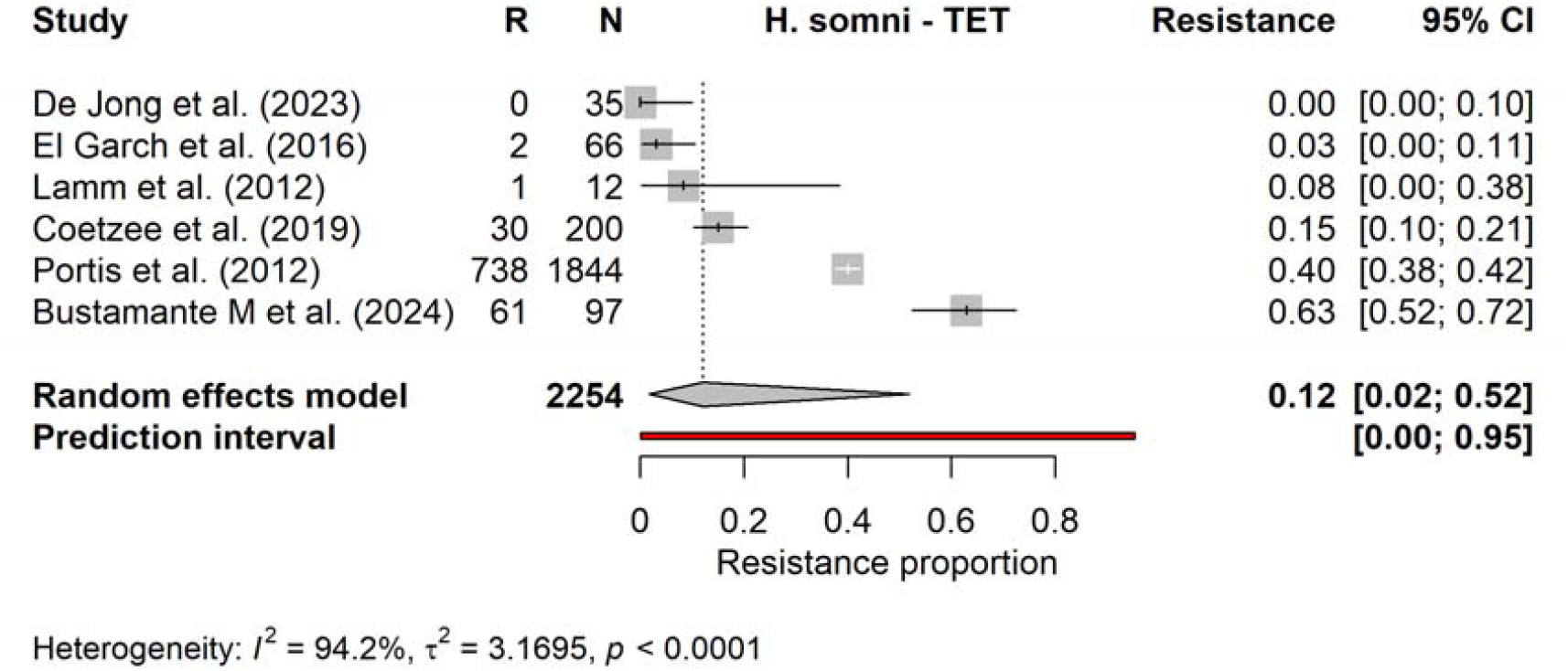
Forest plot of the pooled proportion of resistant isolates to tetracycline (TET) in *Histophilus somni* (*H. somni*).

**Figure 5.**
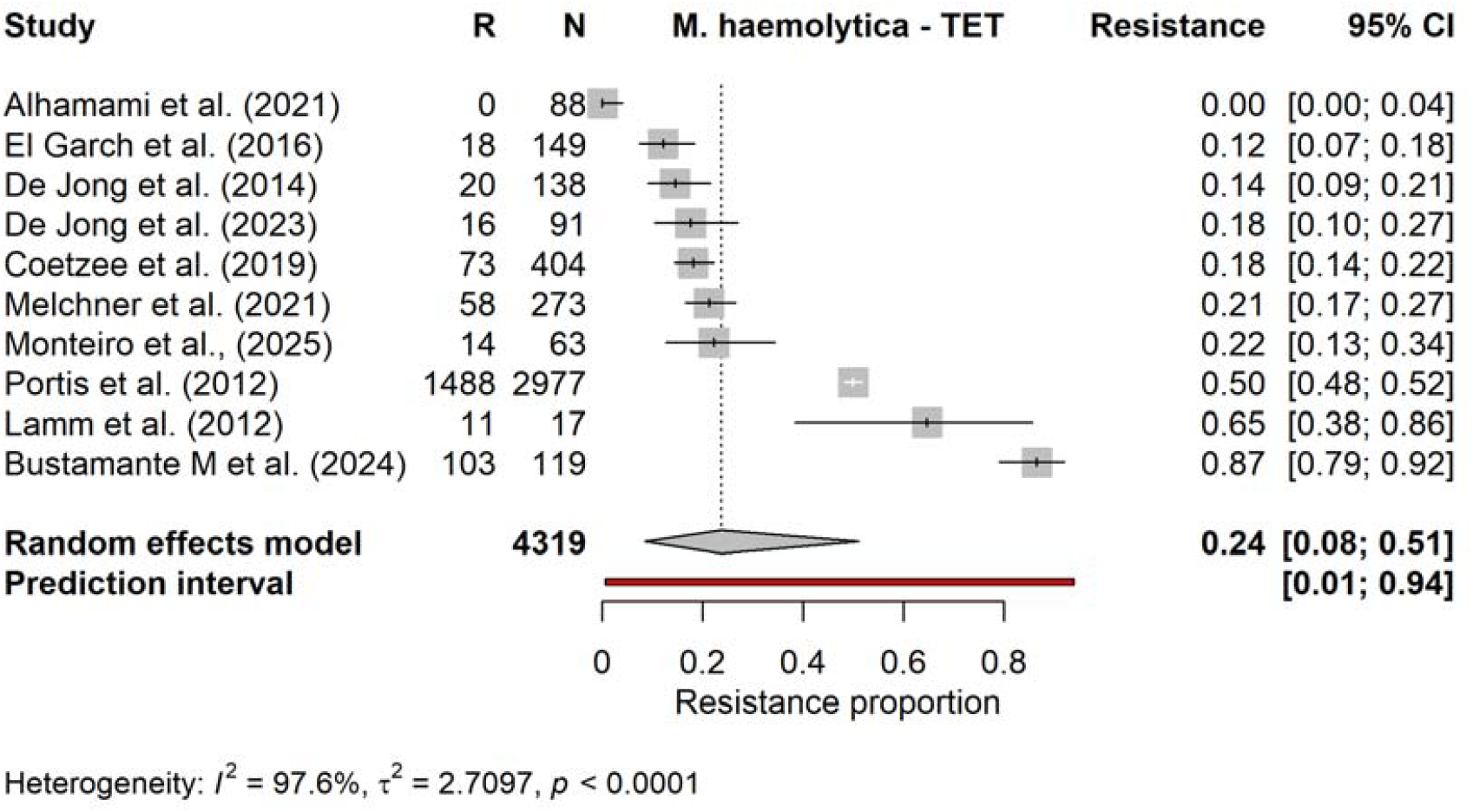
Forest plot of the pooled proportion of resistant isolates to tetracyclines (TET) in Mannheimia haemolytica (*M. haemolytica*).

**Figure 6.**
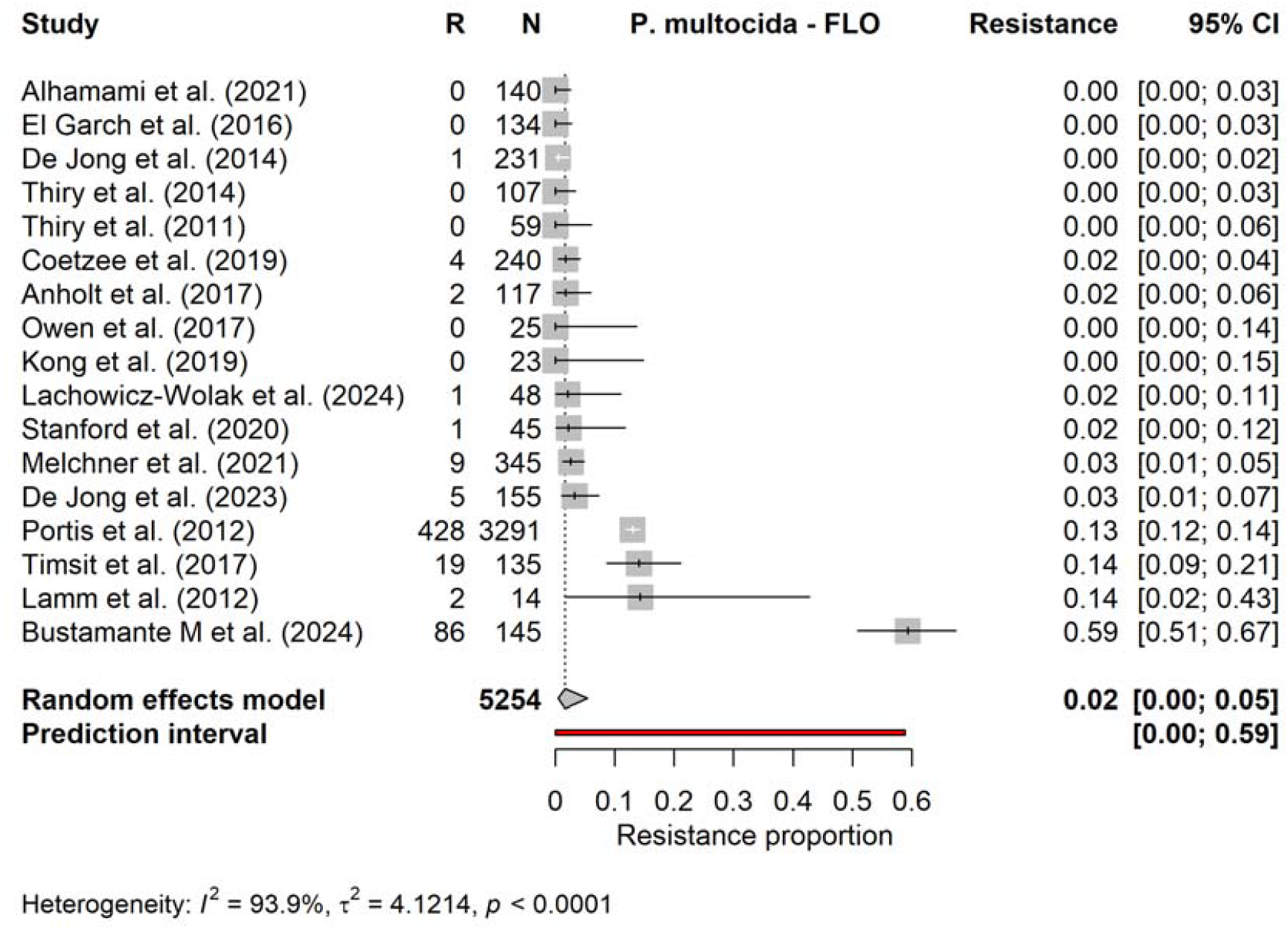
Forest plot of the pooled proportion of resistant isolates to florfenicol (FLO) in Pasteurella multocida (*P. multocida*).

### 3.5 Certainty of evidence for antimicrobial-specific resistance profiles

The pooled resistance estimates were additionally summarised by antimicrobial across the three BRD-associated pathogens to support clinical interpretation (Table 3). FLO and ENR showed the lowest pooled phenotypic resistance estimates across pathogens. FLO resistance was estimated at 0% in *H. somni*, 1% in *M. haemolytica* and 2% in *P. multocida*, with all CI remaining below the prespecified 10% resistance threshold. ENR resistance was similarly low, although the confidence interval for *H. somni* crossed the 10% threshold. PEN and TUL showed low pooled point estimates overall, but CI were wider and crossed multiple prespecified resistance thresholds for several pathogen-specific analyses. TET showed the highest pooled resistance estimates across all three pathogens, ranging from 12% in *H. somni* to 30% in *P. multocida*, although CI were wide and crossed multiple prespecified thresholds.

**Table 3.**
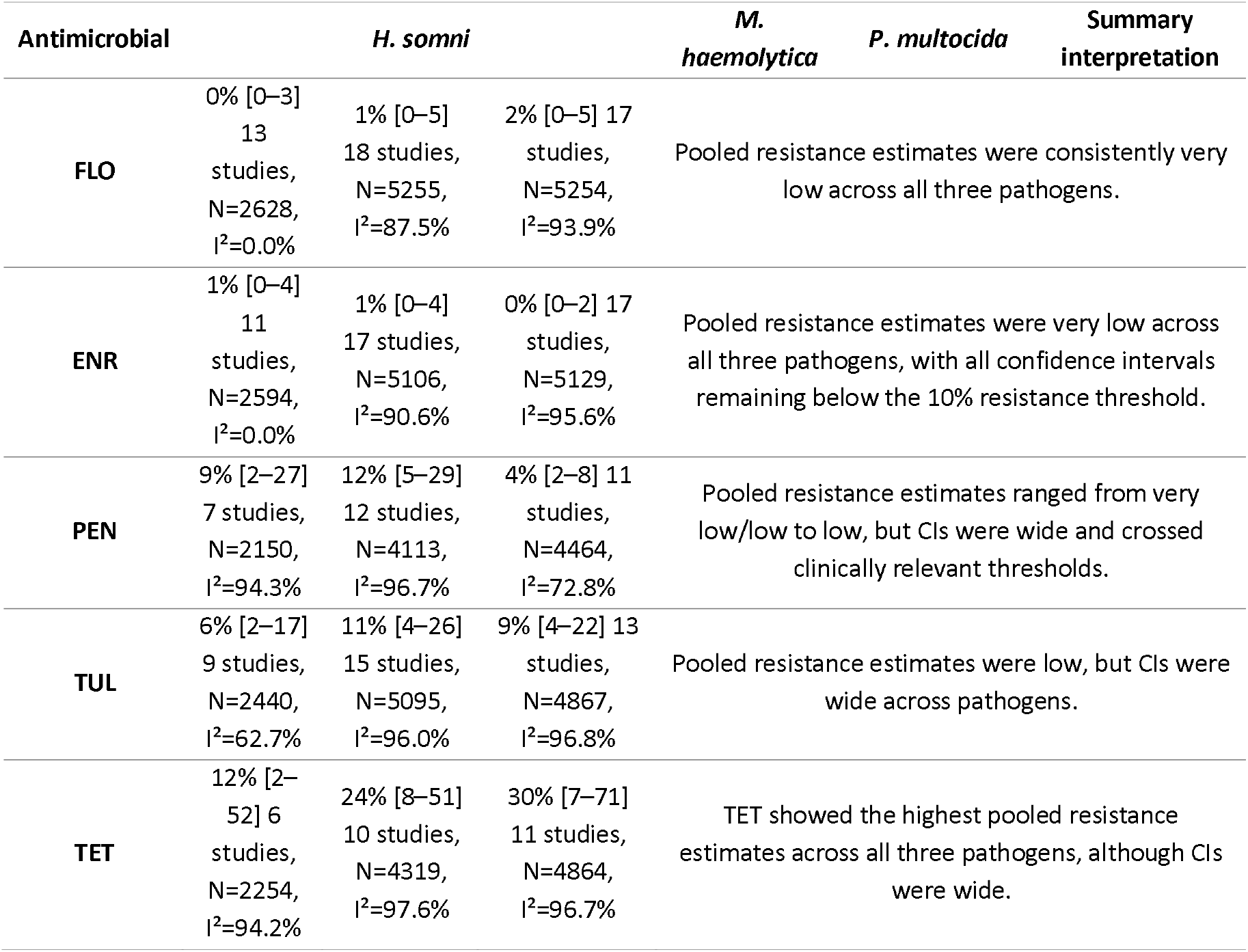
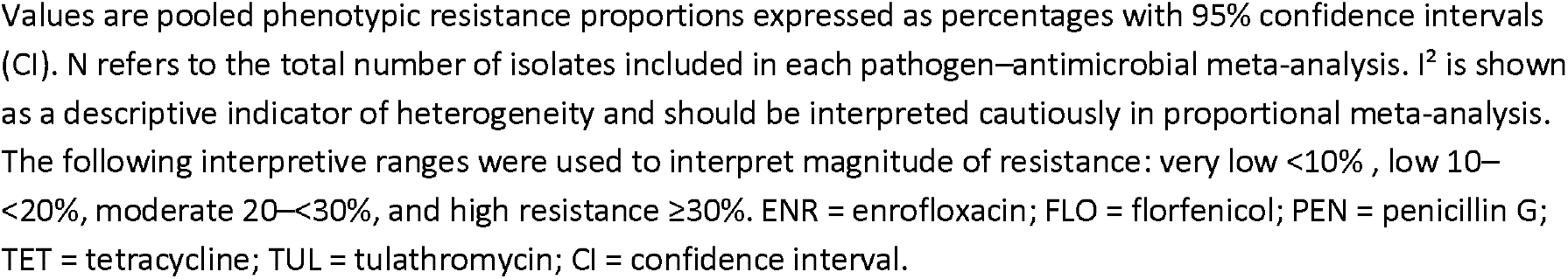
Drug-centred pooled resistance summary of antimicrobials across the three BRD-associated pathogens to support clinical interpretation

The GRADE-informed certainty assessment is summarised in Supplementary Materials II. The pathogen-specific pooled estimates, confidence intervals, I^2^ values, ranges of study-specific resistance estimates and threshold-crossing information supporting these judgements are provided in Supplementary Materials II. Certainty was downgraded for all antimicrobials because of serious concerns regarding risk of bias, mainly related to potential selection bias in isolate inclusion and limited representativeness of the sampled populations. For example, many included studies relied on diagnostic laboratory submissions, surveillance collections or convenience samples, where the underlying source population was not always clearly defined and selected isolates may not fully represent the wider population of cattle with clinical BRD.

Certainty for FLO and ENR was judged low because of serious concerns regarding risk of bias and inconsistency. Although pooled resistance estimates were low across the three pathogens, some pathogen-specific analyses included high-resistance outlying studies, limiting confidence that the pooled estimates were consistent across settings. Certainty for PEN, TUL and TET was judged very low, primarily because study-specific estimates and/or pooled confidence intervals crossed multiple prespecified resistance thresholds. Indirectness was not downgraded because the included studies directly measured phenotypic resistance in clinical isolates of the target pathogens using MIC data interpreted with CLSI clinical breakpoints. However, certainty ratings refer to the in vitro resistance profile only and should not be interpreted as certainty in clinical treatment efficacy. Publication bias was considered to present some concerns but was not downgraded. Although funnel plot asymmetry was reported for some pathogen–antimicrobial combinations, the direction of potential bias in proportional AMR studies is unclear, and both high and low resistance estimates may be publishable. Moreover, funnel plot methods and Egger’s tests are difficult to interpret in proportional meta-analysis and may reflect heterogeneity, sparse events or sampling differences rather than selective dissemination.

## 4. Discussion

Our series of meta-analyses revealed significant variations in resistance to commonly used antimicrobial drugs across different pathogens, as well as differences according to sampling period and geography. Although the pooled means provide a summary of the available literature, the wide confidence and prediction intervals indicate that resistance frequencies vary markedly between settings. Consequently, these averages should not be interpreted as expected resistance frequencies for an individual country, production system, farm, or clinical population.

Consistent with a previous review reporting a decline in susceptibility to antimicrobial agents commonly used for the treatment of BRD ^14^, we observed a significant increase in TUL resistance in *P. multocida* from 2013 (from 3.1% to 17%, *p* = 0.003). Because the two periods comprised different studies, countries, production systems and sampling frameworks, this finding should be regarded as a temporal subgroup association rather than evidence of a longitudinal increase in resistance. A comparable but non-significant increase in TUL resistance was also observed for *M. haemolytica* (2.6% to 17.5%). In contrast, FLO resistance in *H. somni* was significantly lower after 2013 than before (10% to 2.8%), but this comparison drew on a small number of contributing studies in each period and be interpreted cautiously. These differences do not necessarily establish a common temporal trend across the *Pasteurellaceae*. Given the similarity in resistance mechanisms and acquisition pathways, a shared trend cannot be excluded. It has been hypothesized that the pharmacokinetics of long-acting macrolides, particularly their prolonged persistence in plasma and tissues, may increase the duration during which bacterial populations are exposed to antimicrobial concentrations within the mutant selection window (MSW) ^52^, thereby promoting the selection of resistant subpopulations. TUL was introduced as a long-acting formulation in 2005 ^53^. This represents one possible biological explanation, but antimicrobial use and pharmacokinetic data were not analysed in the present study, and the hypothesis could therefore not be tested. Concerns regarding extended exposure to subinhibitory or intermediate concentrations have been raised, particularly for long-acting injectable formulations used in food animals ^54^. Similarly, the association between the use of long-acting macrolides and macrolide resistance in the human pathogen *Streptococcus pneumoniae* has been attributed to the extended selective window of these antimicrobials ^55^. These findings support prudent use of long-acting macrolides in veterinary practice, but do not establish a causal relationship between their use and the observed temporal subgroup difference.

PEN is among the oldest antimicrobial classes used in veterinary medicine, yet resistance to PEN and related β-lactams has generally remained relatively low in several major bacterial pathogens associated with BRD compared with resistance to other antimicrobial classes ^56^. The use of PEN for the treatment of BRD has remained limited in many production systems, largely because of the widespread availability and marketing of newer antimicrobial agents perceived to offer superior efficacy, broader antimicrobial spectra, longer durations of activity, and more convenient dosing regimens (e.g. no oral dosing of penicillin available). Economic factors and the labour-saving benefits of single-dose long-acting formulations have likely played a significant role in driving this shift in prescribing practices ^57^. However, increasing regional variation in antimicrobial susceptibility, along with the emergence of resistant isolates, has been reported with growing frequency in recent years. In Scandinavia, PEN remains a first choice for treatment for BRD and has long been recommended in local antimicrobial treatment guidelines ^58^. This approach is supported by the limited occurrence of *Mycoplasma bovis* in the region ^59, 60^, as this pathogen is intrinsically resistant to PEN owing to its lack of a cell wall. Consistent with this regional treatment strategy, a randomised clinical trial conducted in Sweden found no significant difference in the efficacy of PEN compared with TET or FLO ^61^. In contrast, both FLO and TET are more widely used both in the rest of Europe and in America ^62, 63^. Together with the relatively low resistance proportions observed in some pathogen-region combinations, these findings support consideration of PEN as a sustainable treatment option in appropriate contexts, particularly where local pathogen epidemiology and resistance patterns are favourable.

Resistance to FLO in *Pasteurellaceae* was infrequently detected in most included studies, whereas a substantially higher level of resistance was observed for TET. Possible explanations include differences in the historical intensity and duration of antimicrobial use, administration practices and co-selection of mobile resistance determinants. Oral administration of TET via medicated feed can lead to inconsistent dosing as well as underdosing, potentially fostering the development of AMR ^64^. Controlled experiments in chickens and mice have shown that administration route can influence AMR selection within the gut microbiota ^65^.

With respect to geographical differences, the continent-based subgroup analysis revealed significantly higher point estimates in North America than in Europe for TET resistance in *M. haemolytica* (48.1% vs. 16.6%), TET resistance in *P. multocida* (73.6% vs. 13.7%), and FLO resistance in *P. multocida* (6.3% vs. 0.9%). Higher point estimates in North America were also seen for several other combinations (e.g., FLO and TUL in *M. haemolytica*, and TUL in *P. multocida*), although these differences were not statistically significant. The observed differences may reflect several correlated factors, including antimicrobial use practices, production systems, previous antimicrobial treatment, case severity and the source of diagnostic isolates. Reported overall consumption of antimicrobials in livestock in North America (170.8 mg active ingredient/kg of livestock) in 2020 was estimated to be approximately 86% higher than in Europe ^66^. Cattle-specific antimicrobial consumption data from the same year also indicate a significantly higher usage in the US (161.3 mg/kg of livestock) compared to Denmark (26.5 mg/Population Correction Unit (PCU)) and France (38.3 mg/PCU) ^33, 66^. it is important to note that quantification methods between EUUSA cannot be compared as the calculation of PCU and kg of livestock biomass is likely to vary between continents. When comparing antimicrobial consumption across countries, it is also important to account for differences in production systems.

Most beef cattle in the US enter specialized feedlot production systems for the finishing phase, where feeder calves originating from multiple cow-calf and stocker operations are collected and intensively fattened, typically on high-energy grain-based diets, before slaughter ^67^. This production system carries a high risk of BRD because of the circulation of major BRD pathogens and the substantial stress experienced by animals shortly after arrival. This high disease pressure has contributed to the use of feedlot prophylactic and subsequently metaphylactic antimicrobial therapy to alleviate disease and maintain production. However, the feedlot industry is not the only cattle production system reliant on purchasing calves from multiple sources. The veal calf industry and more recently the dairy beef industry, both widely prevalent in Europe, may exhibit similar levels of commingling and antimicrobial usage. Differences between these production systems may contribute to geographical variation in antimicrobial exposure and resistance, but their effects could not be separated from those of study design, sampling strategy and other regional factors in the present analysis.

Certain potential biases in our meta-analyses should be acknowledged, particularly those related to the representativeness of isolates included in the individual studies. Many studies relied on routine diagnostic submissions, postmortem material, outbreak investigations or other selected clinical populations. Such samples may disproportionately represent animals with severe disease, previous antimicrobial exposure, treatment failure or mortality and may therefore contain higher resistance proportions than isolates from the broader population of cattle with BRD. The pooled estimates consequently describe resistance among isolates represented in the available clinical literature rather than population-level resistance proportion among all BRD cases. The studies also covered highly diverse cattle populations, production systems, geographical regions, and sampling frameworks, ranging from large multicentre investigations to studies involving only a limited number of farms. Consequently, the clinical isolates analysed in some studies may not fully reflect the wider BRD pathogen population within the respective regions or production systems, thereby limiting the external validity and generalisability of the resulting AMR estimates.

Differences in sampling strategies between studies may also have influenced our ability to pool results. For example, studies varied with respect to case definitions, diagnostic approaches, inclusion criteria, the number of farms sampled, and whether isolates originated from routine diagnostic submissions, outbreak investigations, or targeted surveillance programmes. Such differences may contribute to heterogeneity in the observed resistance estimates. Furthermore, it cannot be excluded that multiple isolates originating from a limited number of farms contributed disproportionately to the pooled estimates in some analyses. However, we were reluctant to introduce additional exclusion criteria because of the relatively limited number of eligible studies available for inclusion. In contrast, we anticipate only limited bias arising from laboratory procedures owing to the relatively strict eligibility criteria applied for antimicrobial susceptibility testing methods and interpretation of susceptibility data.

The GRADE-informed certainty assessment indicated that confidence in the exact antimicrobial-specific resistance profiles was limited. Although FLO and ENR showed low pooled proportions of phenotypic resistance across the three main BRD-associated pathogens, certainty was judged low because of concerns regarding selection bias/representativeness and inconsistency between studies. For PEN, TUL and TET, certainty was judged very low, reflecting wider CI and substantial between-study variation, with study-specific estimates and/or pooled CI crossing multiple prespecified resistance thresholds. These findings indicate that the pooled estimates should be interpreted as broad summaries of available in vitro resistance data rather than as precise estimates. Local epidemiology, production system, recent antimicrobial exposure and sampling framework may substantially influence the likelihood of encountering resistant BRD pathogens.

Importantly, phenotypic resistance in vitro should not be equated with clinical treatment failure, and low in vitro resistance should not be interpreted as direct evidence of superior clinical efficacy. Clinical outcome in BRD is influenced by multiple factors beyond MIC classification, including timing of treatment, disease severity and chronicity, extent of lung pathology, pathogen involvement, host immunity, pharmacokinetic/pharmacodynamic exposure at the site of infection, and management conditions. This distinction is supported by the companion ENOVAT systematic review of antimicrobial treatment efficacy for BRD, which did not identify clinically meaningful short-term differences between antimicrobial classes in reducing re-treatment, although certainty ranged from very low to moderate and re-treatment was a surrogate for short-term therapeutic failure ^68^. The frequently cited “90–60 rule” further illustrates the imperfect relationship between antimicrobial susceptibility and clinical outcome: infections caused by susceptible bacteria respond clinically to appropriate antimicrobial therapy in approximately 90% of cases, whereas infections caused by resistant bacteria may still respond in approximately 60% of cases ^69^. Thus, AMR is an important risk factor for reduced probability of clinical success, but resistant isolates do not translate directly into treatment failure, and low resistance proportions should not be interpreted as proof of clinical effectiveness.

The methodological quality appraisal tool provided a robust framework for evaluating the internal validity of the predominantly cross-sectional studies in the dataset of the present study (n=19). The assessment of potential publication bias provided additional context for interpreting the evidence base. However, it is important to note that the tool is tailored primarily for cross-sectional designs ^19^ and its application to the experimental and cohort studies in our dataset (n=6) represents a limitation in providing a uniform quality metric. In addition, because the AXIS tool does not generate a composite study-level risk-of-bias score, precluding a formal risk-of-bias-based subgroup or sensitivity analysis. Furthermore, satisfactory reporting according to the appraisal tool does not eliminate concerns regarding selection bias, representativeness, or the external validity of the resistance estimates.

Egger’s test indicated significant funnel-plot asymmetry for ENR resistance in *H. somni* and PEN resistance in *M. haemolytica*, which could reflect publication bias of small-study effects. However, because asymmetry tests have limitations in meta-analyses of proportions, particularly in the presence of substantial heterogeneity and zero-event studies, these findings should be interpreted cautiously and not as definitive evidence of publication bias ^25^. Stratifying the data based on production systems would have been valuable, but it was not feasible since most of the retrieved publications were passive surveillance studies analysing clinical isolates from multiple sources without providing information on the production system from which each isolate originated.

Our selection of search engines and inclusion criteria excluded national reports on AMR surveillance, which could have provided valuable information ^70^. However, to the authors’ knowledge, BRD pathogens are rarely included in such reports at present, and it is not uncommon for scientific articles to include isolates collected through surveillance programmes itself. Moreover, published studies likely capture only a fraction of available surveillance data, as additional information generated by diagnostic laboratories and national, regional, or local surveillance programmes may remain unpublished, inaccessible, or insufficiently standardized for inclusion in comparative studies. Finally, screening and extraction were not done by two independent reviewers for each paper, as is typically required for systematic reviews to minimize bias and ensure accuracy. While this could be considered a methodological limitation, it is important to note that a calibration exercise was conducted prior to the review process, during which 100% agreement was observed between reviewers of the presented study.

## Conclusions

This systematic review and series of meta-analyses provide critical insights into temporal and geographical variation of AMR in bacterial pathogens associated with BRD. Tetracycline consistently exhibited the highest pooled resistance proportions across all three bovine respiratory pathogens, while florfenicol and enrofloxacin showed comparatively low pooled proportions of resistance, although certainty in these antimicrobial-specific resistance profiles was low because of risk of bias and inconsistency. The observed geographical and temporal differences support the need for enhanced global surveillance and more judicious antimicrobial stewardship within the cattle industry.

The findings highlight the need for locally and regionally informed antimicrobial stewardship strategies contributing to the ENOVAT guidelines for the treatment of BRD and broader One Health efforts to preserve the effectiveness of medically important antimicrobials.

## Supporting information

Supplementary Materials_I, II, IV, VI, VII

Supplementary Materials_III, V

## Transparency declarations

None to declare.

## Author contributions

Qamer Mahmood (Data curation, Formal analysis, Investigation, Methodology, Validation, Visualization, Writing—original draft, Writing—review & editing), Philip Rasmussen (Formal analysis, Methodology, Visualization, Writing—review & editing), Peter Damborg (Conceptualization, Methodology, Writing—original draft, Writing—review & editing, Supervision), Kristine Berg Hansen (Data curation, Investigation, Methodology, Validation, Writing—original draft, Writing— review & editing), Camilla Bro Gregersen (Data curation, Investigation, Methodology, Validation, Writing—original draft, Writing—review & editing), Karolina Scahill (Methodology, Writing—review & editing), Luis Pedro Carmo (Methodology, Writing—review & editing), Clair L Firth (Methodology, Writing—review & editing), Bart Pardon (Methodology, Writing—review & editing), Maria Stokstad (Methodology, Writing—review & editing), Lise Marie Ånestad (Methodology, Writing—review & editing), Luca Guardabassi (Conceptualization, Methodology, Writing—original draft, Writing— review & editing, Supervision), Beate Conrady (Conceptualization, Methodology, Writing—review & editing, Supervision).

## Supplementary Materials

- Supplementary Materials I (PRISMA 2020 Checklist)
- Supplementary Materials II (Review protocol, quality appraisal results, GRADE-assessment)
- Supplementary Materials III (Data extraction tool)
- Supplementary Materials IV (Funnel plots and Egger’s test results)
- Supplementary Materials V (Study design sensitivity analysis results)
- Supplementary Materials VI (Pathogen-antimicrobial forest plots)
- Supplementary Materials VII (Continental and temporal subgroup forest plots)

