## Supplementary Materials_I, II, IV, VI, VII for "In vitro antimicrobial resistance in clinical isolates of bacterial pathogens causing bovine respiratory disease: a systematic review and meta-analysis"

#### PRISMA 2020 Checklist

| Section and Topic | Item # | Checklist item | Location where item is reported in main manuscript (Line Numbers) |
|---|---|---|---|
| **TITLE** | | | |
| Title | 1 | Identify the report as a systematic review. | 1-2 |
| **ABSTRACT** | | | |
| Abstract | 2 | See the PRISMA 2020 for Abstracts checklist. | 19-47 |
| **INTRODUCTION** | | | |
| Rationale | 3 | Describe the rationale for the review in the context of existing knowledge. | 69-78 |
| Objectives | 4 | Provide an explicit statement of the objective(s) or question(s) the review addresses. | 79-83 |
| **METHODS** | | | |
| Eligibility criteria | 5 | Specify the inclusion and exclusion criteria for the review and how studies were grouped for the syntheses. | 99-109 |
| Information sources | 6 | Specify all databases, registers, websites, organisations, reference lists and other sources searched or consulted to identify studies. Specify the date when each source was last searched or consulted. | 88-98 |
| Search strategy | 7 | Present the full search strategies for all databases, registers and websites, including any filters and limits used. | 97-98 |
| Selection process | 8 | Specify the methods used to decide whether a study met the inclusion criteria of the review, including how many reviewers screened each record and each report retrieved, whether they worked independently, and if applicable, details of automation tools used in the process. | 99-109 |
| Data collection process | 9 | Specify the methods used to collect data from reports, including how many reviewers collected data from each report, whether they worked independently, any processes for obtaining or confirming data from study investigators, and if applicable, details of automation tools used in the process. | 99-109 |
| Data items | 10a | List and define all outcomes for which data were sought. Specify whether all results that were compatible with each outcome domain in each study were sought (e.g. for all measures, time points, analyses), and if not, the methods used to decide which results to collect. | 110-123 |
| | 10b | List and define all other variables for which data were sought (e.g. participant and intervention characteristics, funding sources). Describe any assumptions made about any missing or unclear information. | 110-123 |
| Study risk of bias assessment | 11 | Specify the methods used to assess risk of bias in the included studies, including details of the tool(s) used, how many reviewers assessed each study and whether they worked independently, and if applicable, details of automation tools used in the process. | 124-133 |
| Effect measures | 12 | Specify for each outcome the effect measure(s) (e.g. risk ratio, mean difference) used in the synthesis or presentation of results. | 134-162 |
| Synthesis methods | 13a | Describe the processes used to decide which studies were eligible for each synthesis (e.g. tabulating the study intervention characteristics and comparing against the planned groups for each synthesis (item #5)). | 134-162 |
| | 13b | Describe any methods required to prepare the data for presentation or synthesis, such as handling of missing summary statistics, or data conversions. | 134-162 |
| | 13c | Describe any methods used to tabulate or visually display results of individual studies and syntheses. | 134-162 |
| | 13d | Describe any methods used to synthesize results and provide a rationale for the choice(s). If meta-analysis was performed, describe the model(s), method(s) to identify the presence and extent of statistical heterogeneity, and software package(s) used. | 134-162 |
| | 13e | Describe any methods used to explore possible causes of heterogeneity among study results (e.g. subgroup analysis, meta-regression). | 134-162 |
| | 13f | Describe any sensitivity analyses conducted to assess robustness of the synthesized results. | 134-162 |

### PRISMA 2020 Checklist

| Section and Topic | Item # | Checklist item | Location where item is reported in main manuscript (Line Numbers) |
|---|---|---|---|
| Reporting bias assessment | 14 | Describe any methods used to assess risk of bias due to missing results in a synthesis (arising from reporting biases). | 124-133 |
| Certainty assessment | 15 | Describe any methods used to assess certainty (or confidence) in the body of evidence for an outcome. | 134-162 |
| **RESULTS** | | | |
| Study selection | 16a | Describe the results of the search and selection process, from the number of records identified in the search to the number of studies included in the review, ideally using a flow diagram. | 195-205 |
| | 16b | Cite studies that might appear to meet the inclusion criteria, but which were excluded, and explain why they were excluded. | 195-205 |
| Study characteristics | 17 | Cite each included study and present its characteristics. | 206-226 |
| Risk of bias in studies | 18 | Present assessments of risk of bias for each included study. | 227-259 |
| Results of individual studies | 19 | For all outcomes, present, for each study: (a) summary statistics for each group (where appropriate) and (b) an effect estimate and its precision (e.g. confidence/credible interval), ideally using structured tables or plots. | 260-306 |
| Results of syntheses | 20a | For each synthesis, briefly summarise the characteristics and risk of bias among contributing studies. | 227-259 |
| | 20b | Present results of all statistical syntheses conducted. If meta-analysis was done, present for each the summary estimate and its precision (e.g. confidence/credible interval) and measures of statistical heterogeneity. If comparing groups, describe the direction of the effect. | 260-306 |
| | 20c | Present results of all investigations of possible causes of heterogeneity among study results. | 260-306 |
| | 20d | Present results of all sensitivity analyses conducted to assess the robustness of the synthesized results. | 260-306 |
| Reporting biases | 21 | Present assessments of risk of bias due to missing results (arising from reporting biases) for each synthesis assessed. | 227-259 |
| Certainty of evidence | 22 | Present assessments of certainty (or confidence) in the body of evidence for each outcome assessed. | 299-306 |
| **DISCUSSION** | | | |
| Discussion | 23a | Provide a general interpretation of the results in the context of other evidence. | 269-274 |
| | 23b | Discuss any limitations of the evidence included in the review. | 451-474<br>500-516 |
| | 23c | Discuss any limitations of the review processes used. | 517-528 |
| | 23d | Discuss implications of the results for practice, policy, and future research. | 375-438 |
| **OTHER INFORMATION** | | | |
| Registration and protocol | 24a | Provide registration information for the review, including register name and registration number, or state that the review was not registered. | 86-87 |
| | 24b | Indicate where the review protocol can be accessed, or state that a protocol was not prepared. | 86-87 |
| | 24c | Describe and explain any amendments to information provided at registration or in the protocol. | NA |
| Support | 25 | Describe sources of financial or non-financial support for the review, and the role of the funders or sponsors in the review. | 541-542 |

### PRISMA 2020 Checklist

| Section and Topic | Item # | Checklist item | Location where item is reported in main manuscript (Line Numbers) |
|---|---|---|---|
| Competing interests | 26 | Declare any competing interests of review authors. | 544 |
| Availability of data, code and other materials | 27 | Report which of the following are publicly available and where they can be found: template data collection forms; data extracted from included studies; data used for all analyses; analytic code; any other materials used in the review. | 546 |

*From:* Page MJ, McKenzie JE, Bossuyt PM, Boutron I, Hoffmann TC, Mulrow CD, et al. The PRISMA 2020 statement: an updated guideline for reporting systematic reviews. BMJ 2021;372:n71. doi: 10.1136/bmj.n71.
This work is licensed under CC BY 4.0. To view a copy of this license, visit https://creativecommons.org/licenses/by/4.0/

### Supplementary Material II

#### Protocol for a systematic review and meta-analysis

**Title:** In vitro antimicrobial resistance in clinical isolates of bacterial pathogens causing bovine respiratory disease: a systematic review and meta-analysis

**1. Objectives**

The objective of this systematic review and meta-analysis was to estimate the pooled proportions of AMR in BRD pathogens *M. haemolytica*, *P. multocida*, and *H. somni* from cattle (dairy and beef). Furthermore, the review aimed to investigate geographical and temporal variation in resistance to five antimicrobial agents commonly used for the treatment of BRD: enrofloxacin (ENR), florfenicol (FLO), penicillin G (PEN), tetracyclines (TET), and tulathromycin (TUL).

**2. PEO elements:**

**Population (P):** Clinical isolates of *Mannheimia haemolytica*, *Pasteurella multocida*, and *Histophilus somni* obtained from cattle with clinical respiratory disease.

**Exposure (E):** In vitro antimicrobial susceptibility testing based on minimum inhibitory concentration (MIC) determination and interpreted using veterinary Clinical and Laboratory Standards Institute (CLSI) clinical breakpoints.

**Outcome (O):** The proportion of phenotypic antimicrobial resistance to enrofloxacin, florfenicol, penicillin G, tetracyclines, and tulathromycin, including temporal and geographical variation in resistance frequencies.

**3. Inclusion criteria**

Inclusion criteria were comprised of original research articles (i) reporting bacterial isolates obtained from clinical respiratory disease cases in cattle, (ii) providing minimum inhibitory concentration data interpreted using Clinical and Laboratory Standards Institute (CLSI) clinical breakpoints, and (iii) published in English.

| Exclusion criteria | No. studies excluded |
|---|---|
| Studies investigating non-clinical isolates where MIC data from clinical isolates could not be distinguished | 36 |
| Studies including isolates from host species other than cattle which could not be discerned from cattle isolates | 11 |
| Studies including less than 10 isolates per bacterial species | 16 |
| Studies assessing antimicrobial resistance genotypically | 15 |
| Studies where all isolates were derived from the same farm | 10 |
| Studies where animals had been sampled more than once if it exceeded 10% of total amount of samples per bacterial species | 9 |

| | |
|---|---|
| Studies affected by selection bias (e.g., studies focussing on certain serotypes) | 3 |
| Not original publication (e.g. conference abstracts or reviews or online reports) | 7 |
| Studies where >20% of the isolates originated from non-respiratory samples and they could not be separated from respiratory isolates | 1 |
| Studies not providing data on AMR | 9 |
| Studies reporting $MIC_{50}$ values without the possibility to determine the percentage resistant (R) isolates | 5 |
| Studies showing the results by histograms without providing the actual data | 2 |
| Studies testing other bacterial species | 2 |
| Studies with high risk of data-duplication | 2 |
| Studies where intermediate (I) isolates were reported as resistant (R) isolates, and the percentage of R isolates could not be determined | 4 |
| Studies using the disc-diffusion method | 5 |
| Studies using the agar dilution method | 5 |
| Studies using gradient-strip methods | 1 |
| Studies using ECOFFs instead of breakpoints | 1 |
| Studies using breakpoints for bacteria isolated from humans | 1 |
| Studies using breakpoints for other bacterial species | 6 |
| Studies using unknown breakpoints | 2 |
| Studies not reporting data for the antimicrobials selected for the meta-analysis | 2 |
| **Total excluded** | **155** |

#### 5. Literature search

Following the inclusion criteria, a literature search will be conducted in PubMed and Embase. The search strategy will involve a multi-strand approach that uses a series of searches, with different combinations of concepts to gather all possibly related research and thus achieve high sensitivity (Cochrane Handbook for Reviews, 2019). The list of published articles identified from both databases will be merged into a single file using Endnote and de-duplicated. The resulting list will be imported to Rayyan tool to proceed with the two phases screening process. A minimum of two reviewers should be able to perform the literature search.

**List of databases to be searched:**

| Database | Interface | URL |
|---|---|---|
| PubMed | Ovid MEDLINE | https://ovidsp.dc1.ovid.com/ovid-new-b/ovidweb.cgi |
| Elsevier | Embase | https://www.embase.com/search/ |

The concept of the search strategy will be the following:

*[Antimicrobials]* AND *[Bacterial species]* AND *[Host]*

Time period specified across databases: January 1, 2010, to 16 April 2026 (16 years)

The specific search terms and strategy to identify references will be the following:

| Database | Search terms |
| --- | --- |
| PubMed | ((("antibiotic"[Title/Abstract] OR "antibiotics"[Title/Abstract] OR "antimicrobial"[Title/Abstract] OR "antimicrobials"[Title/Abstract] OR "Anti-Bacterial Agents"[MeSH Terms:noexp]) AND ("resistan*"[Title/Abstract] OR "susceptib*"[Title/Abstract])) OR ("Microbial Sensitivity Tests"[MeSH Terms] OR "drug resistance, microbial"[MeSH Terms])) AND (("Cattle"[Title/Abstract] OR "cow"[Title/Abstract] OR "cows"[Title/Abstract] OR "bull"[Title/Abstract] OR "bulls"[Title/Abstract] OR "calf"[Title/Abstract] OR "calves"[Title/Abstract] OR "bovine"[Title/Abstract] OR "Cattle"[MeSH Terms])) AND ("Haemophilus somnus"[MeSH Terms] OR "Mannheimia haemolytica"[MeSH Terms] OR "Pasteurella multocida"[MeSH Terms] OR "Haemophilus somnus"[Title/Abstract] OR "Mannheimia haemolytica"[Title/Abstract] OR "Pasteurella multocida"[Title/Abstract]) |
| Embase | **“Antimicrobials”**<br><br>1. antibiotic resistance/ or exp antibiotic sensitivity/ or exp drug resistance/<br>2. susceptib*.mp. [mp=title, abstract, heading word, drug trade name, original title, device manufacturer, drug manufacturer, device trade name, keyword, floating subheading word, candidate term word]<br>3. resistan*.mp. [mp=title, abstract, heading word, drug trade name, original title, device manufacturer, drug manufacturer, device trade name, keyword, floating subheading word, candidate term word]<br>4. 2 or 3<br>5. antibiotic.mp. [mp=title, abstract, heading word, drug trade name, original title, device manufacturer, drug manufacturer, device trade name, keyword, floating subheading word, candidate term word]<br>6. antibiotics.mp. [mp=title, abstract, heading word, drug trade name, original title, device manufacturer, drug manufacturer, device trade name, keyword, floating subheading word, candidate term word]<br>7. antimicrobial.mp. [mp=title, abstract, heading word, drug trade name, original title, device manufacturer, drug manufacturer, device trade name, keyword, floating subheading word, candidate term word]<br>8. antimicrobials.mp. [mp=title, abstract, heading word, drug trade name, original title, device manufacturer, drug manufacturer, device trade name, keyword, floating subheading word, candidate term word]<br>9. 5 or 6 or 7 or 8<br>10. antibiotic agent/<br>11. 10 or 9<br>12. 11 and 4<br>13. 12 or 1 |

| | |
|---|---|
| | **“Host”**<br><br>1. bovine/<br>2. "calf (bovine)"/<br>3. (Cattle or cow or cows or bull or bulls or calf or calves or bovine).mp. [mp=title, abstract, heading word, drug trade name, original title, device manufacturer, drug manufacturer, device trade name, keyword, floating subheading word, candidate term word]<br>4. 1 or 2 or 3<br><br>**“Bacterial species”**<br><br>1. Histophilus somni/<br>2. Mannheimia haemolytica/<br>3. Pasteurella multocida/<br>4. 1 or 2 or 3<br>5. (“Haemophilus somnus” or “Mannheimia haemolytica” or “Pasteurella multocida”).mp. [mp=title, abstract, heading word, drug trade name, original title, device manufacturer, drug manufacturer, device trade name, keyword, floating subheading word, candidate term word]<br>6. 4 or 5 |

### Quality appraisal of included studies

**Table S1.** Quality appraisal of included studies using AXIS checklist (tool developed by: Downes et al., 2016).

| Sections | Sr. No. | Question | S 1 | S 2 | S 3 | S 4 | S 5 | S 6 | S 7 | S 8 | S 9 | S 10 | S 11 | S 12 | S 13 | S 14 | S 15 | S 16 | S 17 | S 18 | S 19 | S 20 | S 21 | S 22 | S 23 | S 24 |
|---|---|---|---|---|---|---|---|---|---|---|---|---|---|---|---|---|---|---|---|---|---|---|---|---|---|---|
| **Introduction** | 1 | Were the aims/objectives of the study clear? | Y | Y | Y | Y | Y | Y | Y | Y | Y | Y | Y | Y | Y | Y | Y | Y | Y | Y | Y | Y | Y | Y | Y | Y |
| **Methods** | 2 | Was the study design appropriate for the stated aim(s)? | Y | Y | Y | Y | Y | Y | Y | Y | Y | Y | Y | Y | Y | Y | Y | Y | Y | Y | Y | Y | Y | Y | Y | Y |
| | 3 | Was the sample size justified? | Y | N | N | N | Y | N | N | N | Y | N | Y | N | N | N | N | N | N | Y | Y | N | N | N | N | N |
| | 4 | Was the target/reference population clearly defined? (Is it clear who the research was about?) | Y | Y | Y | Y | Y | Y | Y | Y | Y | Y | Y | Y | Y | Y | Y | Y | Y | Y | Y | Y | Y | Y | Y | Y |
| | 5 | Was the sample frame taken from an appropriate population base so that it closely represented the target/reference population under investigation? | Y | Y | Y | Y | Y | Y | Y | Y | Y | Y | Y | Y | Y | Y | DK | Y | Y | Y | Y | Y | Y | Y | Y | Y |
| | 6 | Was the selection process likely to select subjects/participants that were representative of the target/reference population under investigation? | N | N | N | N | Y | N | N | N | Y | N | Y | N | N | N | N | N | N | Y | Y | N | N | N | Y | N |
| | 7 | Were measures undertaken to address and categorise non-responders? | DK | N | N | N | Y | N | N | N | DK | DK | Y | N | N | N | N | N | N | N | N | N | N | N | N | N |
| | 8 | Were the risk factor and outcome variables | Y | Y | Y | Y | Y | Y | Y | Y | Y | Y | Y | Y | Y | Y | Y | Y | Y | Y | Y | Y | Y | Y | Y | Y |

| | | | | | | | | | | | | | | | | | | | | | | | | | | |
|---|---|---|---|---|---|---|---|---|---|---|---|---|---|---|---|---|---|---|---|---|---|---|---|---|---|---|
| | | measured appropriate to the aims of the study? | | | | | | | | | | | | | | | | | | | | | | | | |
| | 9 | Were the risk factor and outcome variables measured correctly using instruments/measurements that had been trialed, piloted or published previously? | Y | Y | Y | Y | Y | Y | Y | Y | Y | Y | Y | Y | Y | Y | Y | Y | Y | Y | Y | Y | Y | Y | Y | Y |
| | 10 | Is it clear what was used to determined statistical significance and/or precision estimates? (eg, p values, CIs) | Y | Y | N | Y | Y | Y | Y | Y | Y | Y | Y | N | Y | Y | Y | Y | Y | Y | Y | N | Y | Y | Y | Y |
| | 11 | Were the methods (including statistical methods) sufficiently described to enable them to be repeated? | Y | Y | Y | Y | Y | Y | Y | Y | Y | Y | Y | Y | Y | Y | Y | Y | Y | Y | Y | Y | Y | Y | Y | Y |
| **Results** | 12 | Were the basic data adequately described? | Y | Y | Y | Y | Y | Y | Y | Y | Y | Y | Y | Y | Y | Y | Y | Y | Y | Y | Y | Y | Y | Y | Y | Y |
| | 13 | Does the response rate raise concerns about non-response bias? | DK | DK | N | N | N | DK | DK | DK | DK | DK | N | DK | DK | DK | DK | DK | DK | N | N | DK | N | DK | N | DK |
| | 14 | If appropriate, was information about non-responders described? | N | N | N | N | Y | N | N | N | N | N | Y | N | N | N | N | N | N | N | N | N | N | N | N | N |
| | 15 | Were the results internally consistent? | Y | Y | Y | Y | Y | Y | Y | Y | Y | Y | Y | Y | Y | Y | Y | Y | Y | Y | Y | Y | Y | Y | Y | Y |
| | 16 | Were the results for the analyses described in the methods, presented? | Y | Y | Y | Y | Y | Y | Y | Y | Y | Y | Y | Y | Y | Y | Y | Y | Y | Y | Y | Y | Y | Y | Y | Y |
| **Discussion** | 17 | Were the authors' discussions and conclusions justified by the results? | Y | Y | Y | Y | Y | Y | Y | Y | Y | Y | Y | Y | Y | Y | Y | Y | Y | Y | Y | Y | Y | Y | Y | Y |

| | | | | | | | | | | | | | | | | | | | | | | | | | | |
|---|---|---|---|---|---|---|---|---|---|---|---|---|---|---|---|---|---|---|---|---|---|---|---|---|---|---|
| | 18 | Were the limitations of the study discussed? | Y | Y | N | Y | Y | Y | Y | Y | Y | N | Y | Y | Y | Y | Y | Y | Y | Y | Y | Y | Y | Y | Y | Y |
| **Other** | 19 | Were there any funding sources or conflicts of interest that may affect the authors' interpretation of the results? | Y | Y | Y | Y | Y | Y | Y | N | Y | Y | Y | Y | Y | Y | Y | Y | Y | Y | Y | Y | Y | Y | Y | Y |
| | 20 | Was ethical approval or consent of participants attained? | Y | N | N | N | Y | Y | N | N | Y | N | Y | Y | N | N | N | N | N | Y | Y | N | Y | N | Y | Y |

Y = Yes, N = No, DK = Do not know/ not given, S = Study

Study 1: Monteiro et al., 2025, Study 2: Coetzee et al., 2019, Study 3: Blondeau et al., 2012, Study 4: Ueno et al., 2022, Study 5: Morgan Bustamante et al., 2024, Study 6: Lachowicz-Wolak et al., 2025, Study 7: Alhamami et al., 2021, Study 8: Melchner et al., 2021, Study 9: Anholt et al., 2017, Study 10: Stanford et al., 2020, Study 11: Timsit et al., 2017, Study 12: Kong et al., 2019, Study 13: Portis et al., 2012, Study 14: Lamm et al., 2012, Study 15: El Garch et al., 2016, Study 16: de Jong et al., 2014, Study 17: de Jong et al., 2023, Study 18: Thiry et al., 2011, Study 19: Thiry et al., 2014, Study 20: Kong et al., 2014, Study 21: Owen et al., 2017, Study 22: Andrés-Lasheras et al., 2019, Study 23: Rérat et al., 2012, Study 24: Dutta et al., 2021

#### Certainty assessment by antimicrobial

| Antimicrobial | Risk of bias | Inconsistency | Imprecision | Indirectness | Publication bias | Overall certainty |
|---|---|---|---|---|---|---|
| **FLO** | Downgraded 1 level for serious concerns regarding potential selection bias and limited representativeness of isolate selection. | Downgraded 1 level for serious inconsistency; estimates for *M. haemolytica* and *P. multocida* included high-resistance outlying studies despite low pooled estimates. | Not downgraded; all pooled CIs remained below the 10% threshold. | Not downgraded; evidence directly addressed the in vitro resistance question. | Not downgraded[1] | **Low** |
| **ENR** | Downgraded 1 level for serious concerns regarding potential selection bias and limited representativeness of isolate selection. | Downgraded 1 level for serious inconsistency; high-resistance outliers/clusters were present in some pathogen-specific analyses. | Some concerns, not downgraded[2] | Not downgraded; evidence directly addressed the in vitro resistance question. | Not downgraded[1] | **Low** |
| **PEN** | Downgraded 1 level for serious concerns regarding potential | Downgraded 2 levels for very serious inconsistency; study | Downgraded 2 levels for very serious imprecision; pooled | Not downgraded; evidence directly | Not downgraded[1] | **Very low** |

| | | | | | | |
|---|---|---|---|---|---|---|
| | selection bias and limited representativeness of isolate selection. | estimates spanned multiple thresholds and high outliers were present. | CIs crossed multiple prespecified thresholds across pathogens. | addressed the in vitro resistance question. | | |
| **TUL** | Downgraded 1 level for serious concerns regarding potential selection bias and limited representativeness of isolate selection. | Downgraded 2 levels for very serious inconsistency; study estimates spanned multiple thresholds across pathogens. | Downgraded 2 levels for very serious imprecision; pooled CIs crossed multiple prespecified thresholds across pathogens. | Not downgraded; evidence directly addressed the in vitro resistance question. | Not downgraded[1] | **Very low** |
| **TET** | Downgraded 1 level for serious concerns regarding potential selection bias and limited representativeness of isolate selection. | Downgraded 2 levels for very serious inconsistency; study estimates spanned multiple thresholds across pathogens. | Downgraded 2 levels for very serious imprecision; pooled CIs crossed multiple prespecified thresholds across pathogens. | Not downgraded; evidence directly addressed the in vitro resistance question. | Not downgraded[1] | **Very low** |

Certainty was initially rated as high for the descriptive question of phenotypic resistance proportions and downgraded according to GRADE principles here adapted for proportional meta-analysis. Prespecified resistance thresholds were: <10% very low, 10–<20% low, 20–<30% moderate and ≥30% high resistance. Inconsistency was assessed using study-specific point estimates and whether they crossed threshold-defined ranges; imprecision was assessed using the number of prespecified thresholds crossed by the pooled 95% CI. Where wide random-effects confidence intervals appeared partly attributable to between-study heterogeneity, inconsistency and imprecision were considered overlapping to avoid double-counting. Certainty ratings refer to the in vitro resistance profile and not to clinical treatment efficacy. ENR = enrofloxacin; FLO = florfenicol; PEN = penicillin G; TET = tetracycline; TUL = tulathromycin; CI = confidence interval.

[1] Direction and magnitude of potential bias were unclear, and funnel methods are difficult to interpret in proportional meta-analysis. These are concerns but it was not judged substantial enough to downgrade due to the unclarity

[2] For ENR, the *H. somni* confidence interval crossed the prespecified 10% resistance threshold, but confidence intervals for *M. haemolytica* and *P. multocida* remained below 10%. Because the threshold crossing was limited to one pathogen-specific estimate and did not alter the antimicrobial-level interpretation of very low pooled resistance across pathogens, imprecision was noted but not downgraded.

#### Evidence base and pooled estimates underlying the GRADE-informed certainty assessment.

| Antimicrobial | Pathogen | Studies, n | Isolates, N | Pooled resistance % [95% CI] | $I^2$ (%) | Range of study-specific resistance estimates, % | CI thresholds crossed | Notes |
|---|---|---|---|---|---|---|---|---|

| | | | | | | | | |
|---|---|---|---|---|---|---|---|---|
| FLO | *H. somni* | 12 | 706 | 1 [0–3] | 0.0 | 0–4% | 0 | Consistently very low |
| FLO | *M. haemolytica* | 18 | 2215 | 1 [0–6] | 87.8 | 0–47% | 0 | Several high outliers |
| FLO | *P. multocida* | 19 | 2173 | 2 [1–6] | 92.6 | 0–59% | 0 | Several high outliers |
| ENR | *H. somni* | 11 | 675 | 2 [0–13] | 80.3 | 0–65% | 1 | One high outlier |
| ENR | *M. haemolytica* | 17 | 2201 | 2 [0–7] | 94.9 | 0–69% | 0 | High cluster/outliers |
| ENR | *P. multocida* | 18 | 2040 | 0 [0–3] | 91.9 | 0–58% | 0 | One high outlier |
| PEN | *H. somni* | 7 | 494 | 6 [1–29] | 92.7 | 0–65% | 2 | One high outlier |
| PEN | *M. haemolytica* | 12 | 1511 | 14 [5–33] | 93.9 | 0–71% | 3 | Broad spread |
| PEN | *P. multocida* | 12 | 1558 | 5 [1–16] | 92.2 | 0–89% | 1 | One high outlier |
| TUL | *H. somni* | 10 | 729 | 8 [1–36] | 80.3 | 0–91% | 3 | One extreme outlier |
| TUL | *M. haemolytica* | 15 | 2185 | 13 [4–32] | 96.0 | 0–77% | 3 | Broad spread |
| TUL | *P. multocida* | 14 | 1781 | 10 [4–27] | 95.9 | 0–81% | 2 | Broad spread |
| TET | *H. somni* | 7 | 562 | 19 [4–60] | 87.4 | 0–63% | 3 | Low/high clusters |
| TET | *M. haemolytica* | 12 | 1400 | 28 [12–51] | 95.9 | 0–87% | 2 | Broad spread |
| TET | *P. multocida* | 13 | 1724 | 36 [12–70] | 95.2 | 0–100% | 2 | Broad spread |

Thresholds used for imprecision were 10%, 20%, and 30%. “CI thresholds crossed” indicates the number of prespecified thresholds crossed by the pooled 95% CI. The range of study-specific resistance estimates was used to support assessment of inconsistency and indicates the lowest and highest resistance proportions reported by individual studies contributing to each pathogen–antimicrobial meta-analysis; it is not a confidence interval. $I^2$ is presented descriptively and interpreted cautiously in proportional meta-analysis. ENR = enrofloxacin; FLO = florfenicol; PEN = penicillin G; TET = tetracycline; TUL = tulathromycin; CI = confidence interval.

### Supplementary Material IV

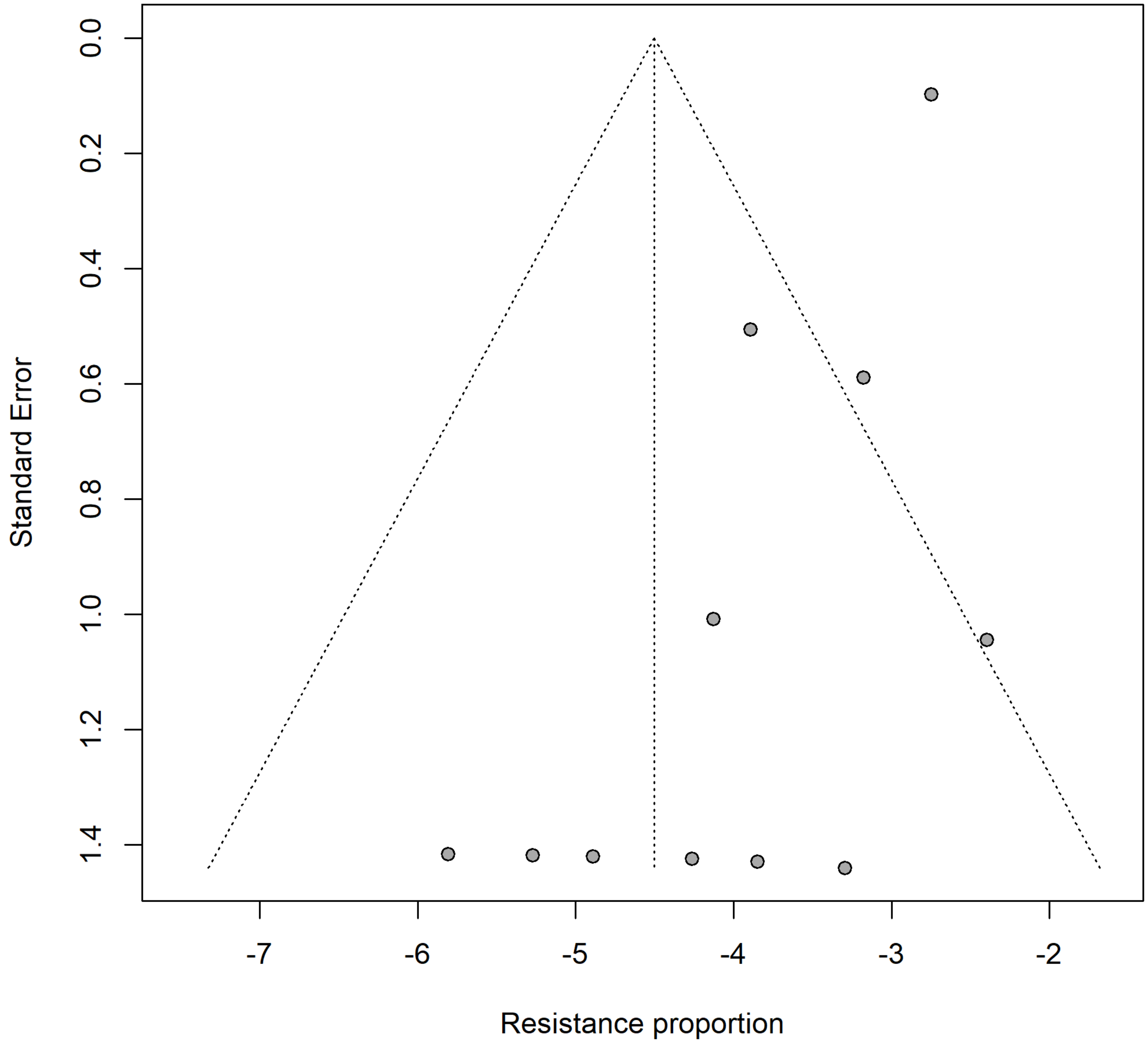


Supplementary figure S4.1. Funnel plot for the meta−analysis of enrofloxacin resistance in Histophilus somni. Egger's linear regression test for funnel−plot asymmetry (k = 11 studies): p = 0.002.

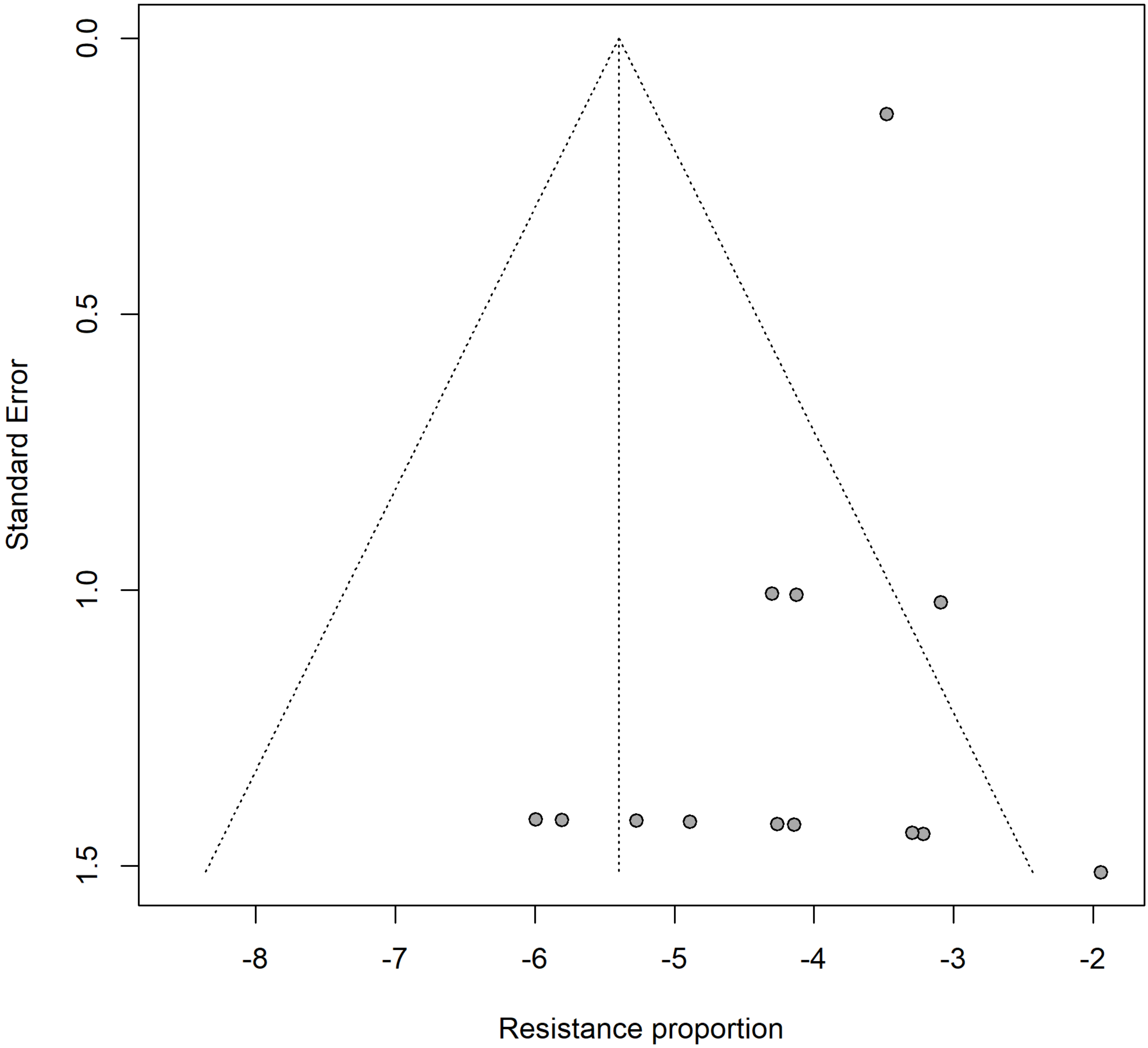


Supplementary figure S4.2. Funnel plot for the meta−analysis of florfenicol resistance in Histophilus somni. Egger's linear regression test for funnel−plot asymmetry (k = 13 studies): p = 0.062.

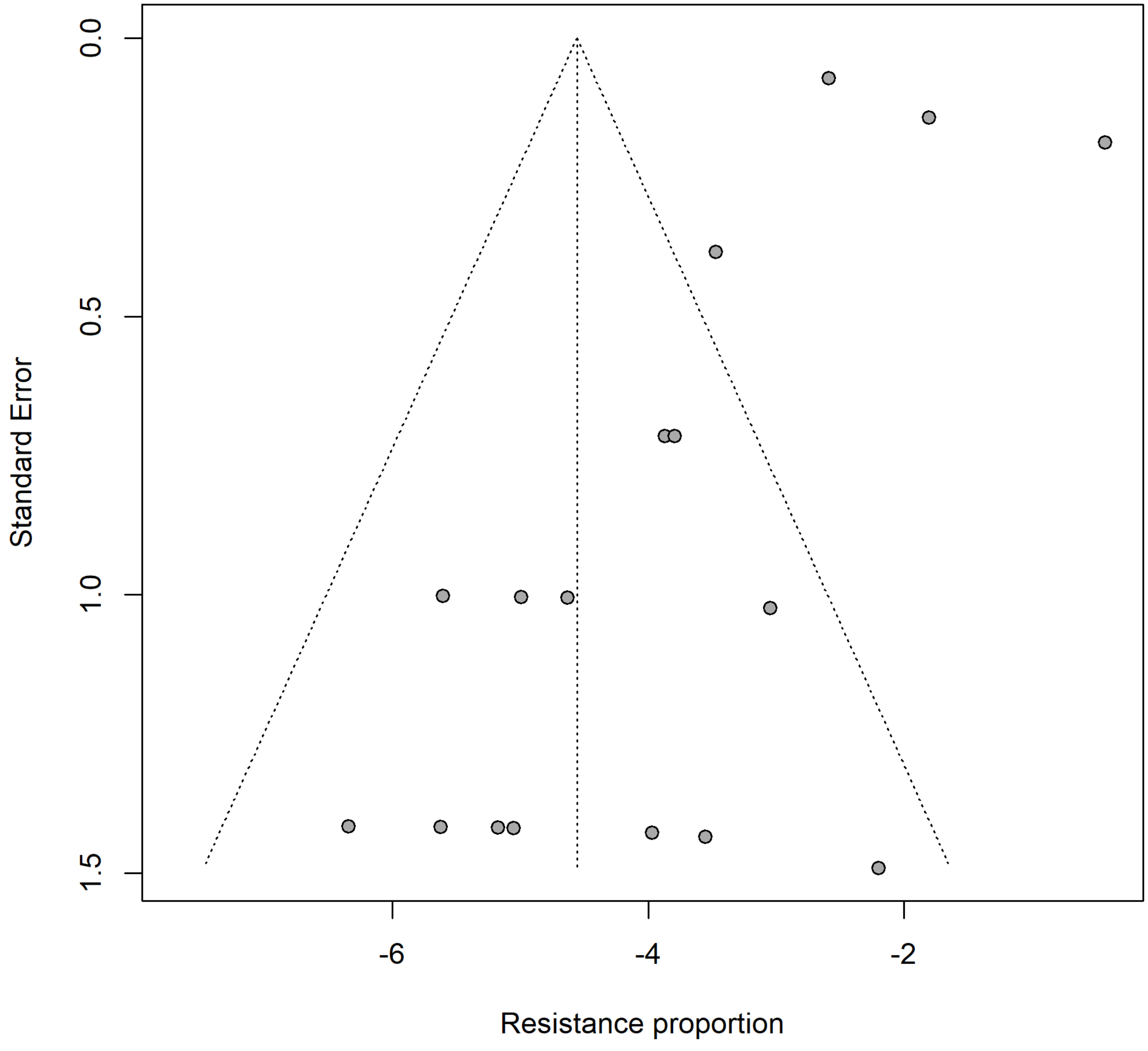


Supplementary figure S4.3. Funnel plot for the meta−analysis of enrofloxacin resistance in Mannheimia haemolytica. Egger's linear regression test for funnel−plot asymmetry (k = 17 studies): p = 0.157.

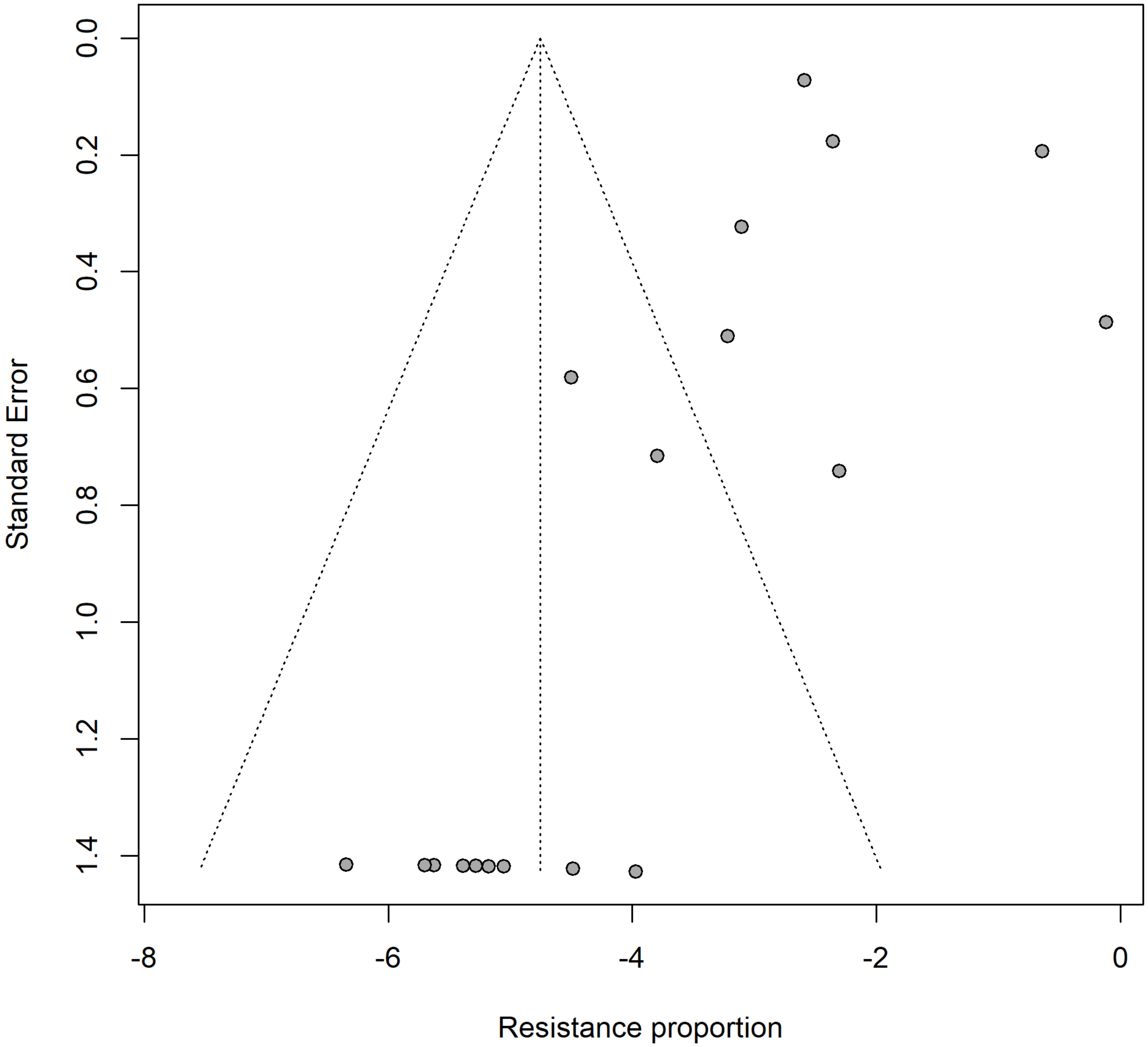


Supplementary figure S4.4. Funnel plot for the meta−analysis of florfenicol resistance in Mannheimia haemolytica. Egger's linear regression test for funnel−plot asymmetry (k = 18 studies): p = 0.196.

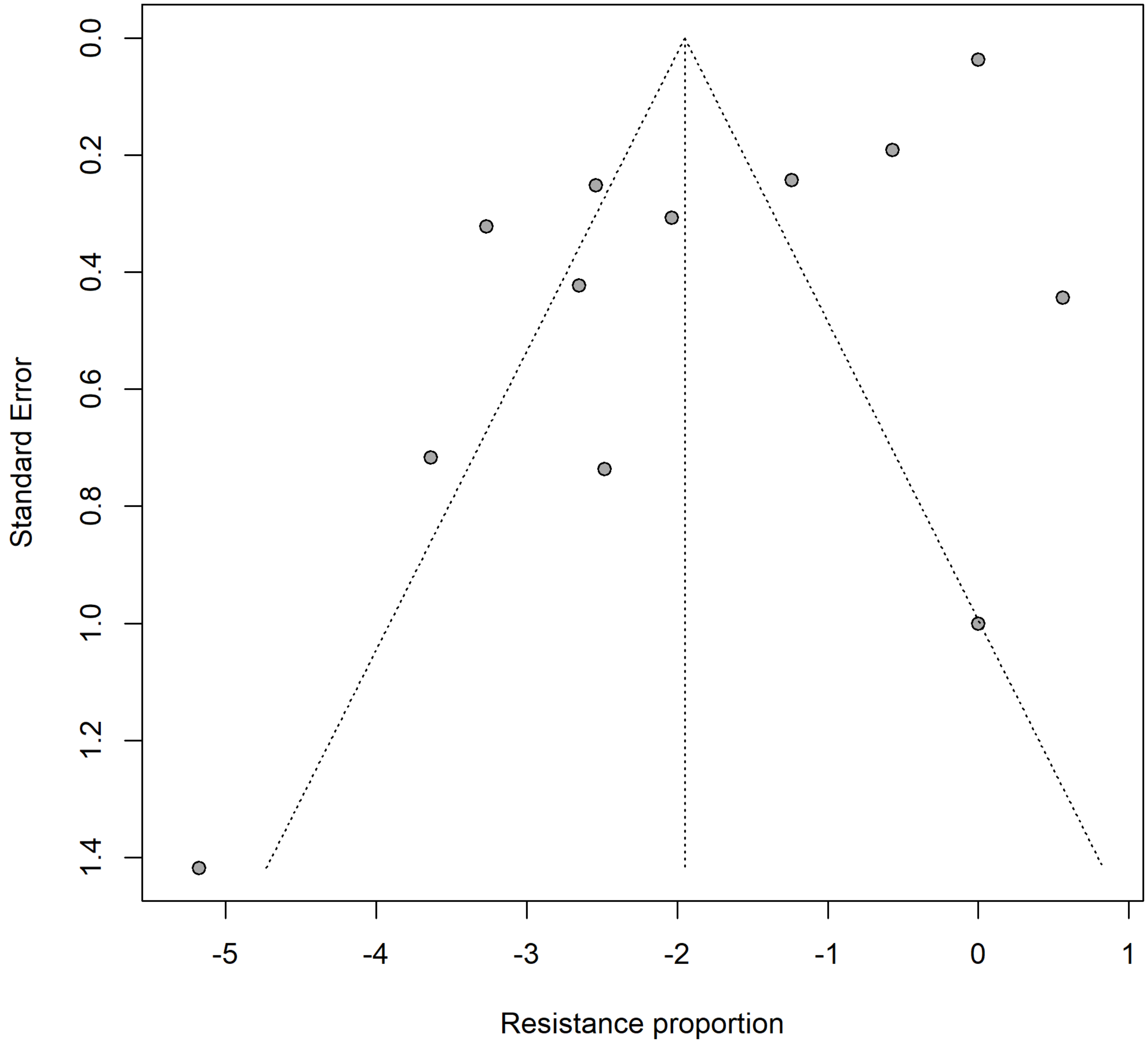


Supplementary figure S4.5. Funnel plot for the meta−analysis of penicillin resistance in Mannheimia haemolytica. Egger's linear regression test for funnel−plot asymmetry (k = 12 studies): p = 0.003.

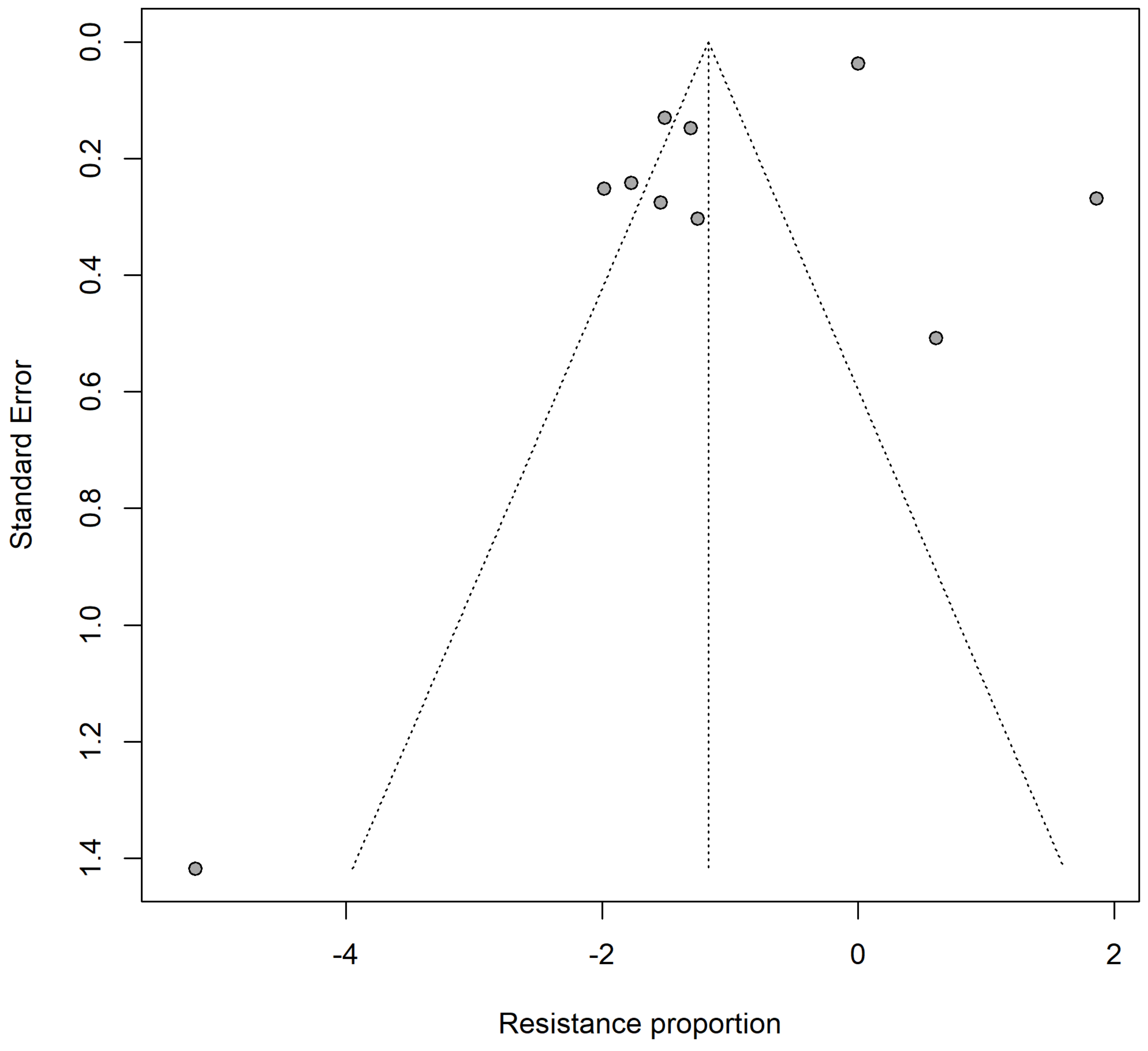


Supplementary figure S4.6. Funnel plot for the meta−analysis of tetracycline resistance in Mannheimia haemolytica. Egger's linear regression test for funnel−plot asymmetry (k = 10 studies): p = 0.098.

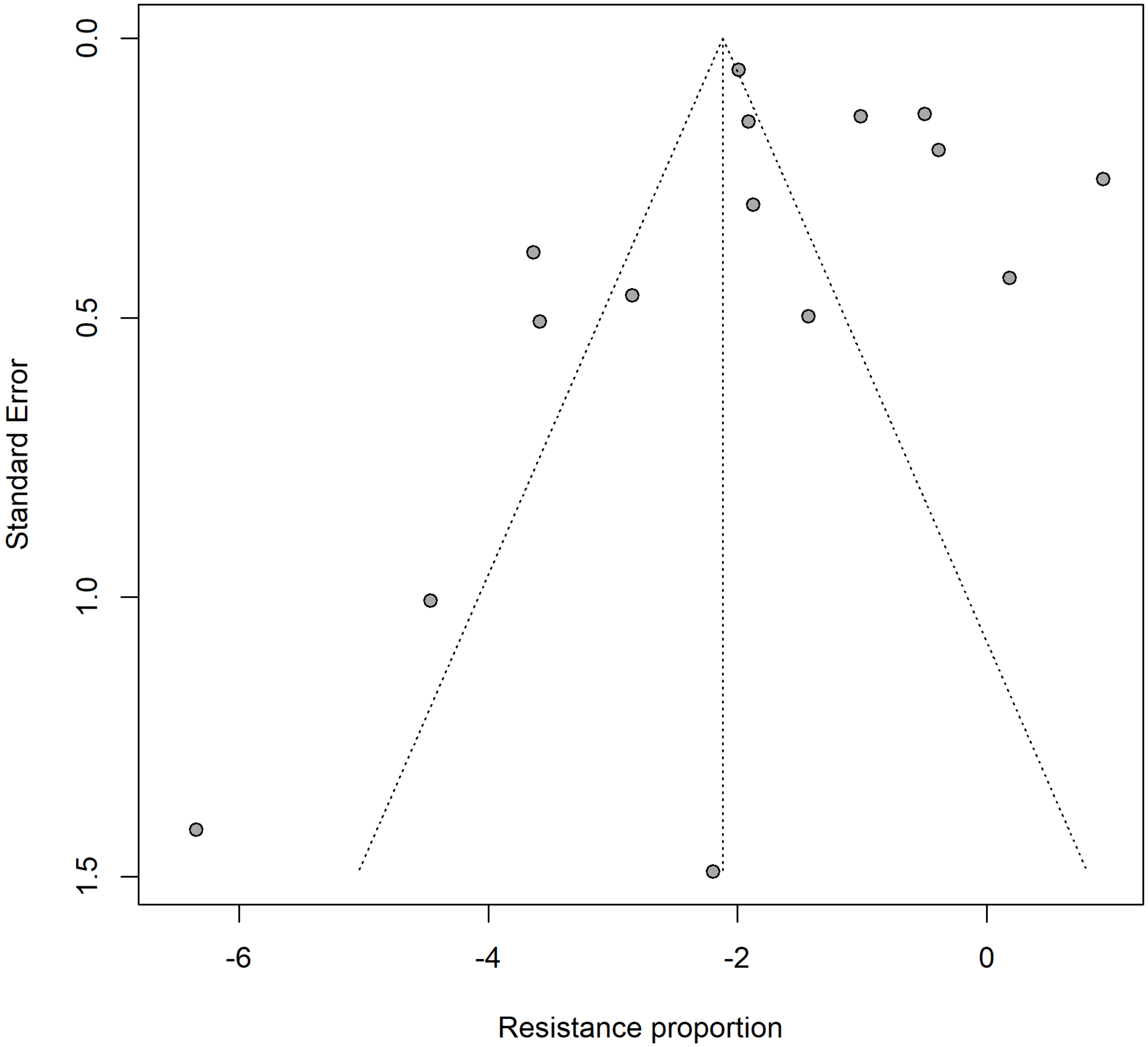


Supplementary figure S4.7. Funnel plot for the meta−analysis of tulathromycin resistance in Mannheimia haemolytica. Egger's linear regression test for funnel−plot asymmetry (k = 15 studies): p = 0.745.

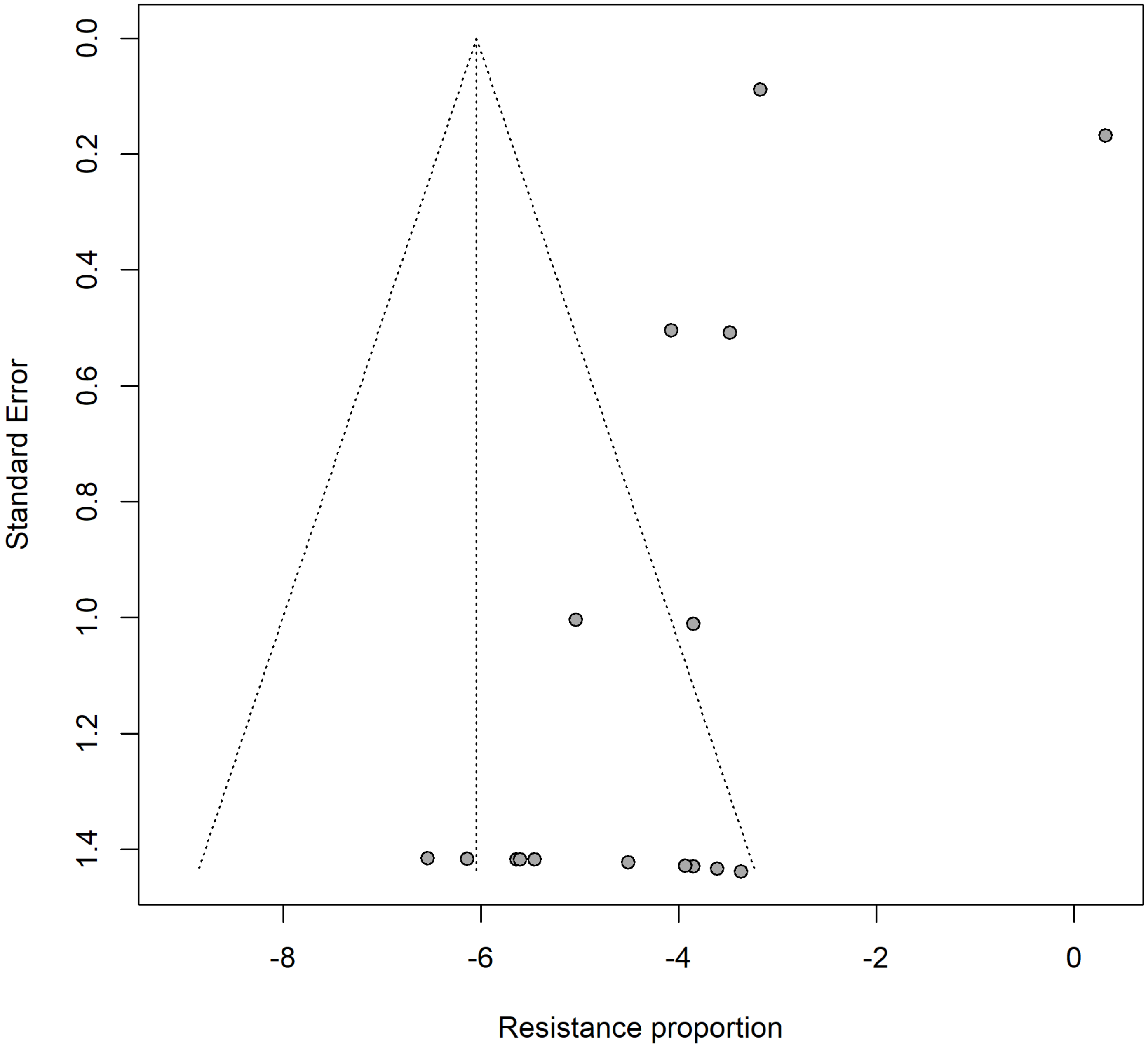


Supplementary figure S4.8. Funnel plot for the meta−analysis of enrofloxacin resistance in Pasteurella multocida. Egger's linear regression test for funnel−plot asymmetry (k = 17 studies): p = 0.385.

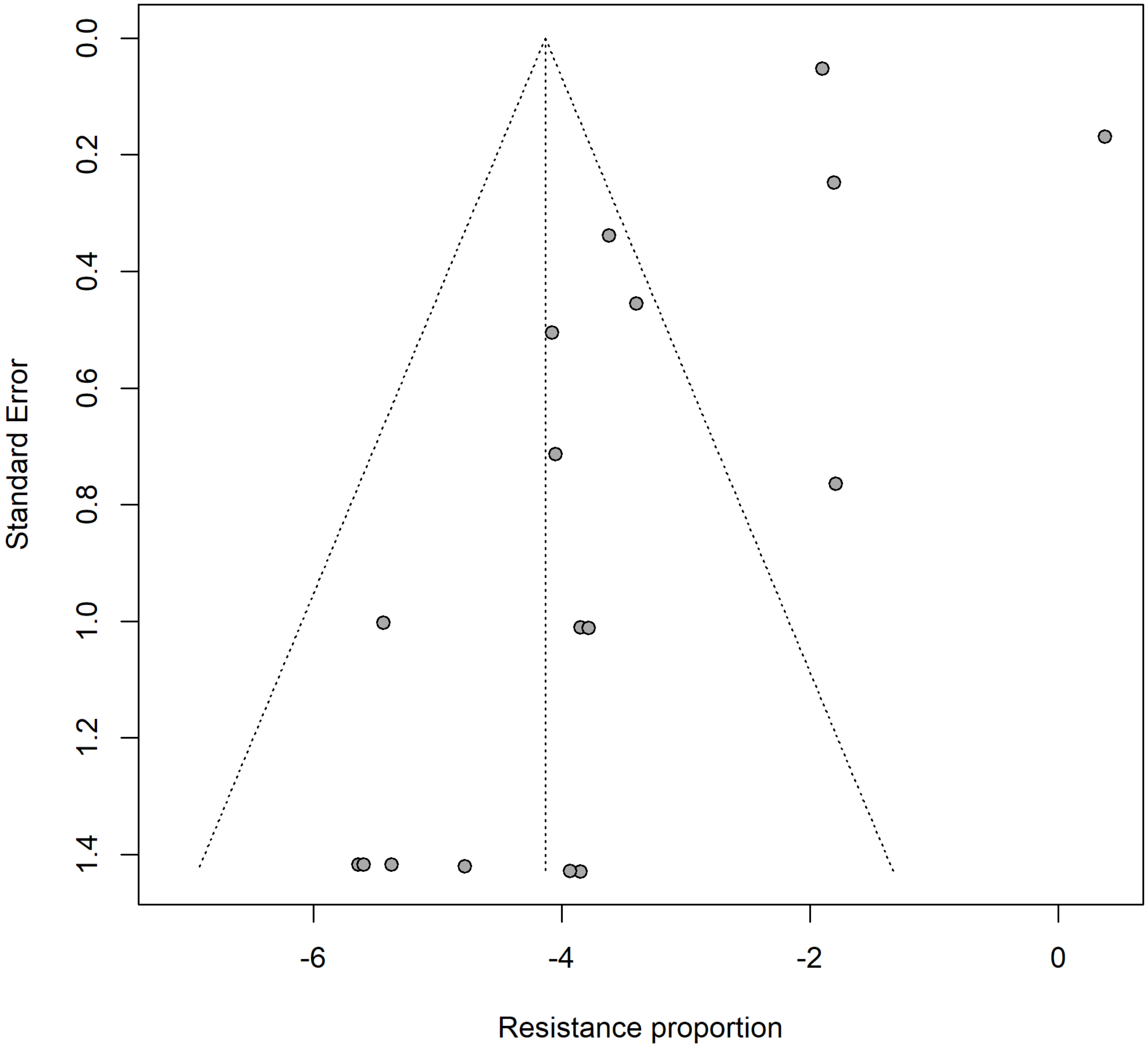


Supplementary figure S4.9. Funnel plot for the meta−analysis of florfenicol resistance in Pasteurella multocida. Egger's linear regression test for funnel−plot asymmetry (k = 17 studies): p = 0.100.

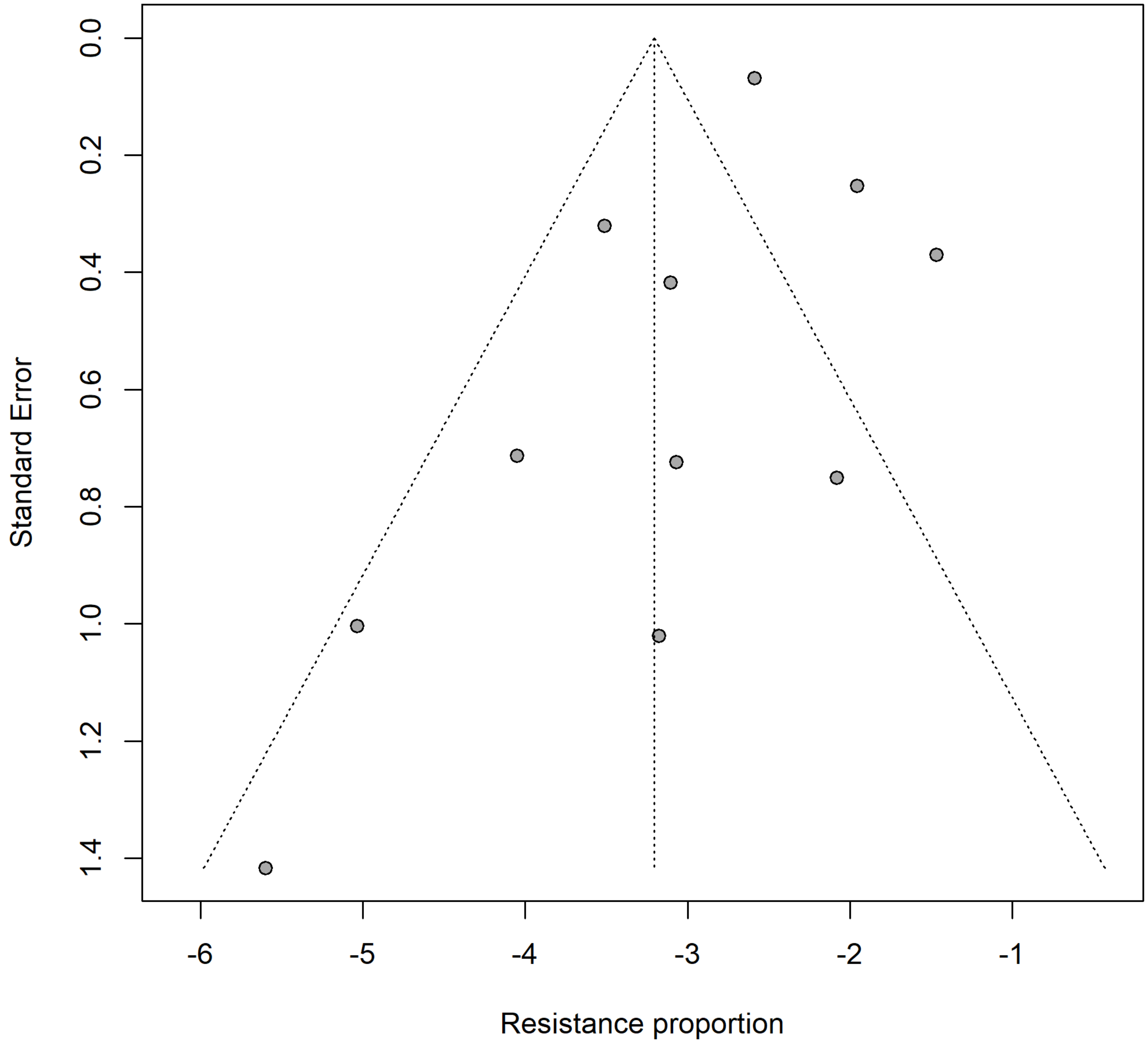


Supplementary figure S4.10. Funnel plot for the meta−analysis of penicillin resistance in Pasteurella multocida. Egger's linear regression test for funnel−plot asymmetry (k = 11 studies): p = 0.310.

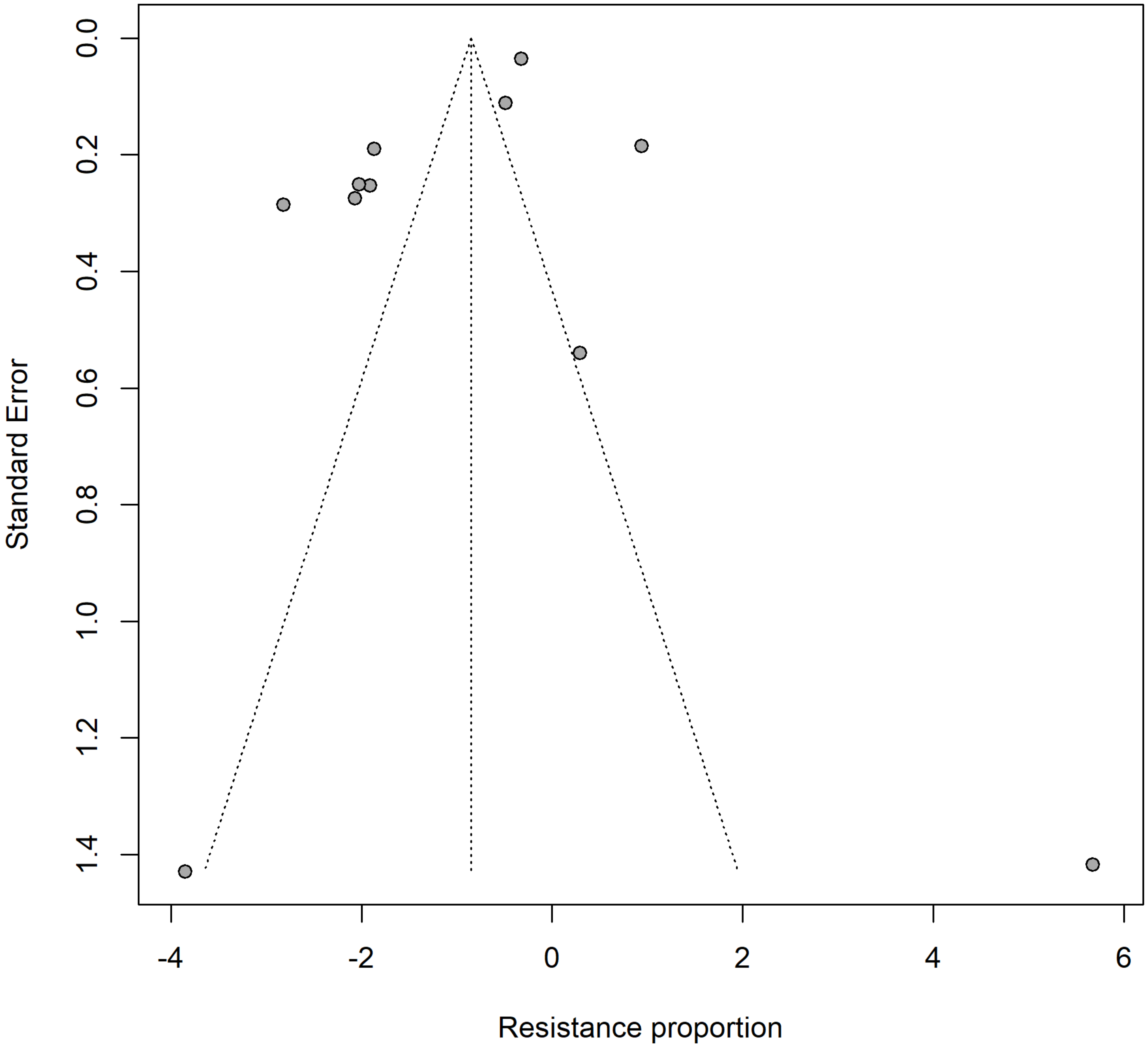


Supplementary figure S4.11. Funnel plot for the meta−analysis of tetracycline resistance in Pasteurella multocida. Egger's linear regression test for funnel−plot asymmetry (k = 11 studies): p = 0.187.

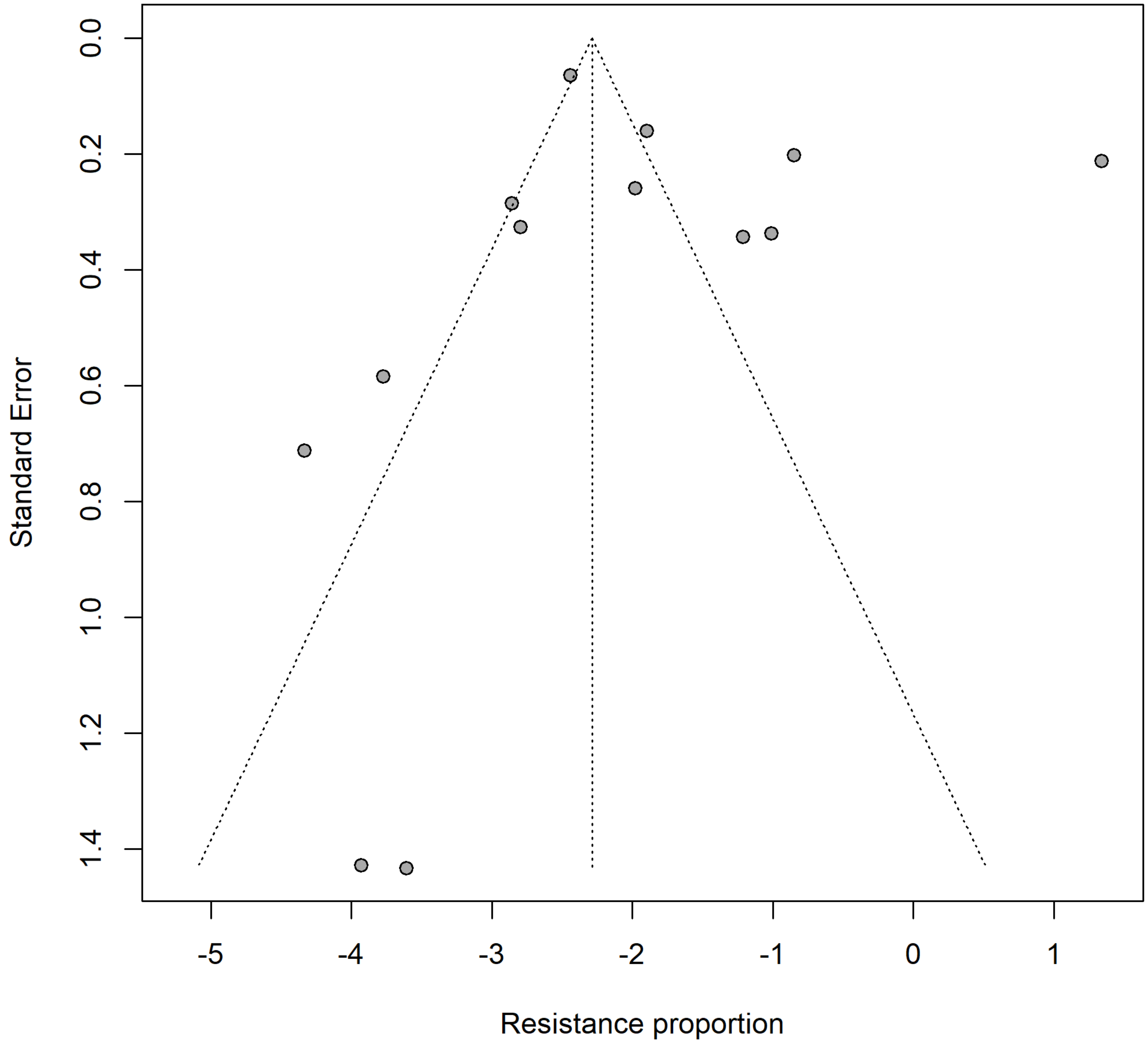


Supplementary figure S4.12. Funnel plot for the meta−analysis of tulathromycin resistance in Pasteurella multocida. Egger's linear regression test for funnel−plot asymmetry (k = 13 studies): p = 0.571.

### Supplementary Material VI

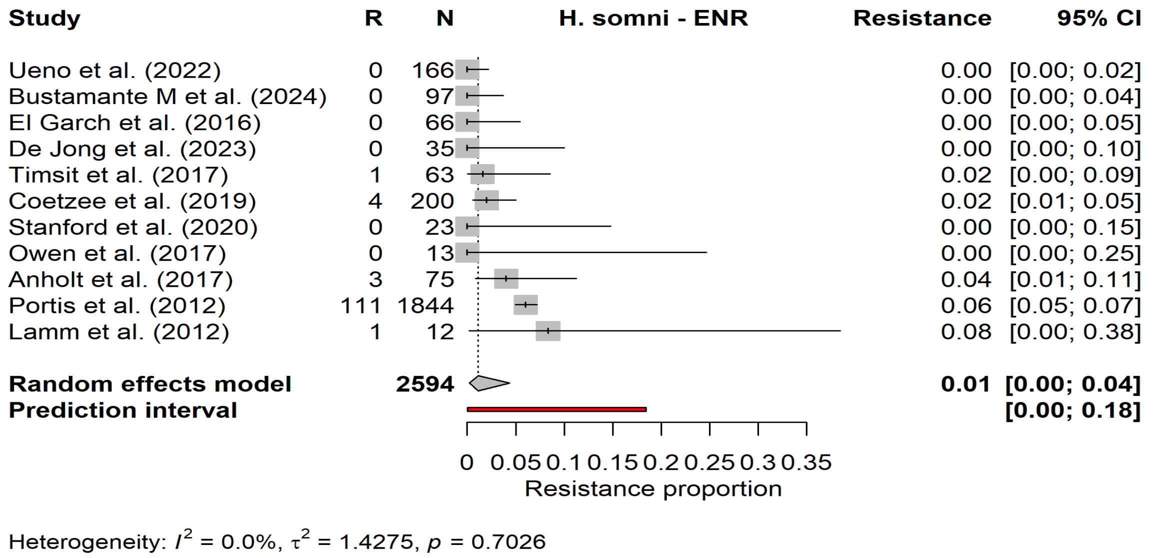


Supplementary figure S6.1. Forest plot for the meta−analysis of enrofloxacin resistance in Histophilus somni.

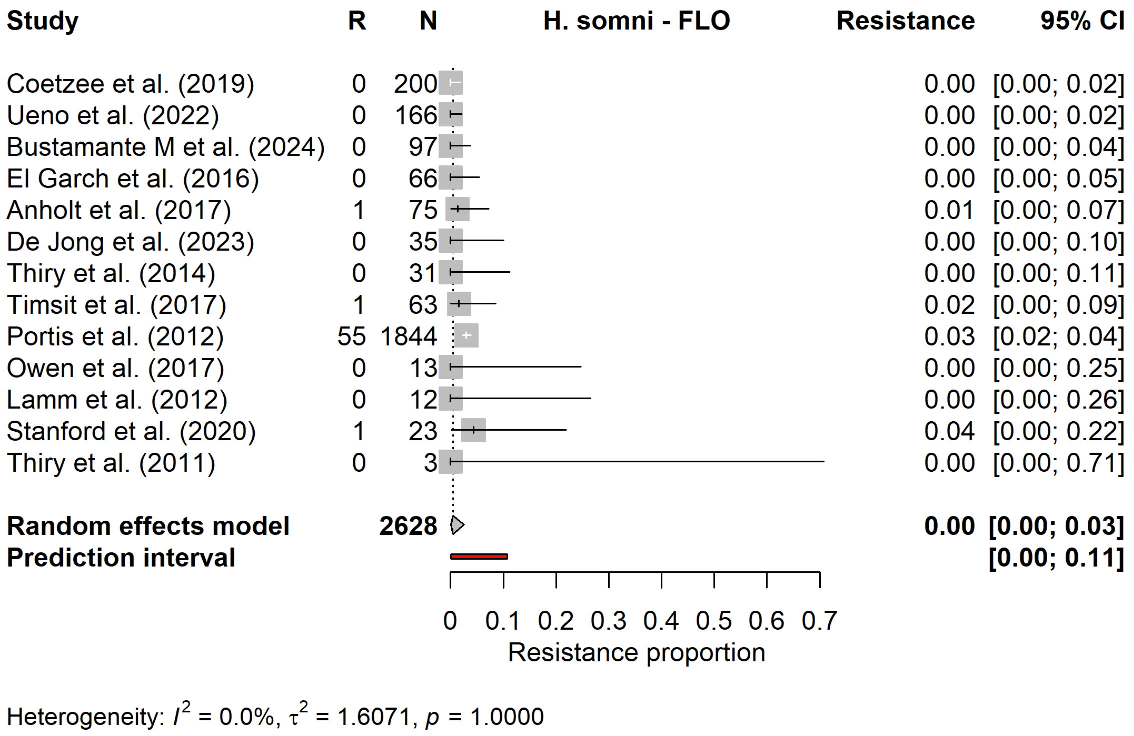


Supplementary figure S6.2. Forest plot for the meta−analysis of florfenicol resistance in Histophilus somni.

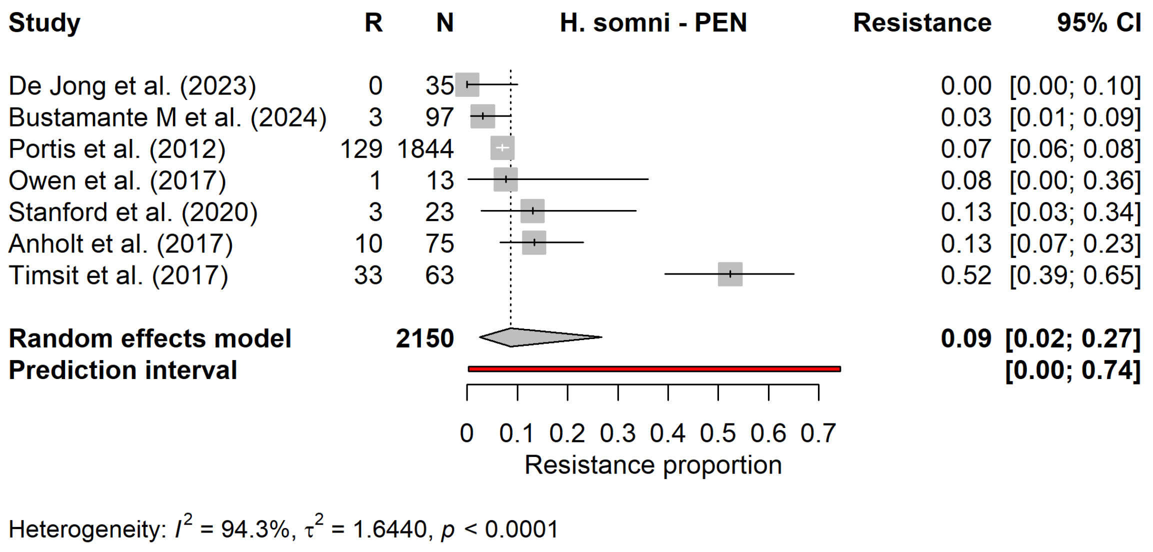


Supplementary figure S6.3. Forest plot for the meta−analysis of penicillin resistance in Histophilus somni.

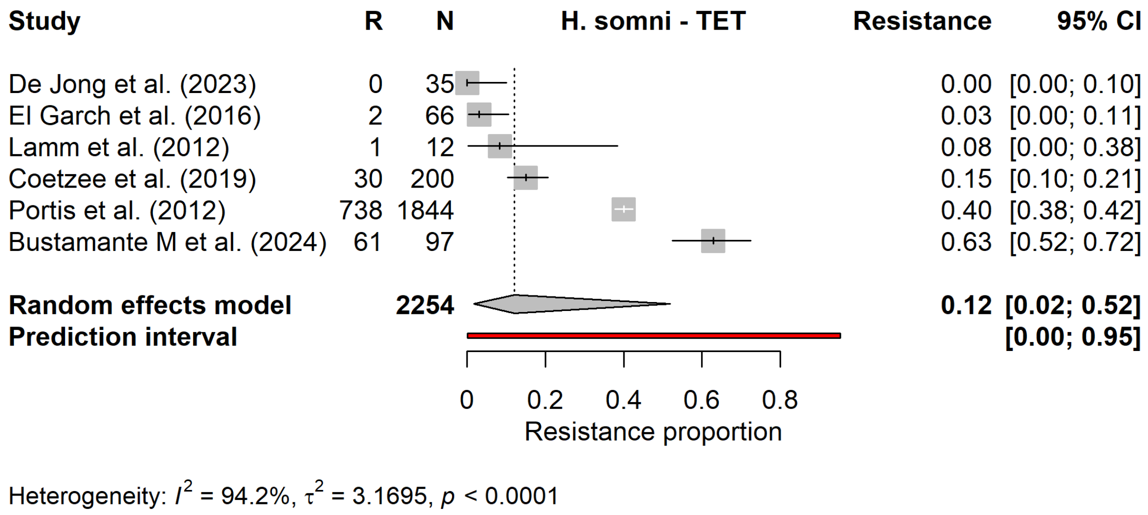


Supplementary figure S6.4. Forest plot for the meta−analysis of tetracycline resistance in Histophilus somni.

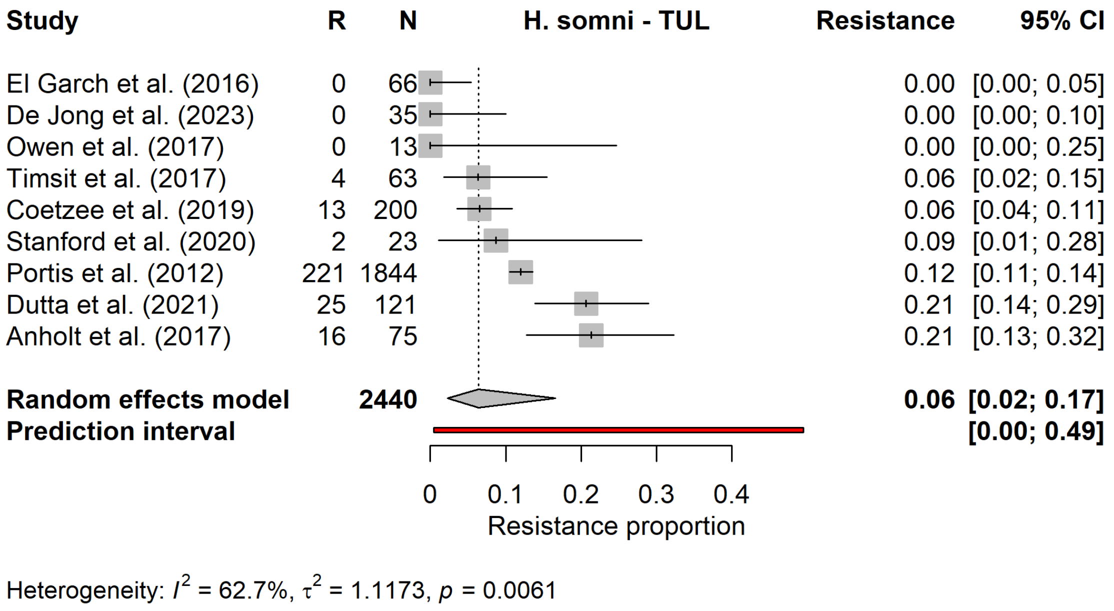


Supplementary figure S6.5. Forest plot for the meta−analysis of tulathromycin resistance in Histophilus somni.

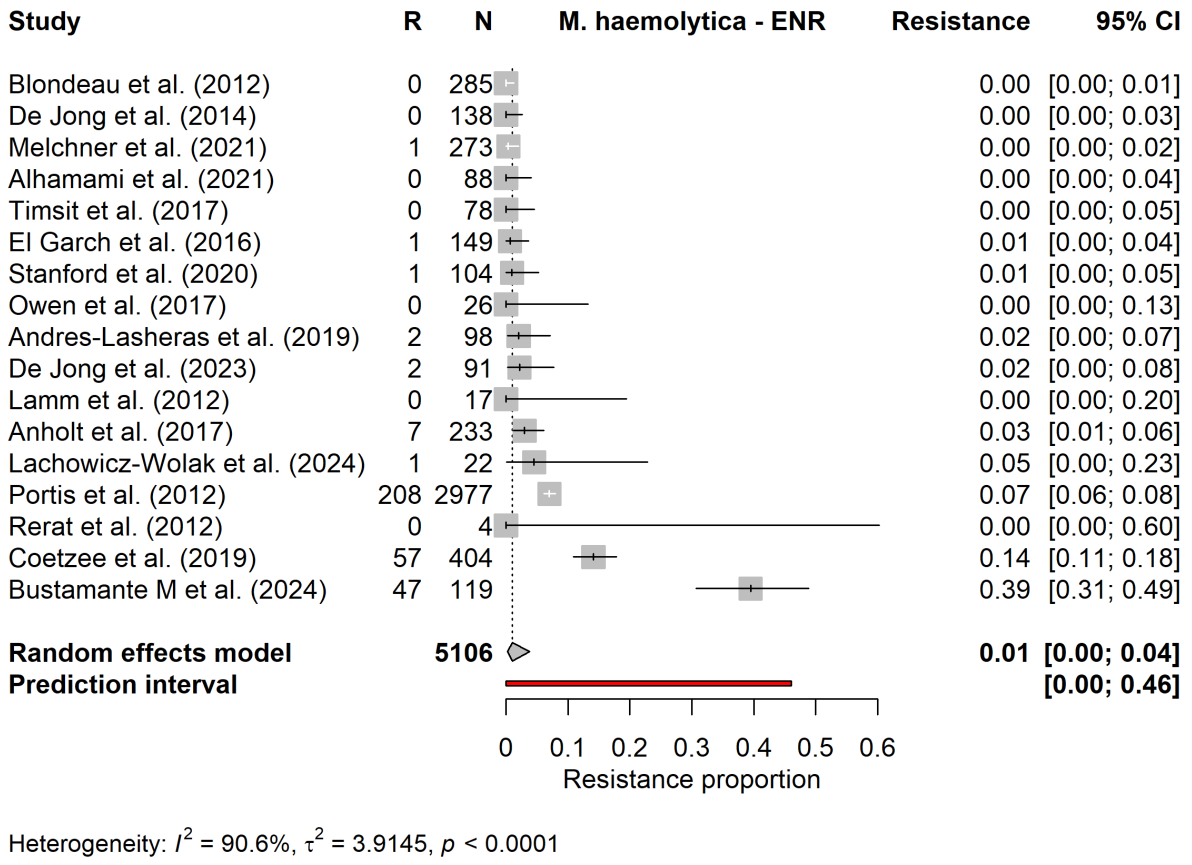


Supplementary figure S6.6. Forest plot for the meta−analysis of enrofloxacin resistance in Mannheimia haemolytica.

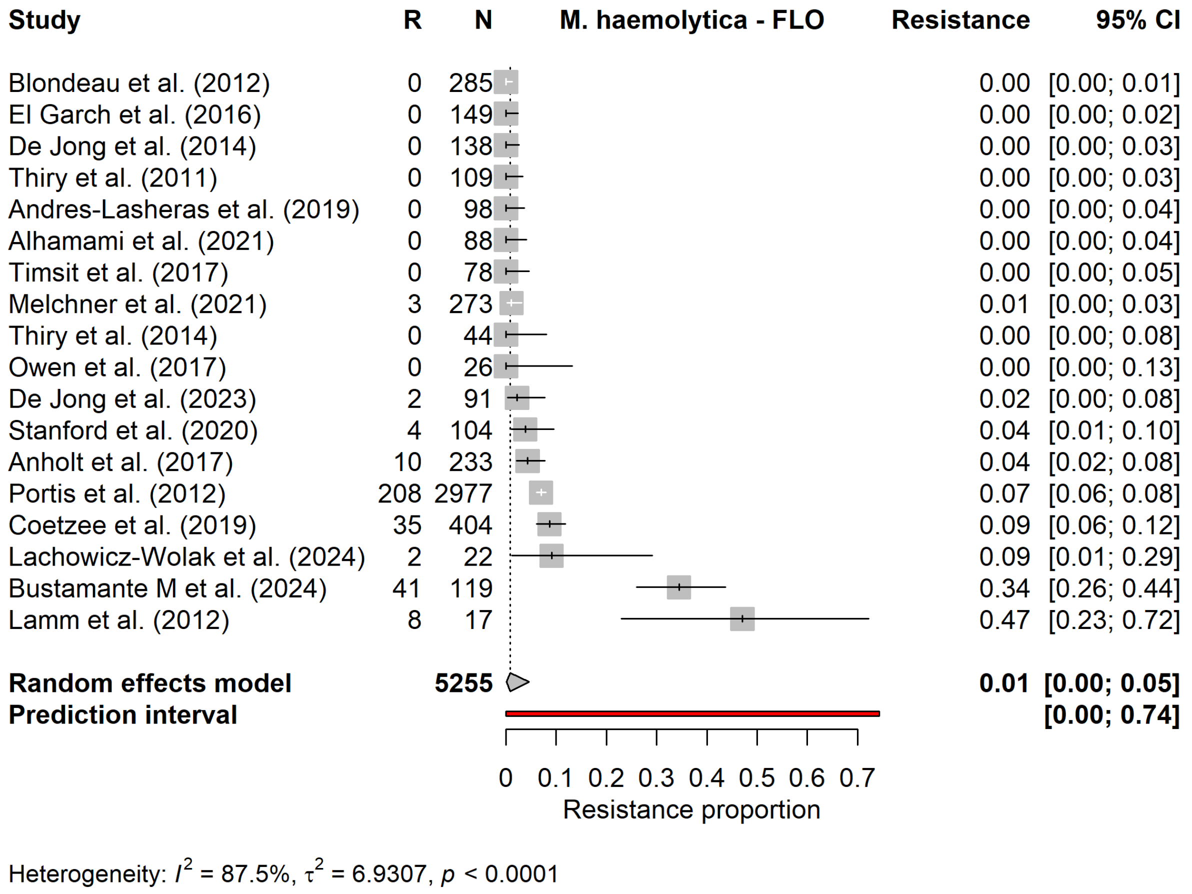


Supplementary figure S6.7. Forest plot for the meta−analysis of florfenicol resistance in Mannheimia haemolytica.

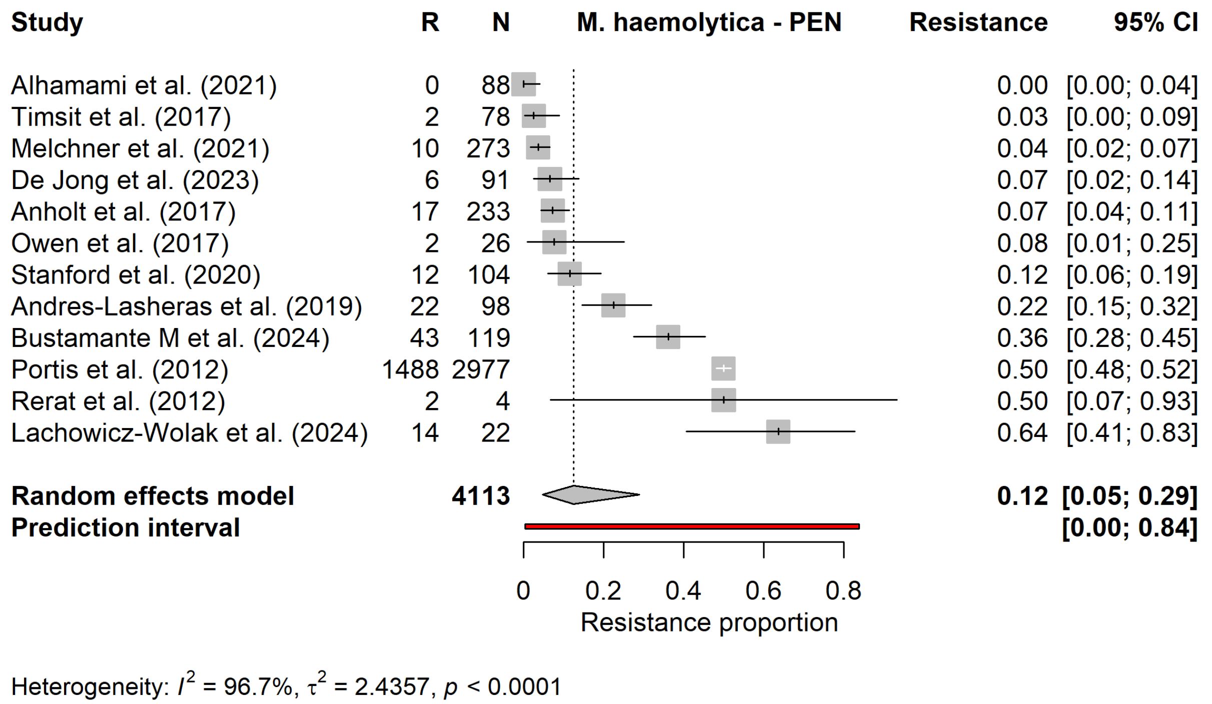


Supplementary figure S6.8. Forest plot for the meta−analysis of penicillin resistance in Mannheimia haemolytica.

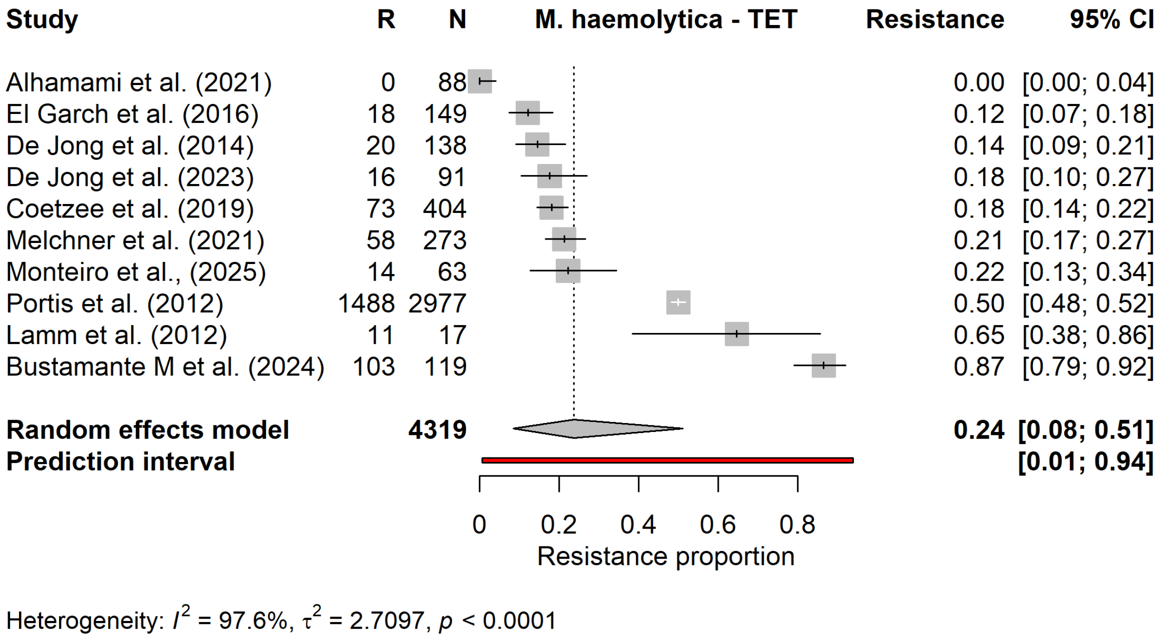


Supplementary figure S6.9. Forest plot for the meta−analysis of tetracycline resistance in Mannheimia haemolytica.

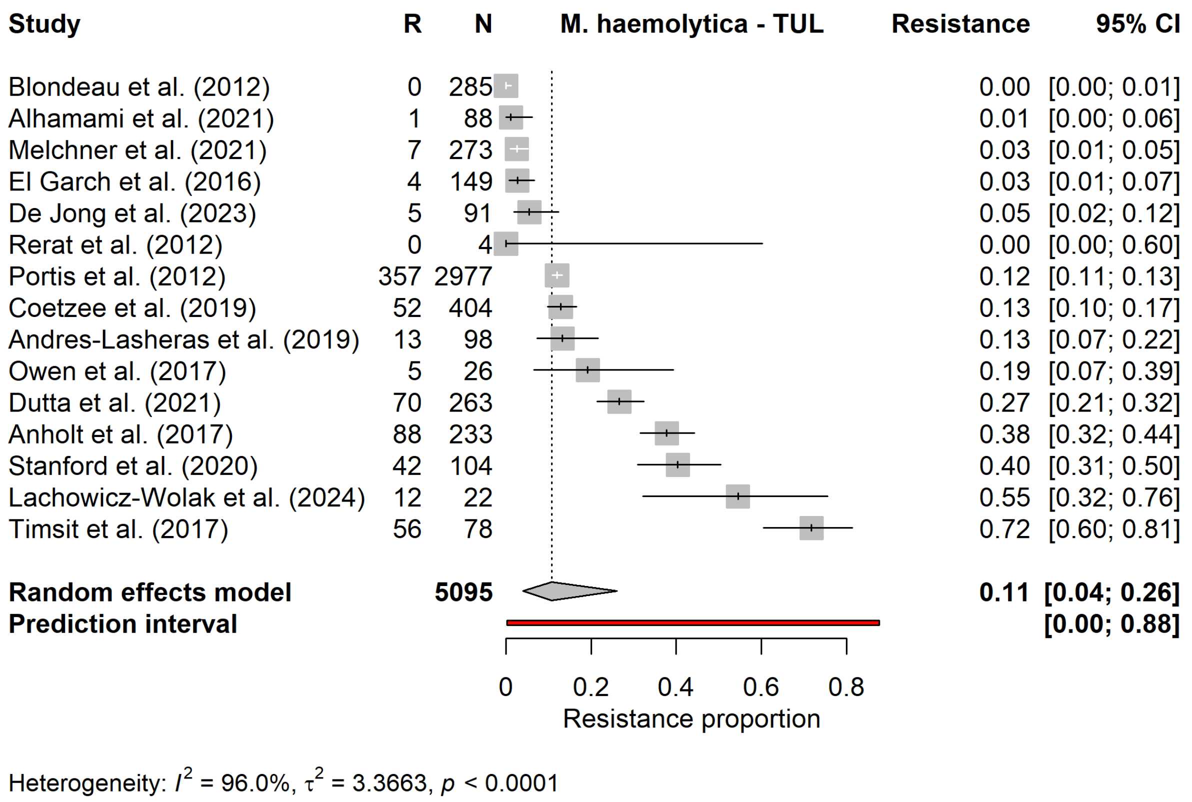


Supplementary figure S6.10. Forest plot for the meta−analysis of tulathromycin resistance in Mannheimia haemolytica.

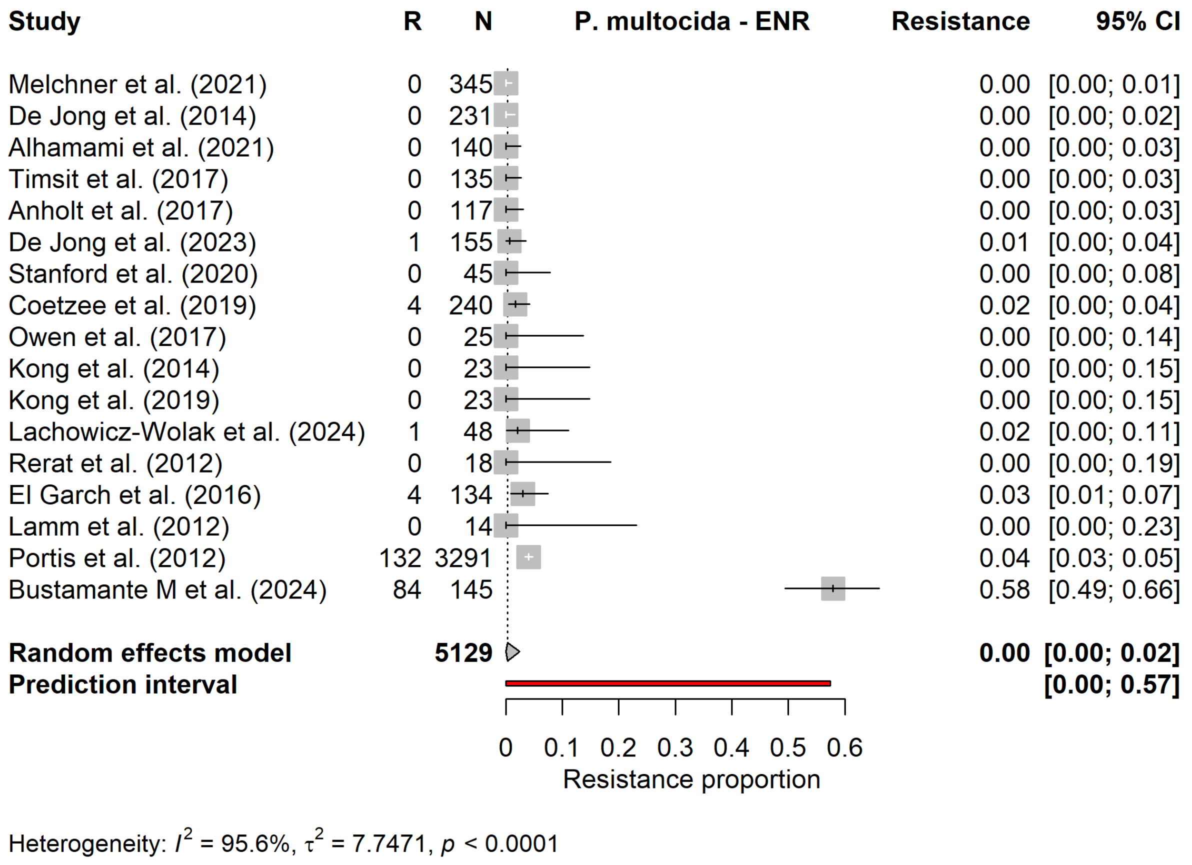


Supplementary figure S6.11. Forest plot for the meta−analysis of enrofloxacin resistance in Pasteurella multocida.

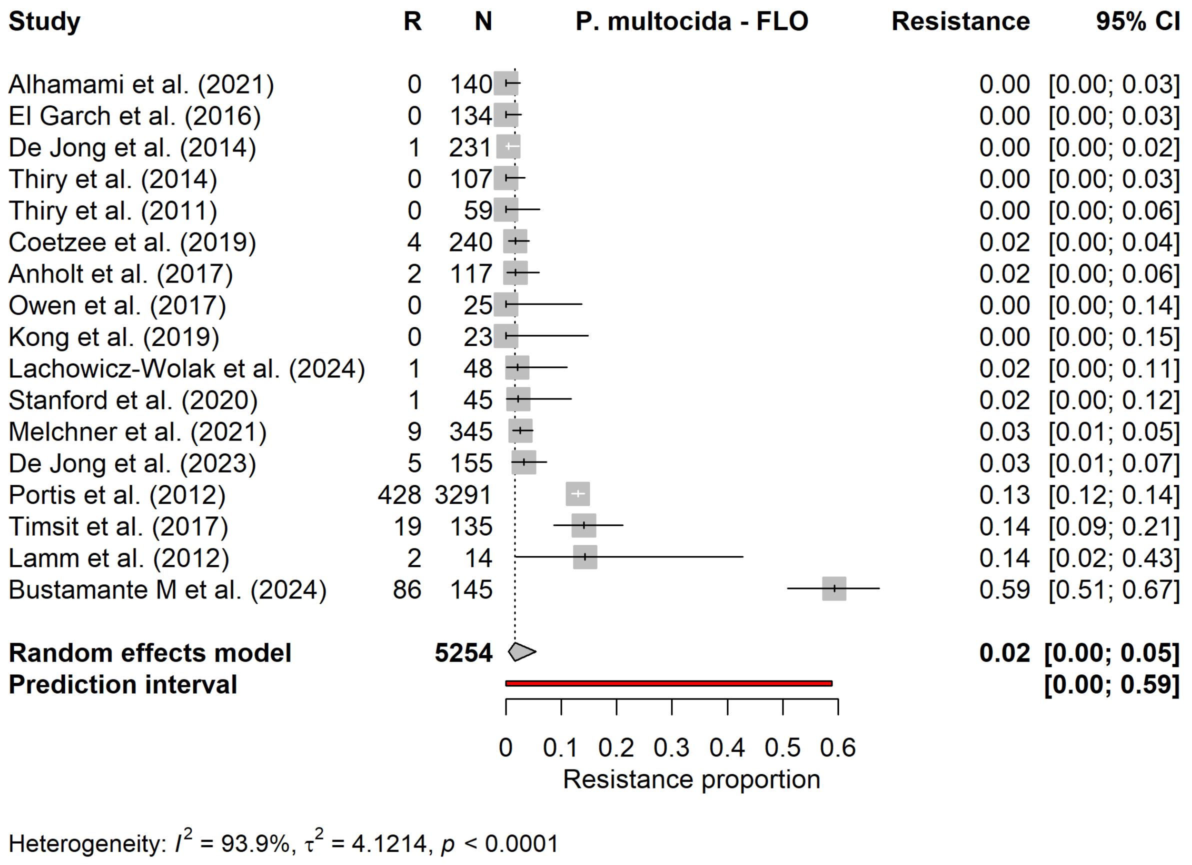


Supplementary figure S6.12. Forest plot for the meta−analysis of florfenicol resistance in Pasteurella multocida.

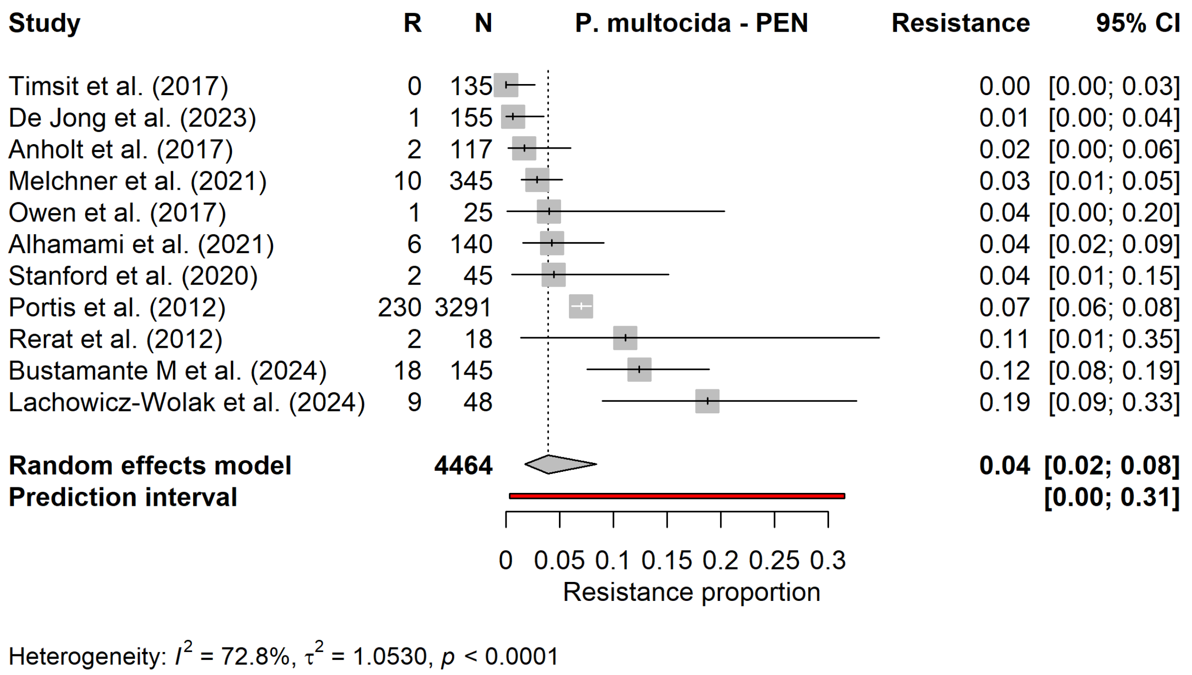


Supplementary figure S6.13. Forest plot for the meta−analysis of penicillin resistance in Pasteurella multocida.

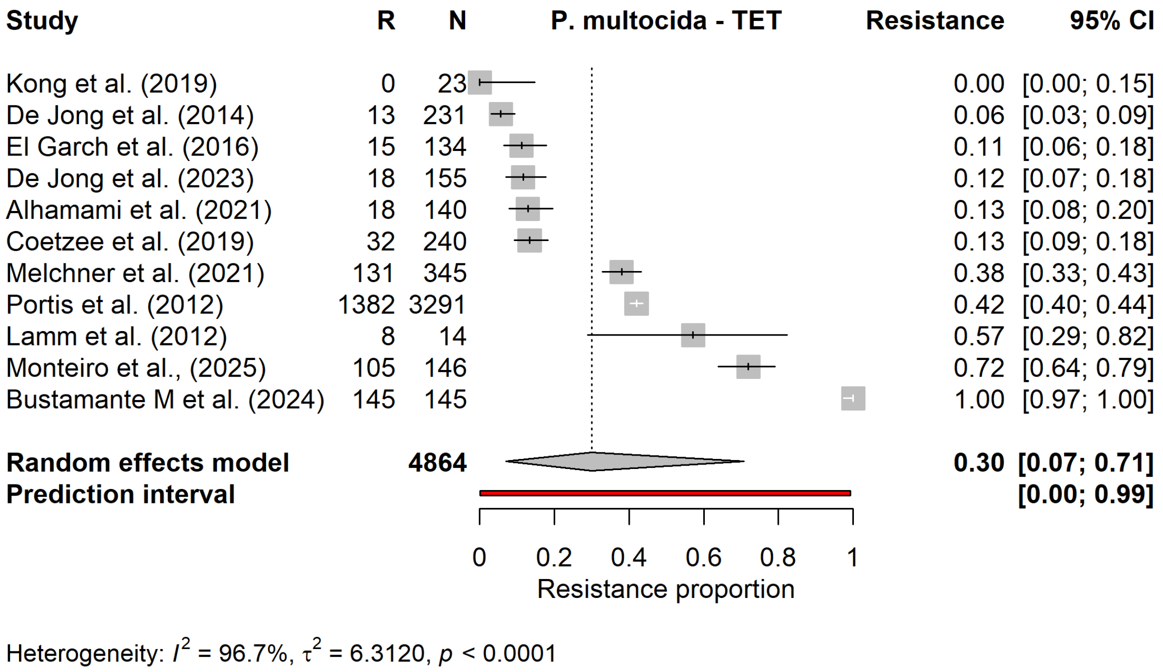


Supplementary figure S6.14. Forest plot for the meta−analysis of tetracycline resistance in Pasteurella multocida.

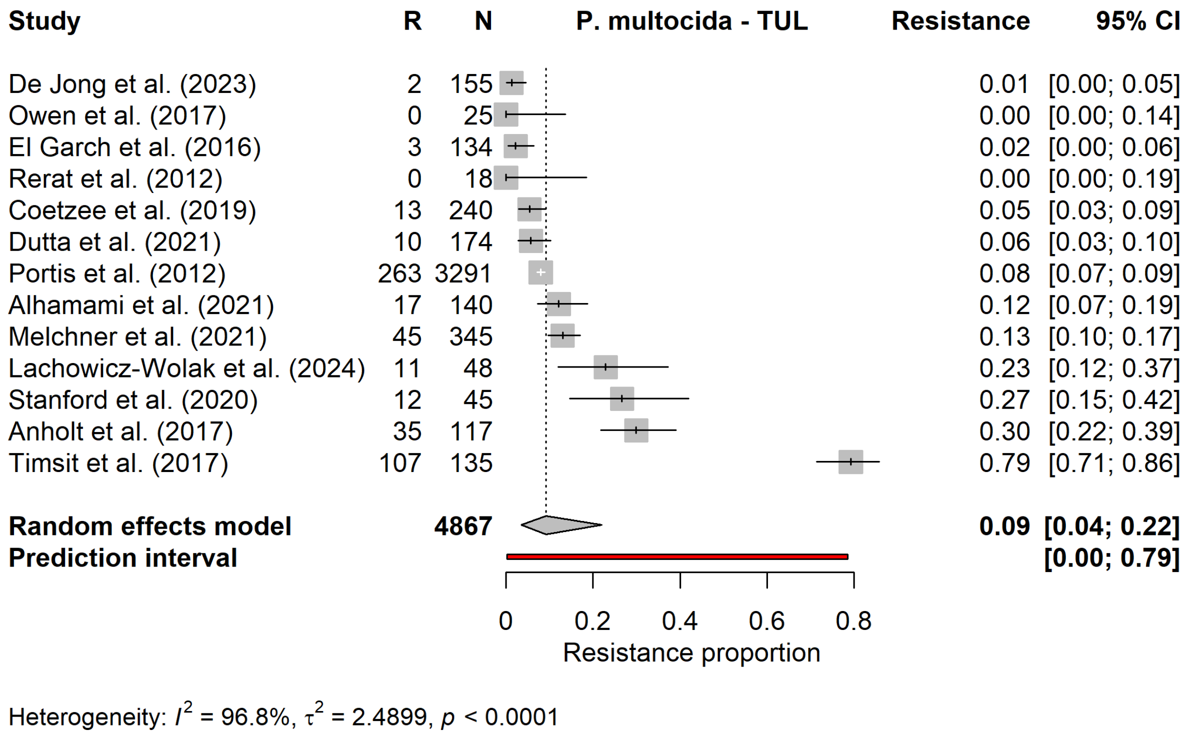


Supplementary figure S6.15. Forest plot for the meta−analysis of tulathromycin resistance in Pasteurella multocida.

### Supplementary Material VII

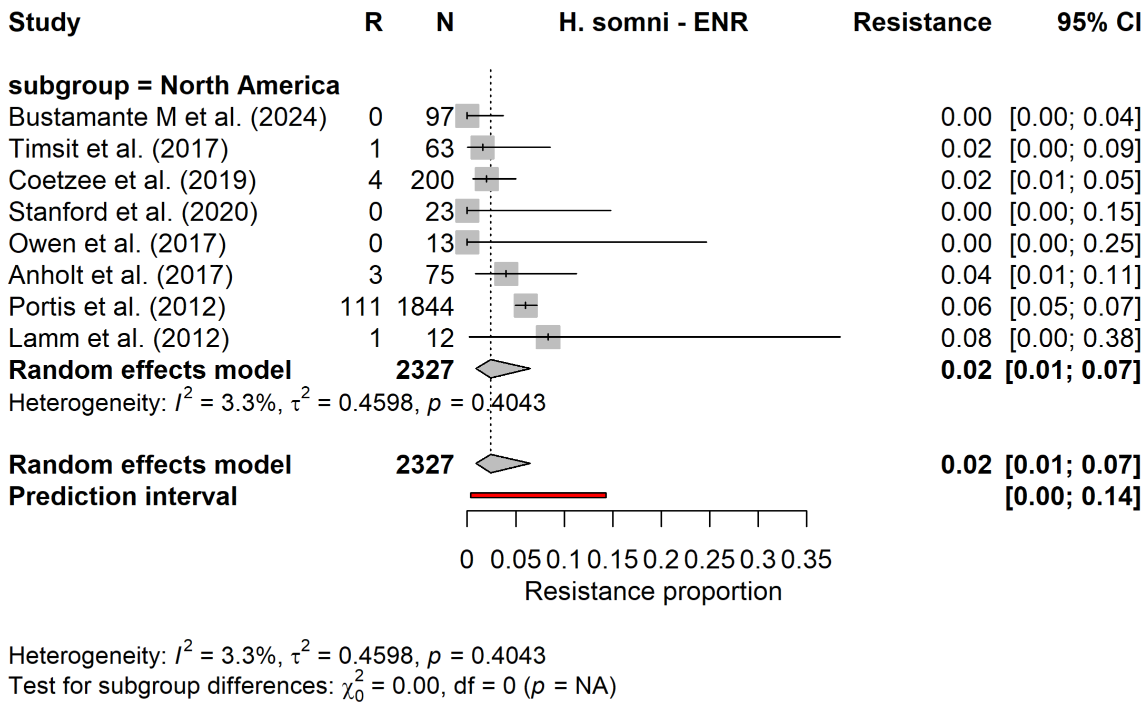


Supplementary figure S7.1. Forest plot for the subgroup meta−analysis of enrofloxacin resistance in Histophilus somni, stratified by continent.

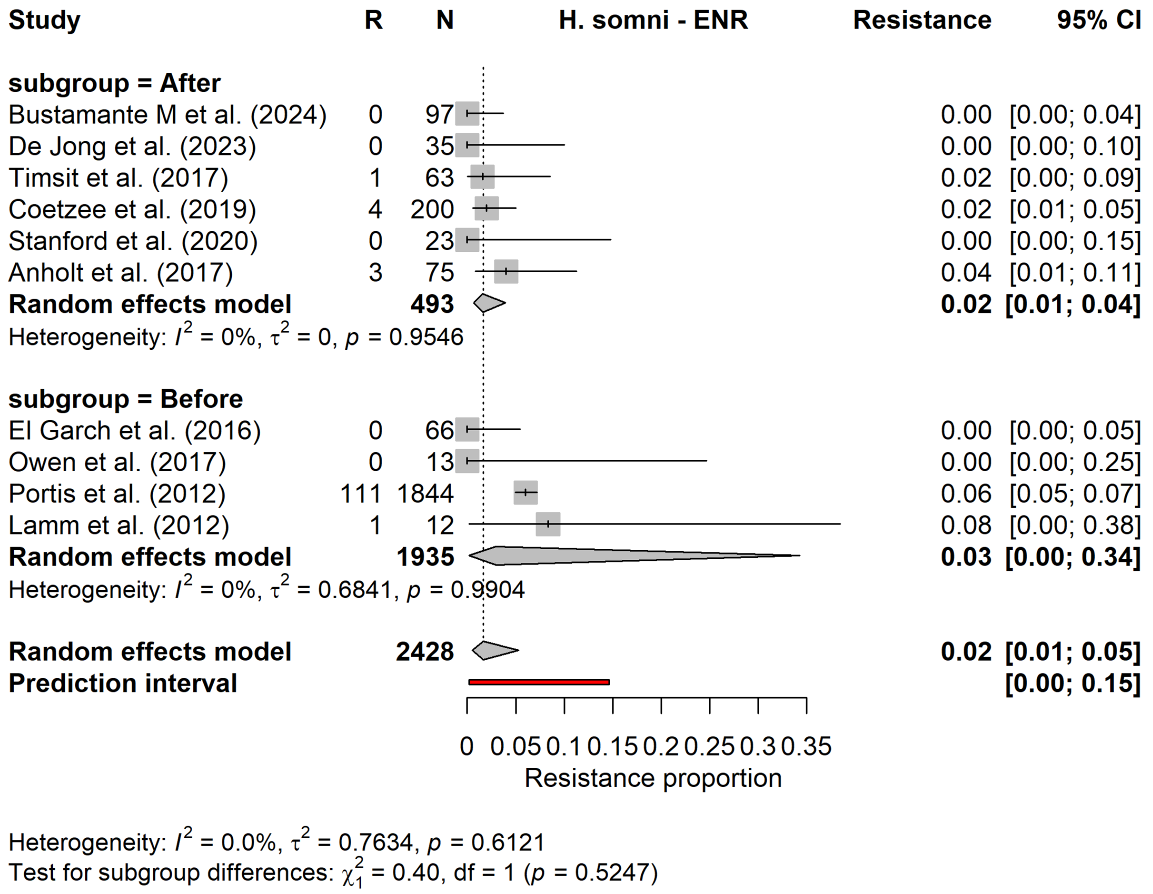


Supplementary figure S7.2. Forest plot for the subgroup meta−analysis of enrofloxacin resistance in Histophilus somni, stratified by study period.

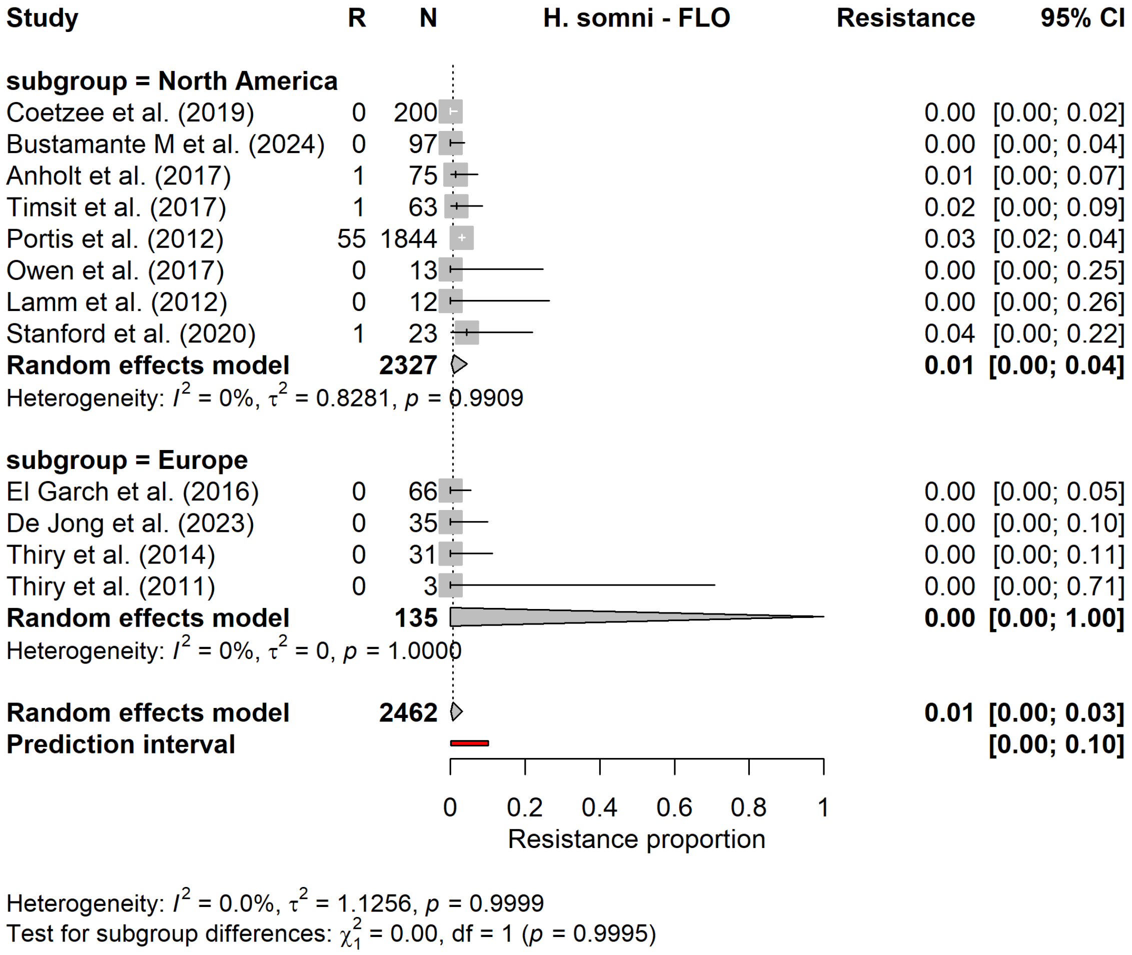


Supplementary figure S7.3. Forest plot for the subgroup meta−analysis of florfenicol resistance in Histophilus somni, stratified by continent.

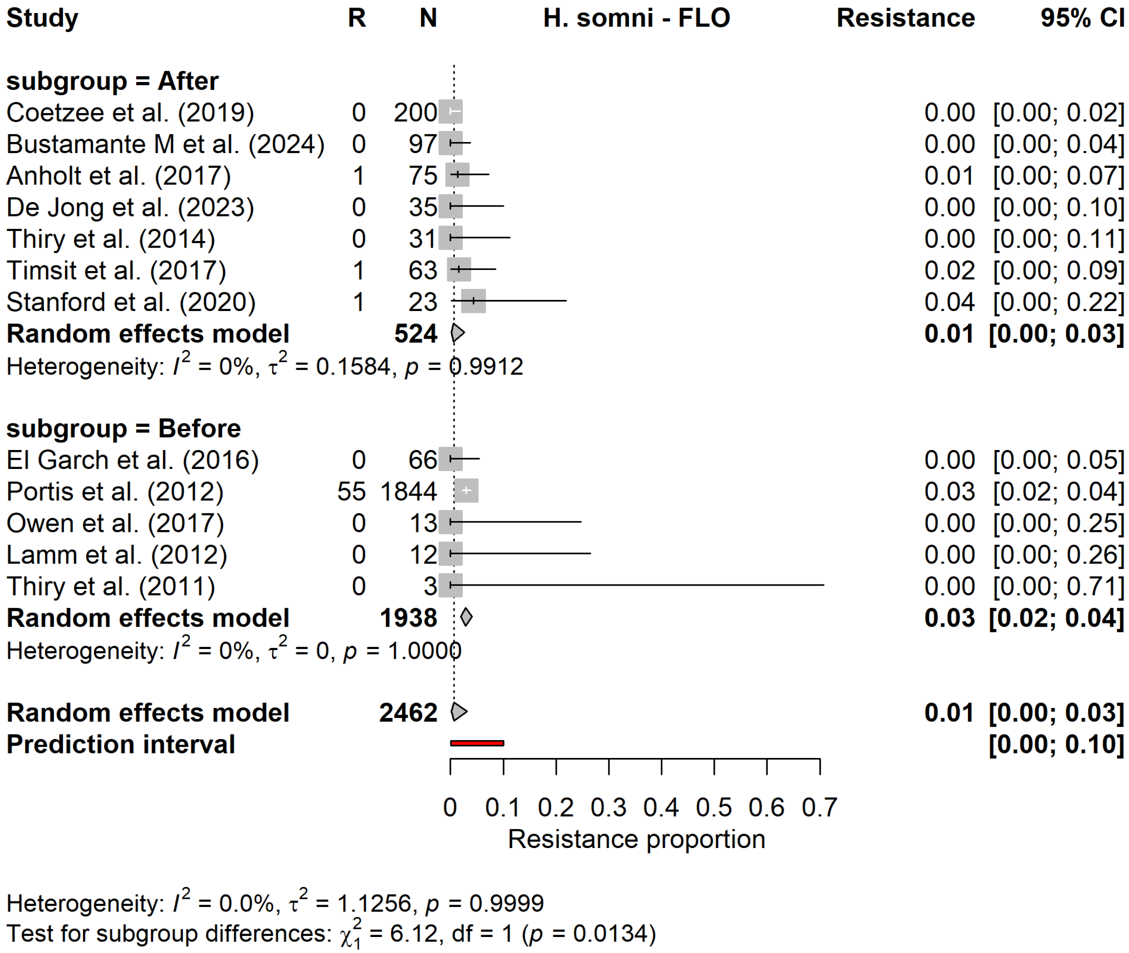


Supplementary figure S7.4. Forest plot for the subgroup meta−analysis of florfenicol resistance in Histophilus somni, stratified by study period.

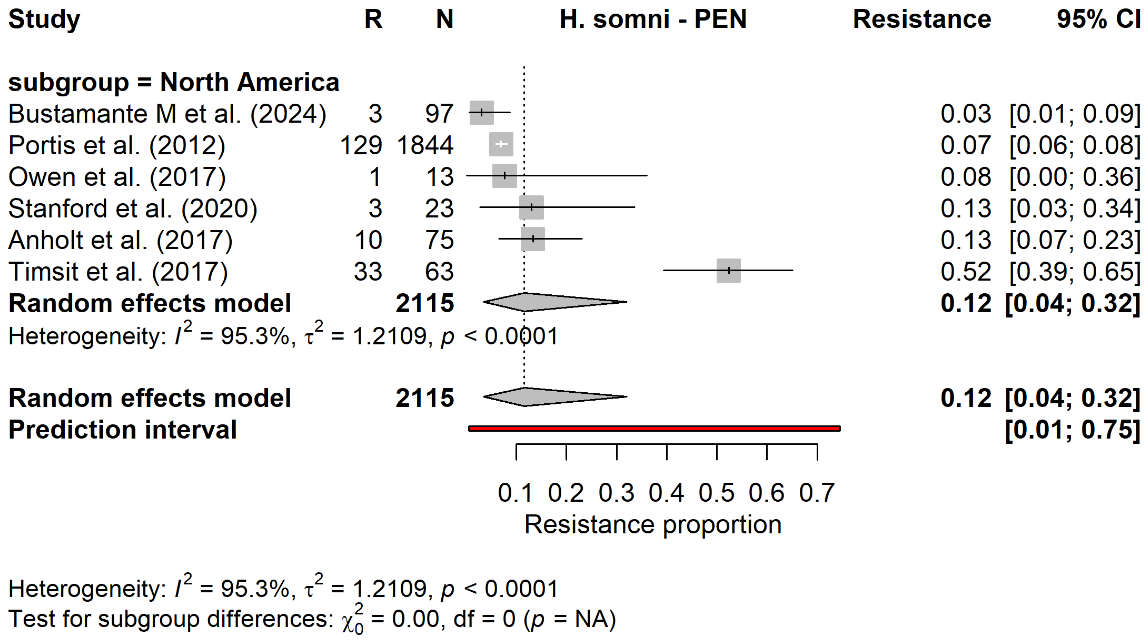


Supplementary figure S7.5. Forest plot for the subgroup meta−analysis of penicillin resistance in Histophilus somni, stratified by continent.

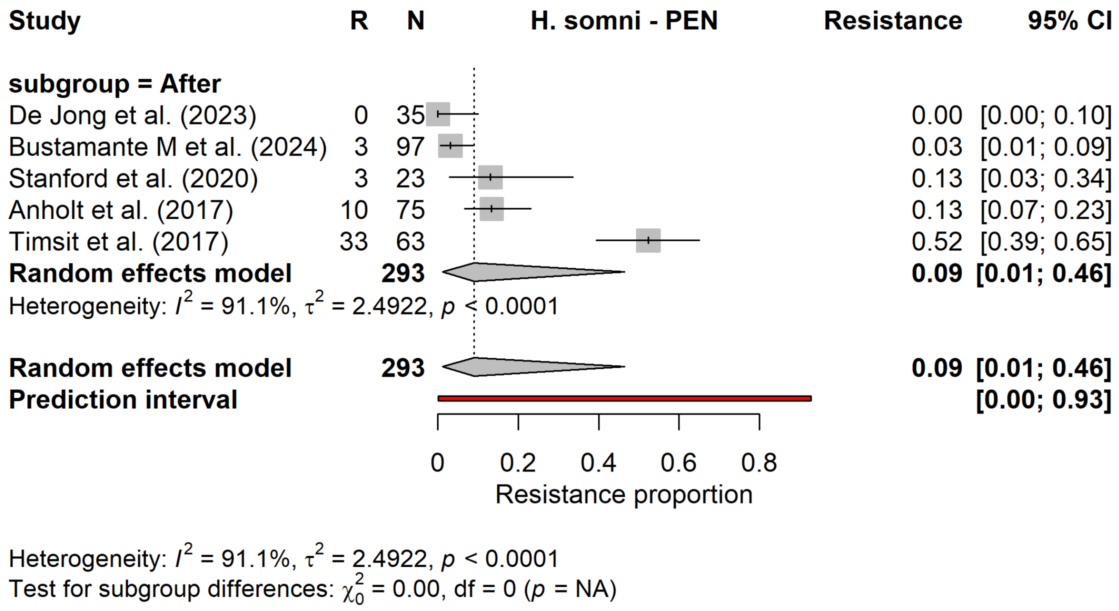


Supplementary figure S7.6. Forest plot for the subgroup meta−analysis of penicillin resistance in Histophilus somni, stratified by study period.

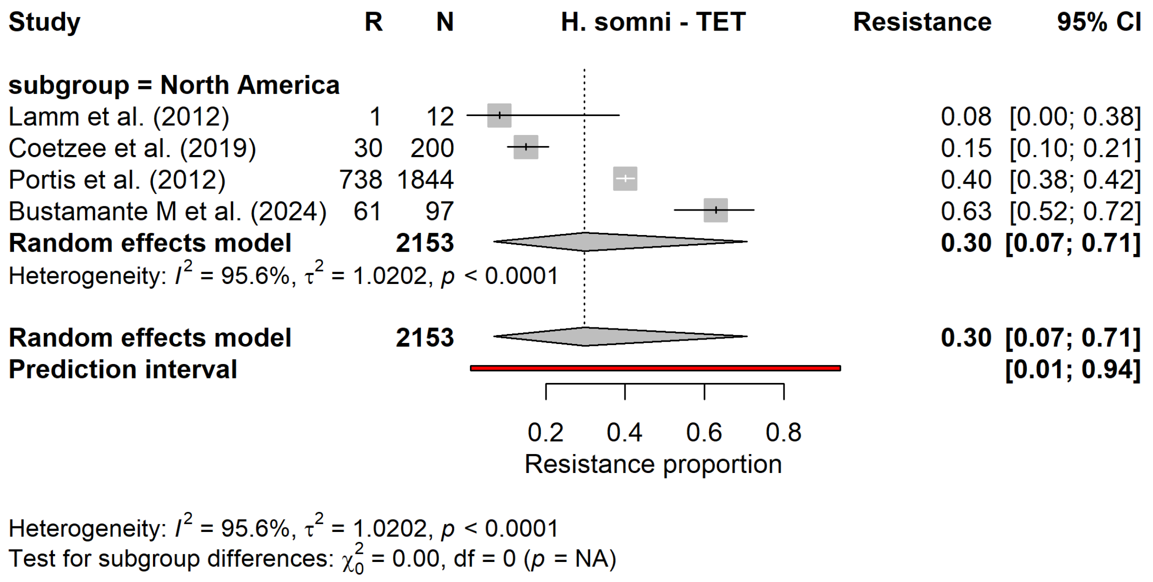


Supplementary figure S7.7. Forest plot for the subgroup meta−analysis of tetracycline resistance in Histophilus somni, stratified by continent.

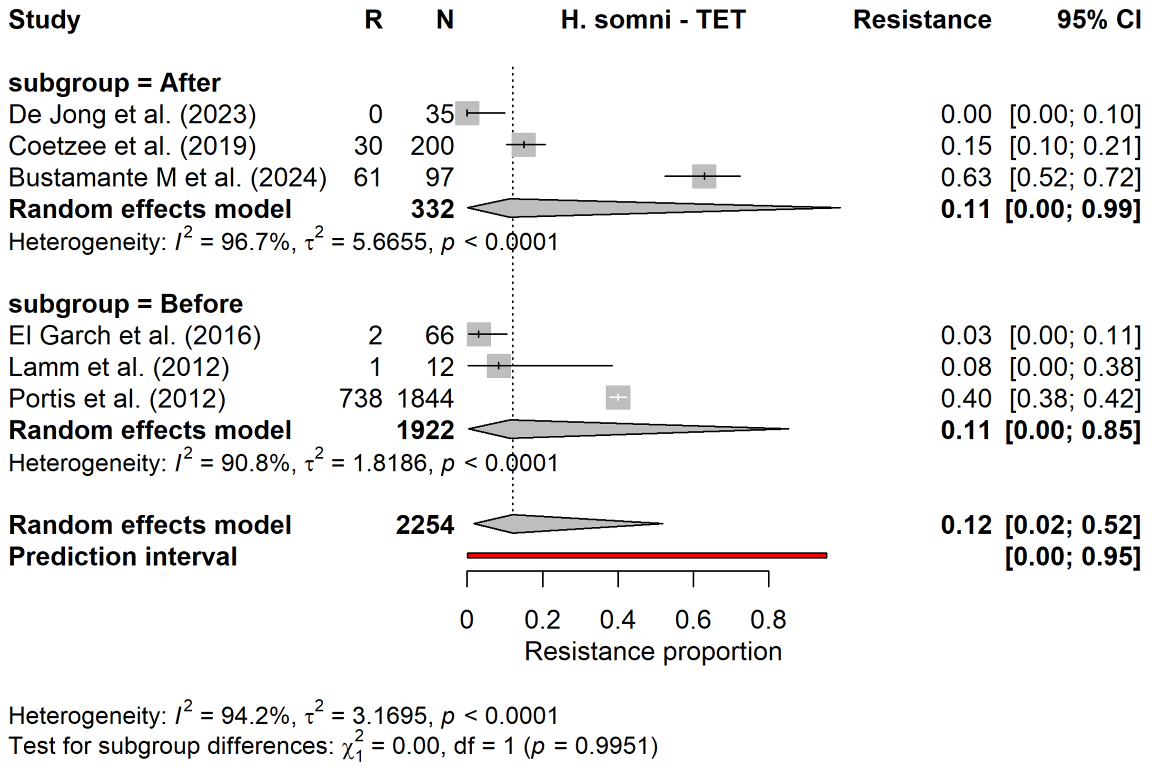


Supplementary figure S7.8. Forest plot for the subgroup meta−analysis of tetracycline resistance in Histophilus somni, stratified by study period.

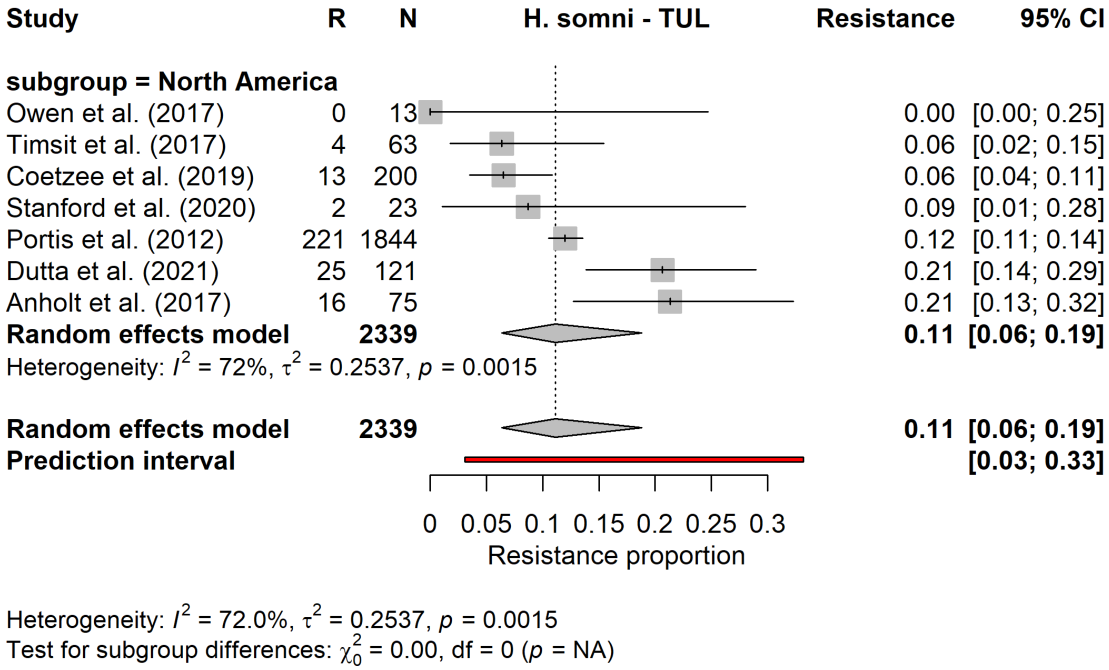


Supplementary figure S7.9. Forest plot for the subgroup meta−analysis of tulathromycin resistance in Histophilus somni, stratified by continent.

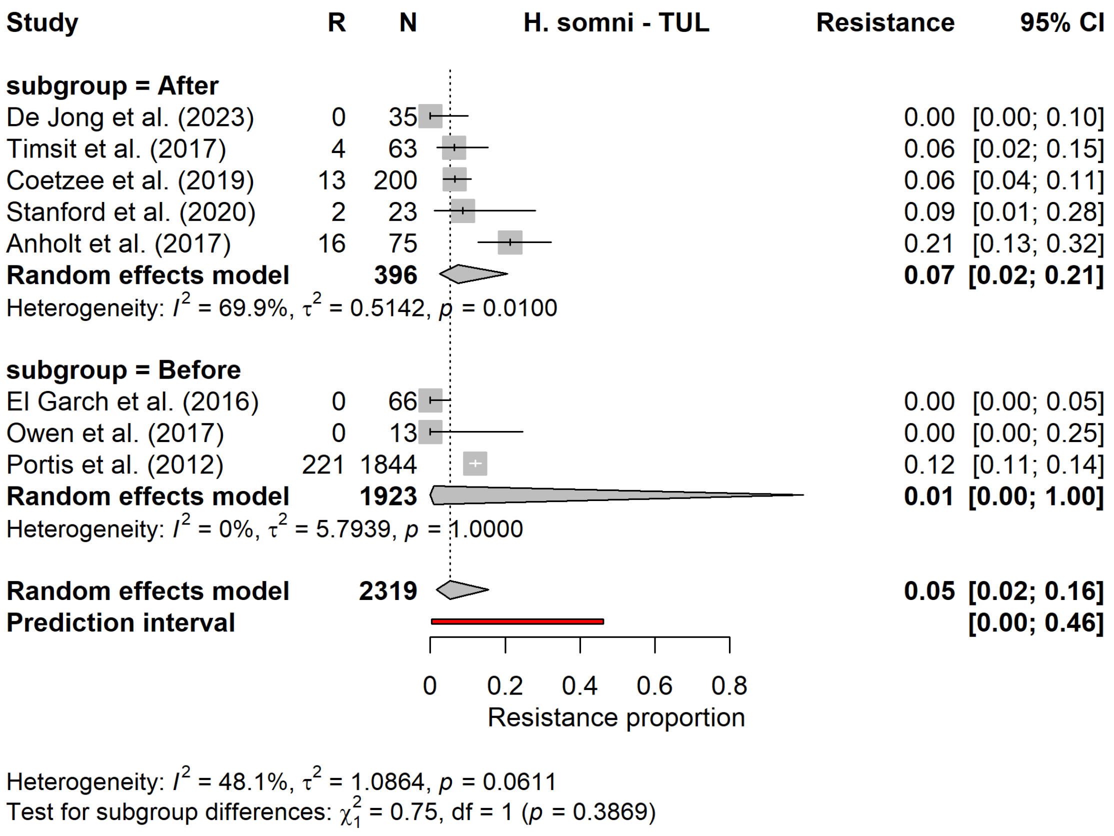


Supplementary figure S7.10. Forest plot for the subgroup meta−analysis of tulathromycin resistance in Histophilus somni, stratified by study period.

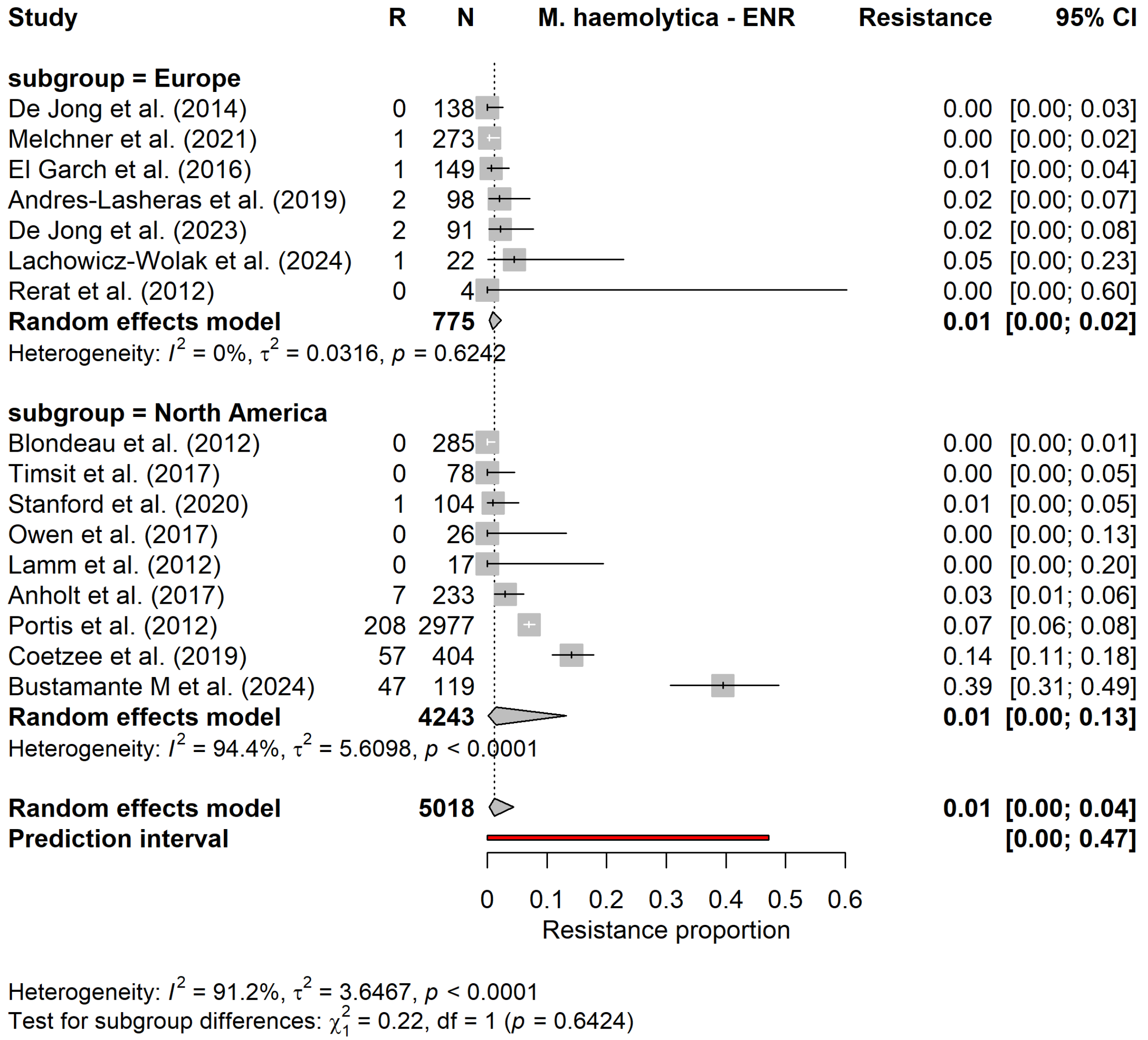


Supplementary figure S7.11. Forest plot for the subgroup meta−analysis of enrofloxacin resistance in Mannheimia haemolytica, stratified by continent.

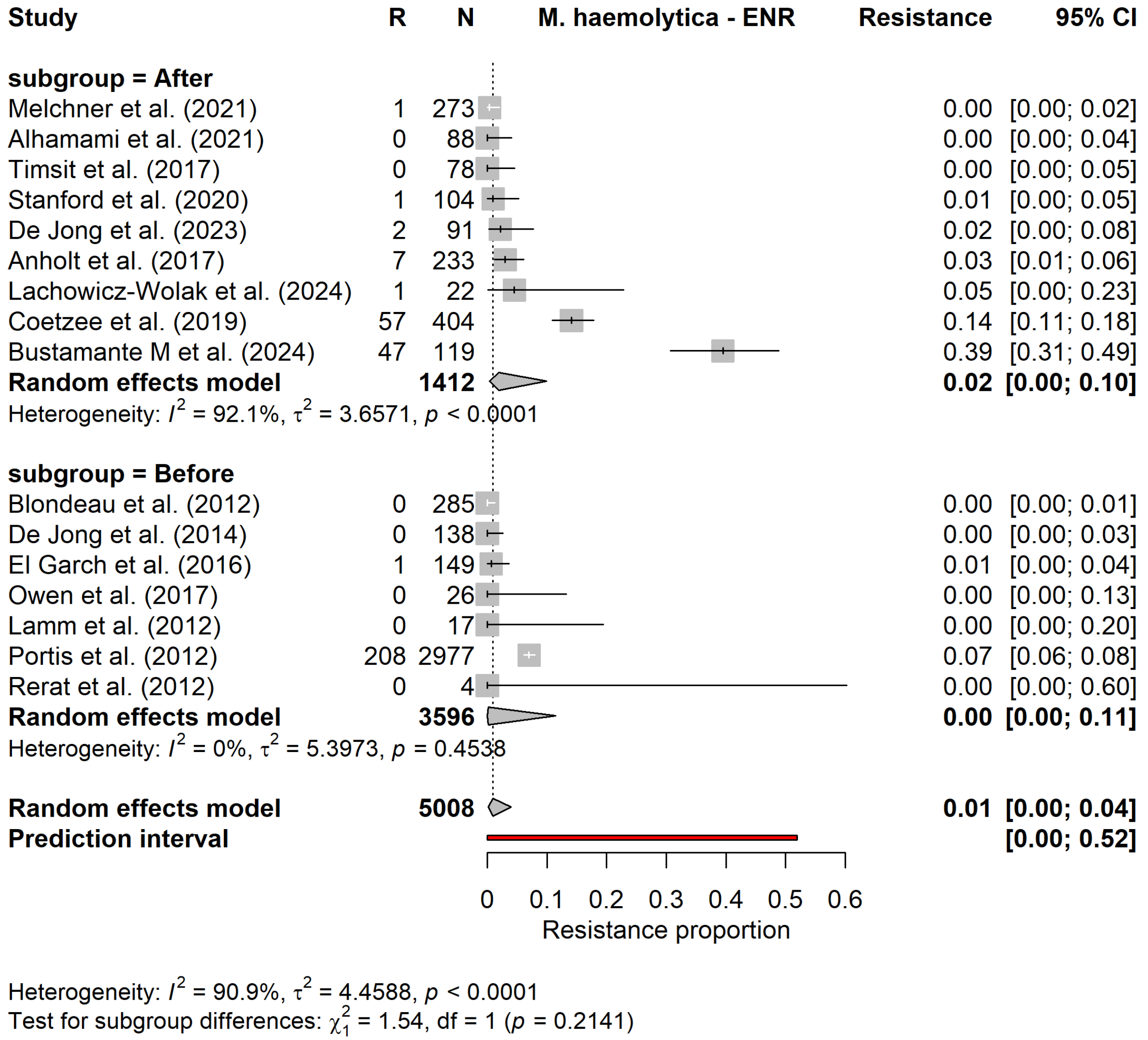


Supplementary figure S7.12. Forest plot for the subgroup meta−analysis of enrofloxacin resistance in Mannheimia haemolytica, stratified by study period.

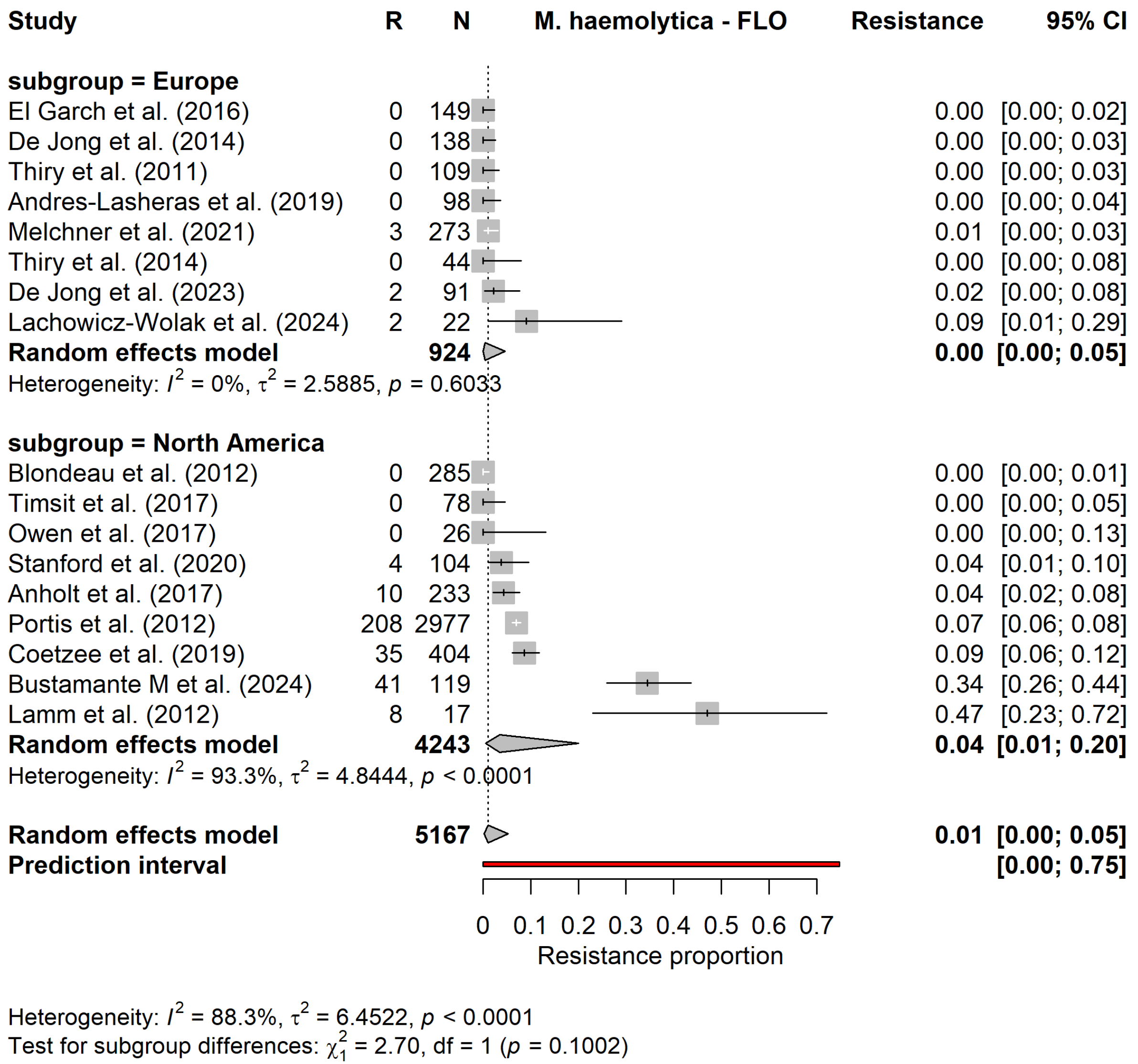


Supplementary figure S7.13. Forest plot for the subgroup meta−analysis of florfenicol resistance in Mannheimia haemolytica, stratified by continent.

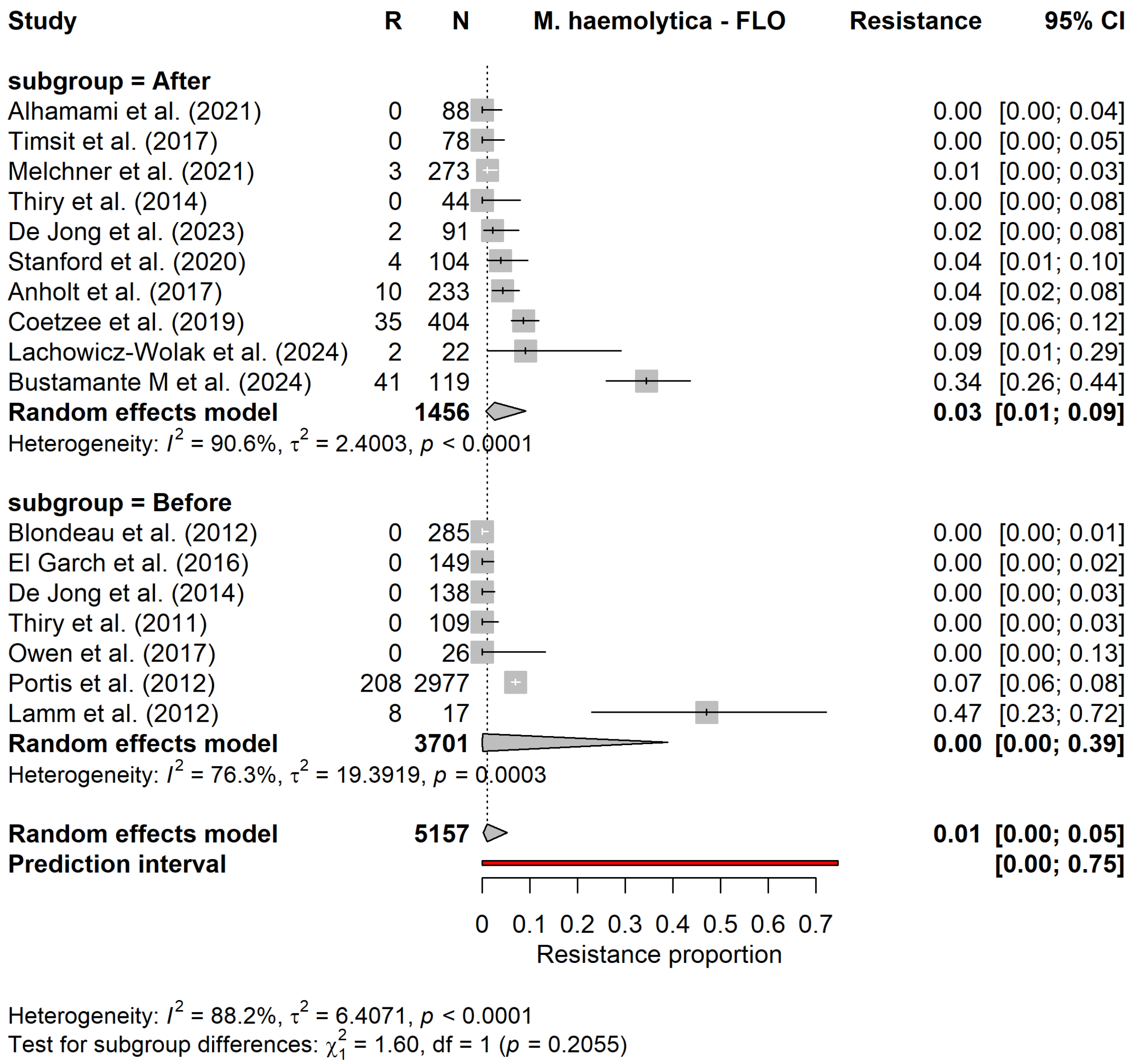


Supplementary figure S7.14. Forest plot for the subgroup meta−analysis of florfenicol resistance in Mannheimia haemolytica, stratified by study period.

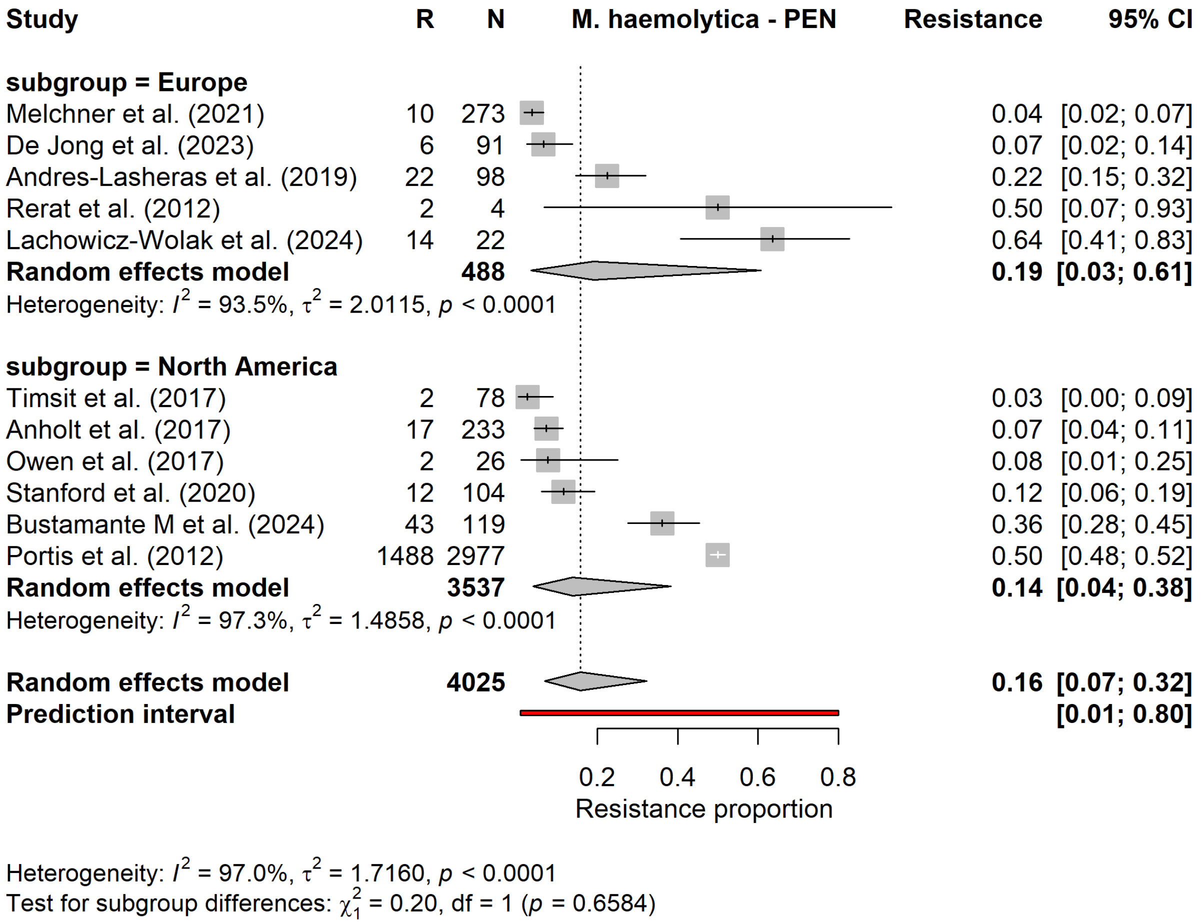


Supplementary figure S7.15. Forest plot for the subgroup meta−analysis of penicillin resistance in Mannheimia haemolytica, stratified by continent.

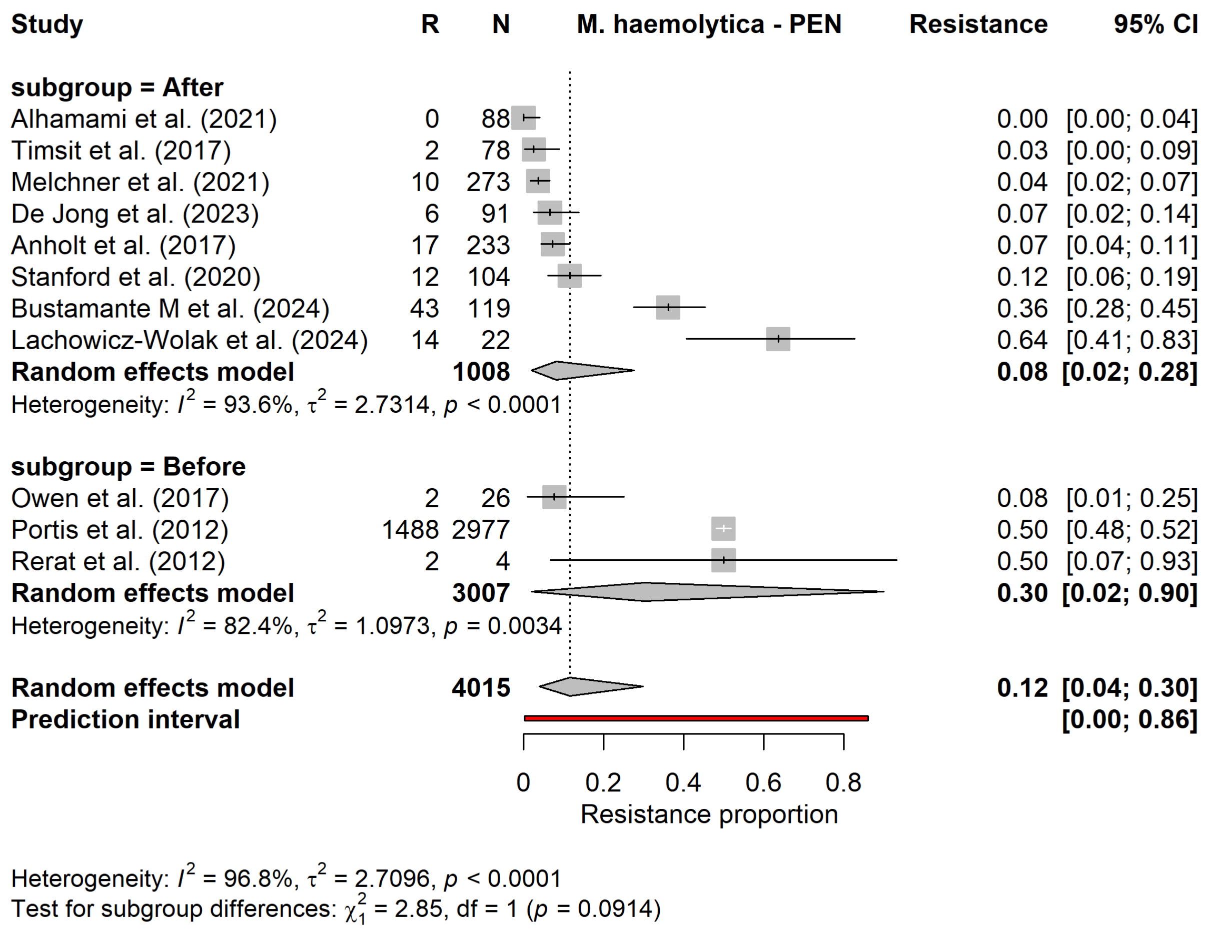


Supplementary figure S7.16. Forest plot for the subgroup meta−analysis of penicillin resistance in Mannheimia haemolytica, stratified by study period.

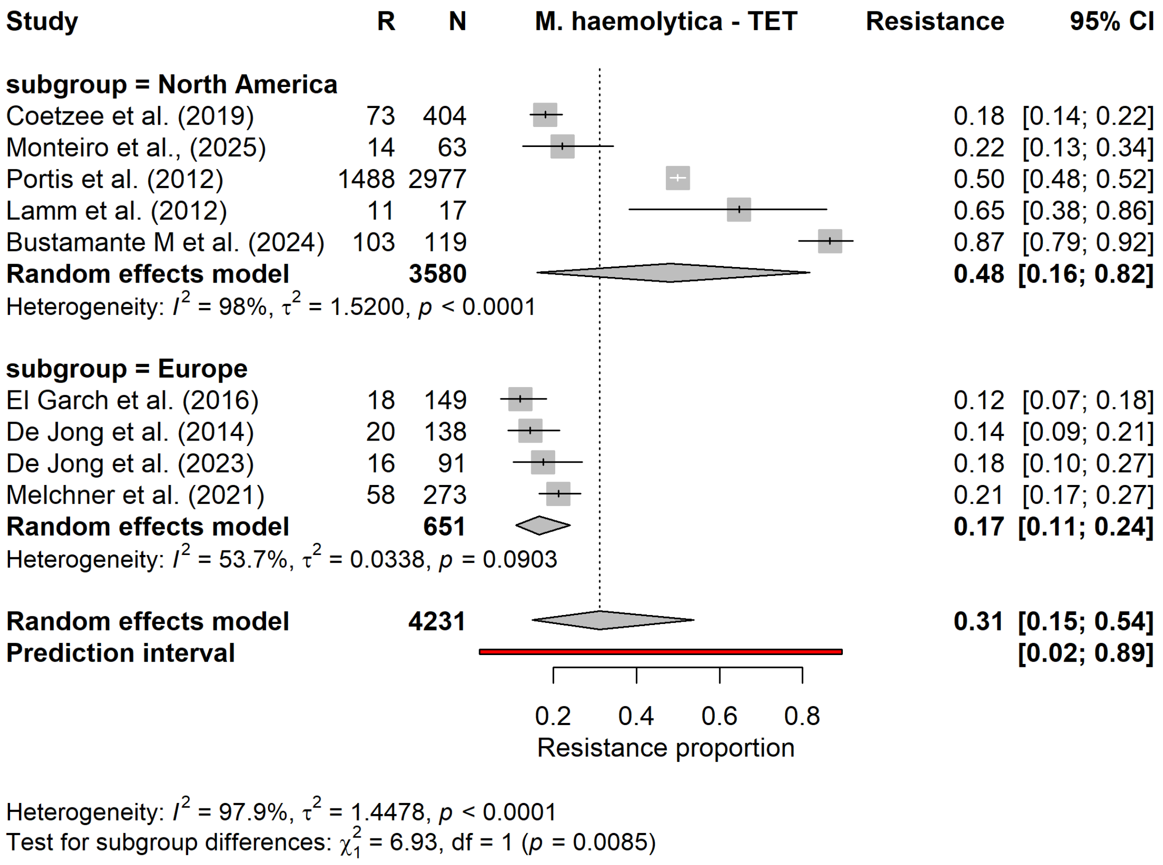


Supplementary figure S7.17. Forest plot for the subgroup meta−analysis of tetracycline resistance in Mannheimia haemolytica, stratified by continent.

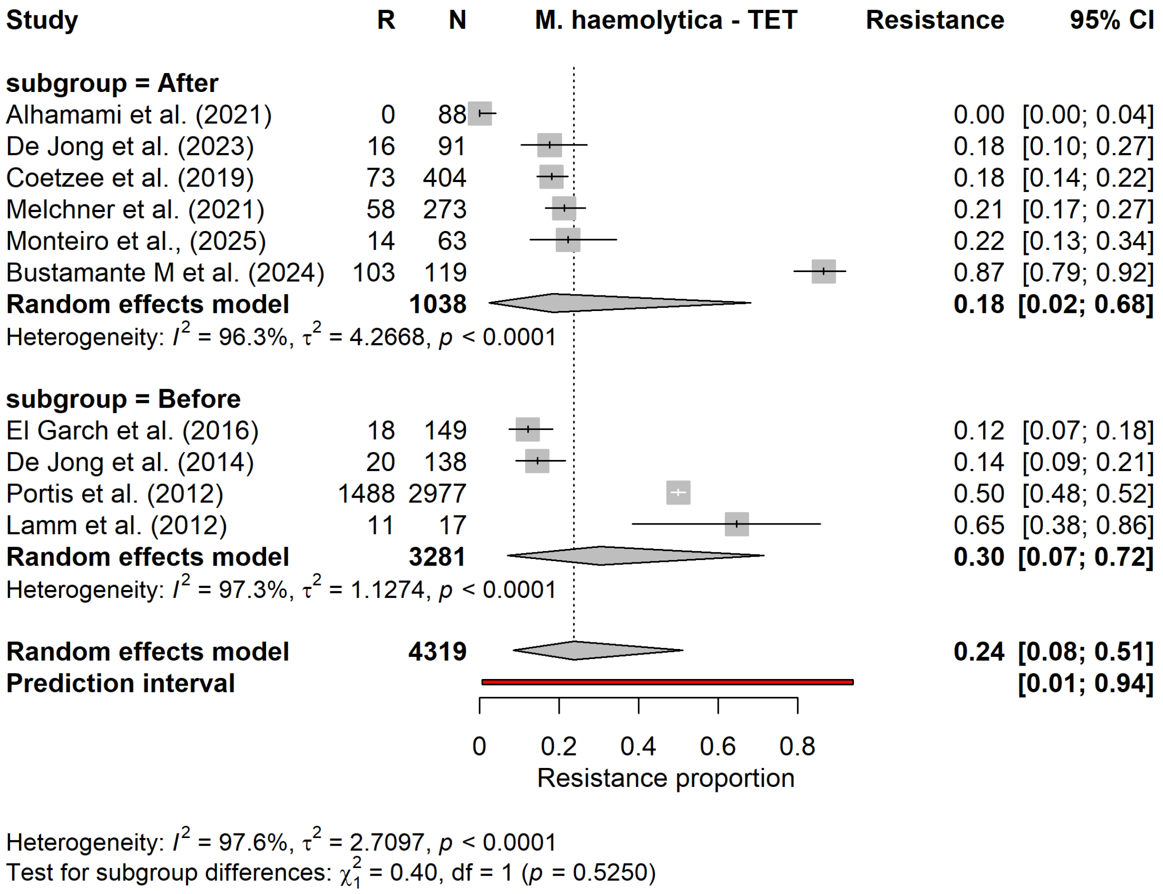


Supplementary figure S7.18. Forest plot for the subgroup meta−analysis of tetracycline resistance in Mannheimia haemolytica, stratified by study period.

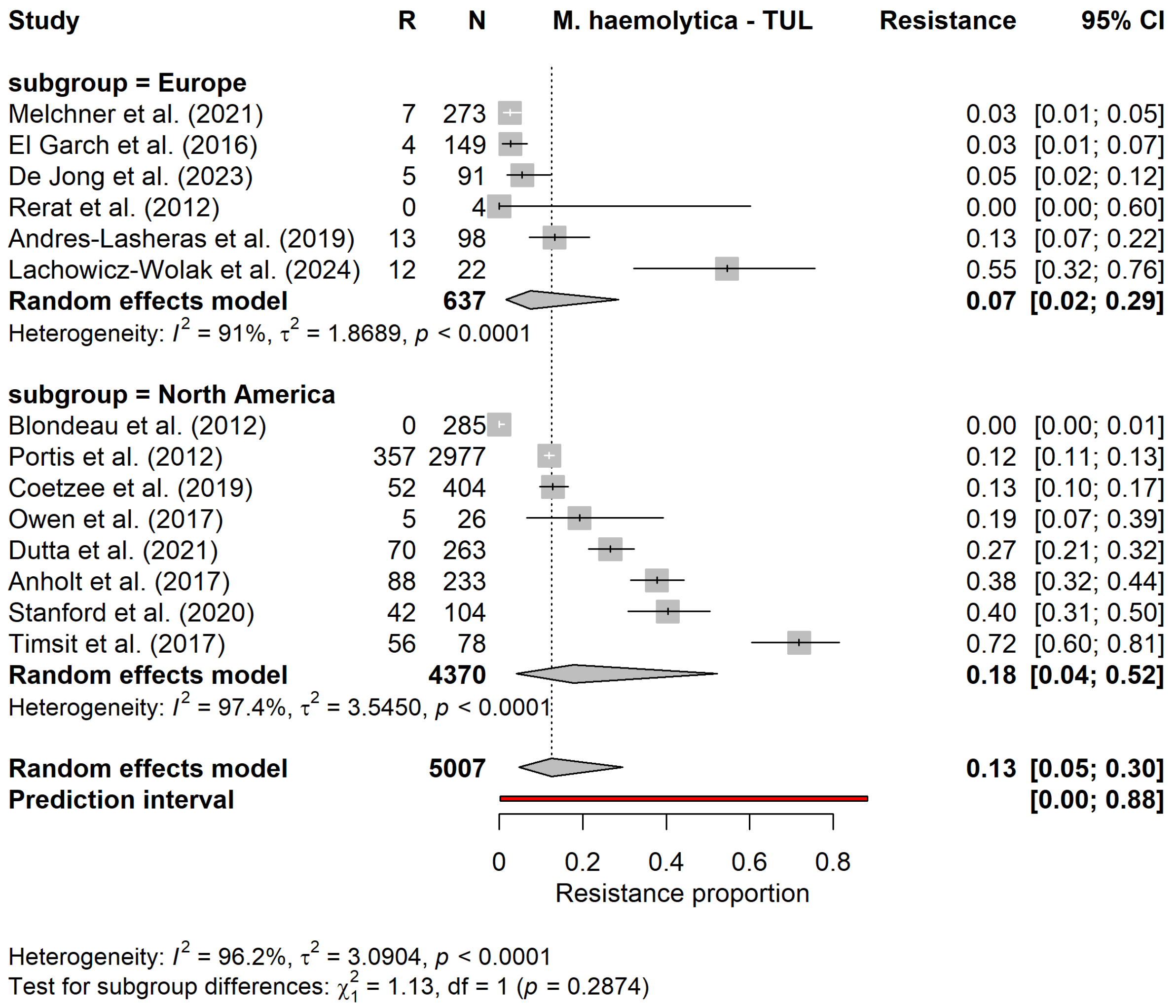


Supplementary figure S7.19. Forest plot for the subgroup meta−analysis of tulathromycin resistance in Mannheimia haemolytica, stratified by continent.

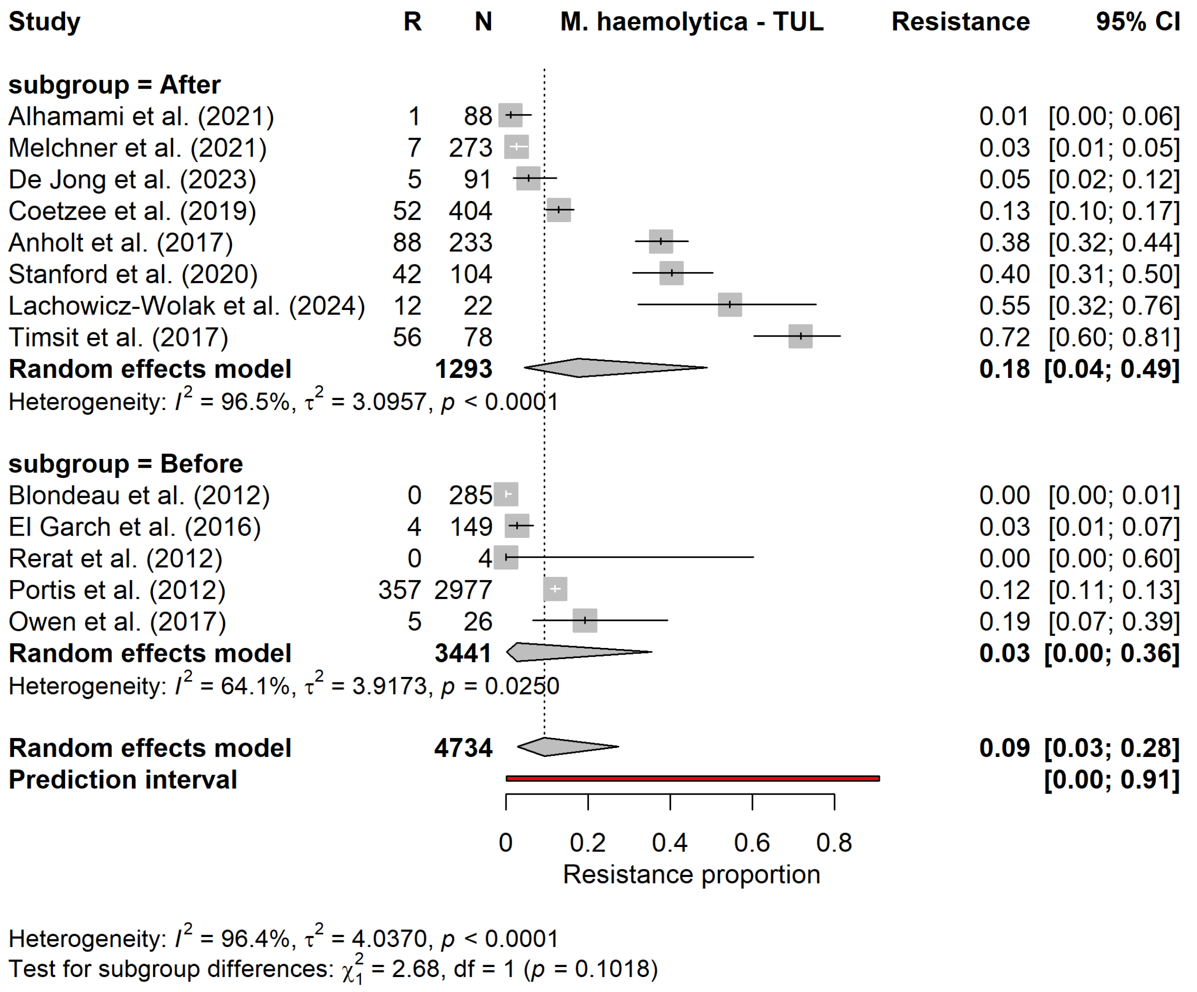


Supplementary figure S7.20. Forest plot for the subgroup meta−analysis of tulathromycin resistance in Mannheimia haemolytica, stratified by study period.

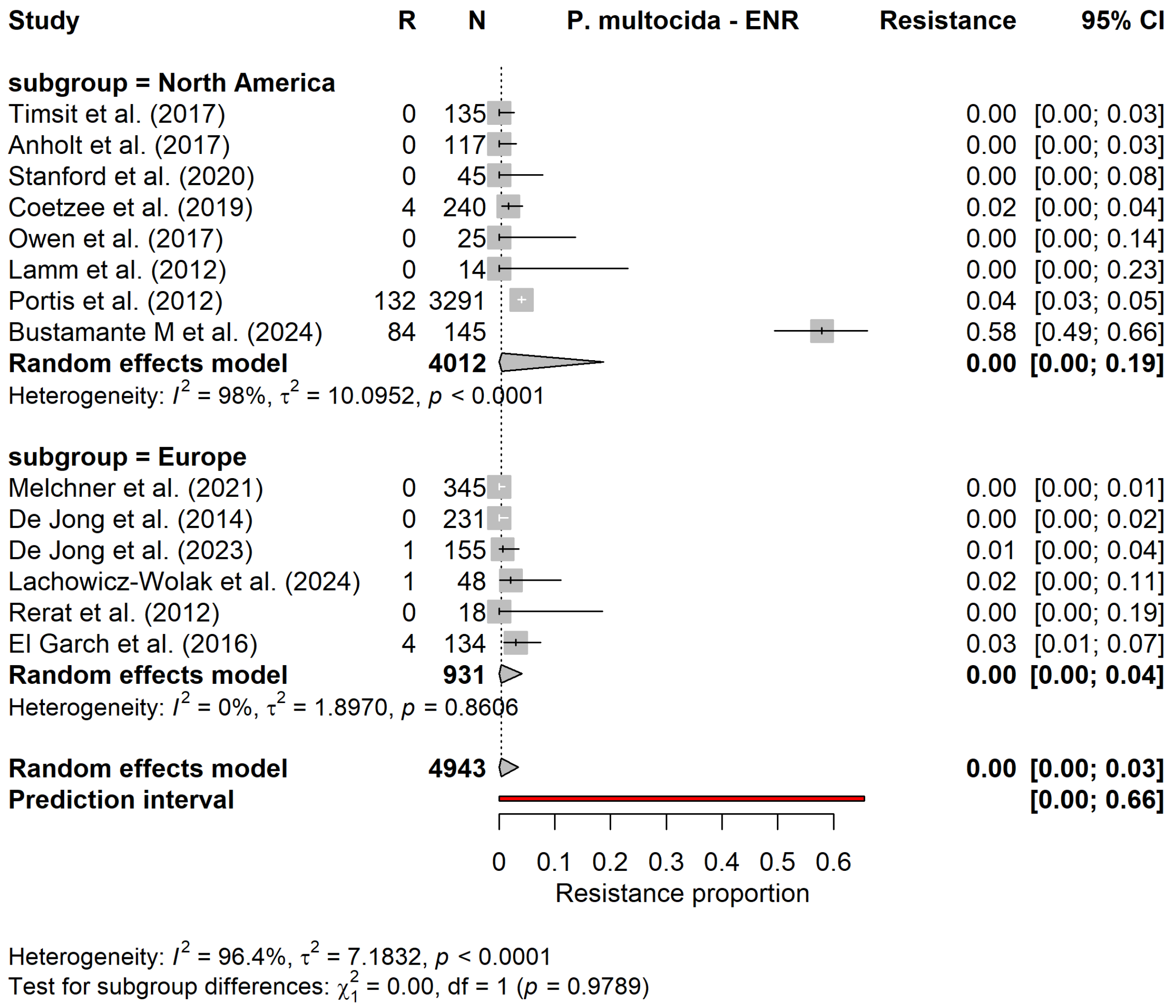


Supplementary figure S7.21. Forest plot for the subgroup meta−analysis of enrofloxacin resistance in Pasteurella multocida, stratified by continent.

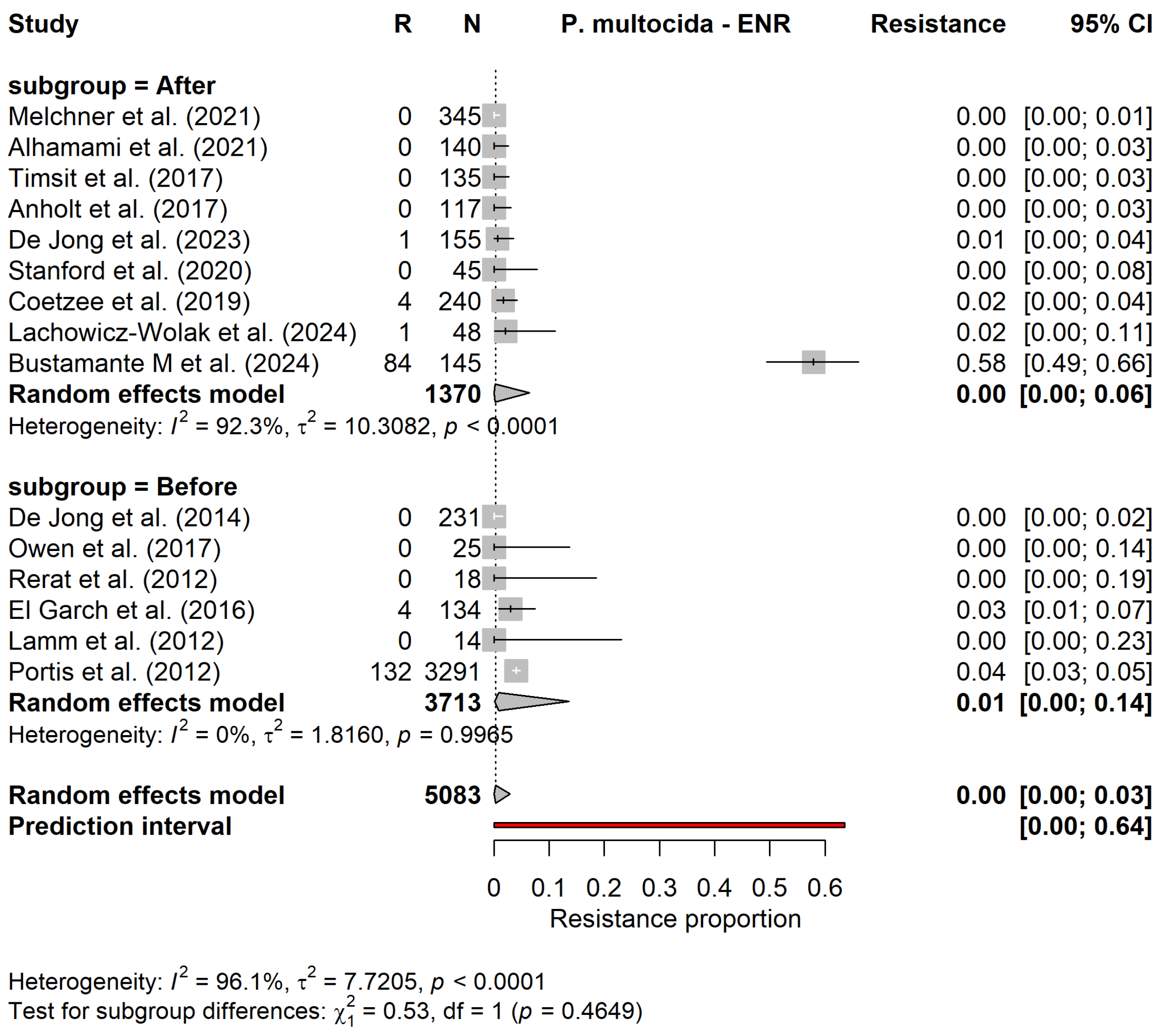


Supplementary figure S7.22. Forest plot for the subgroup meta−analysis of enrofloxacin resistance in Pasteurella multocida, stratified by study period.

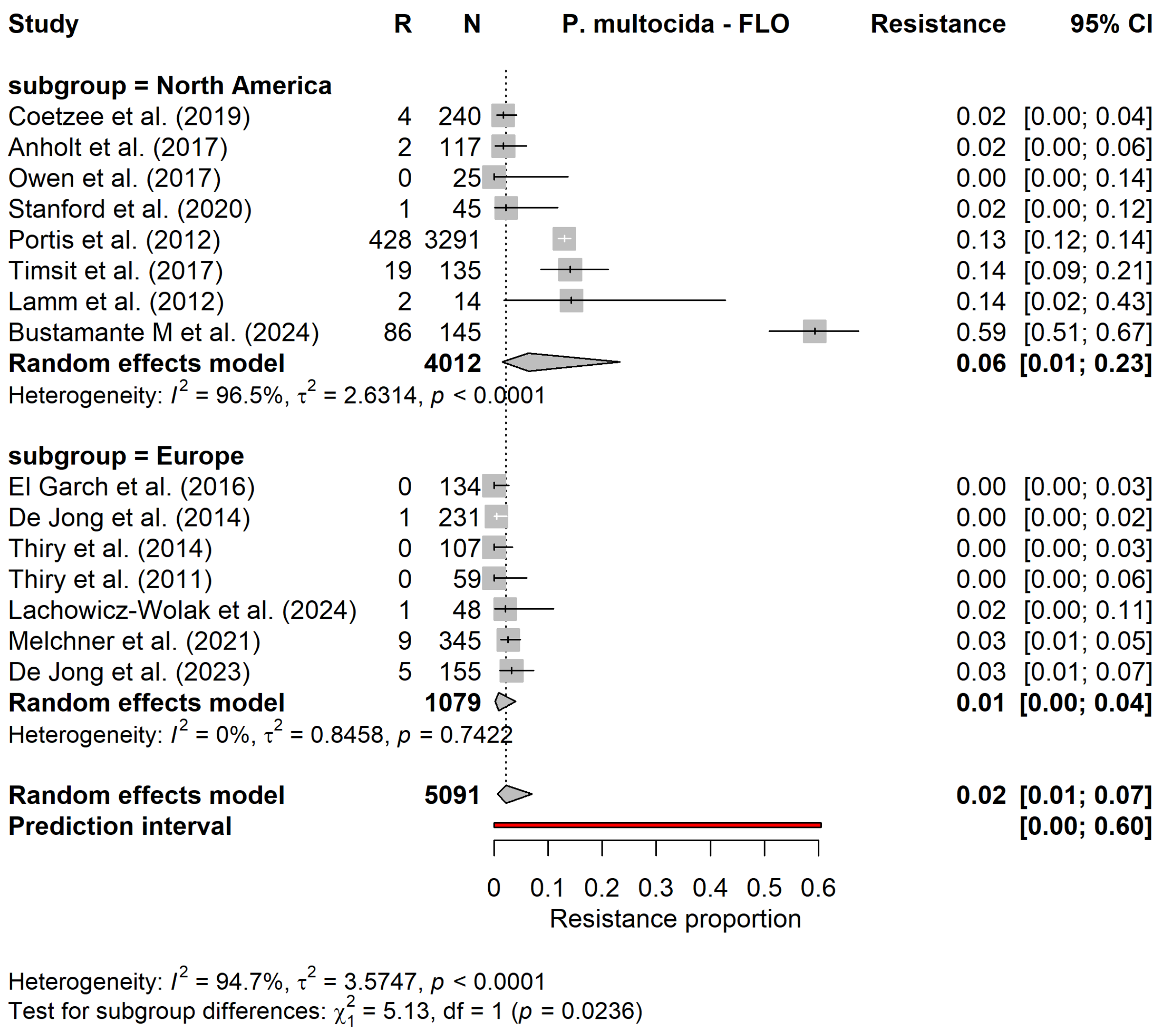


Supplementary figure S7.23. Forest plot for the subgroup meta−analysis of florfenicol resistance in Pasteurella multocida, stratified by continent.

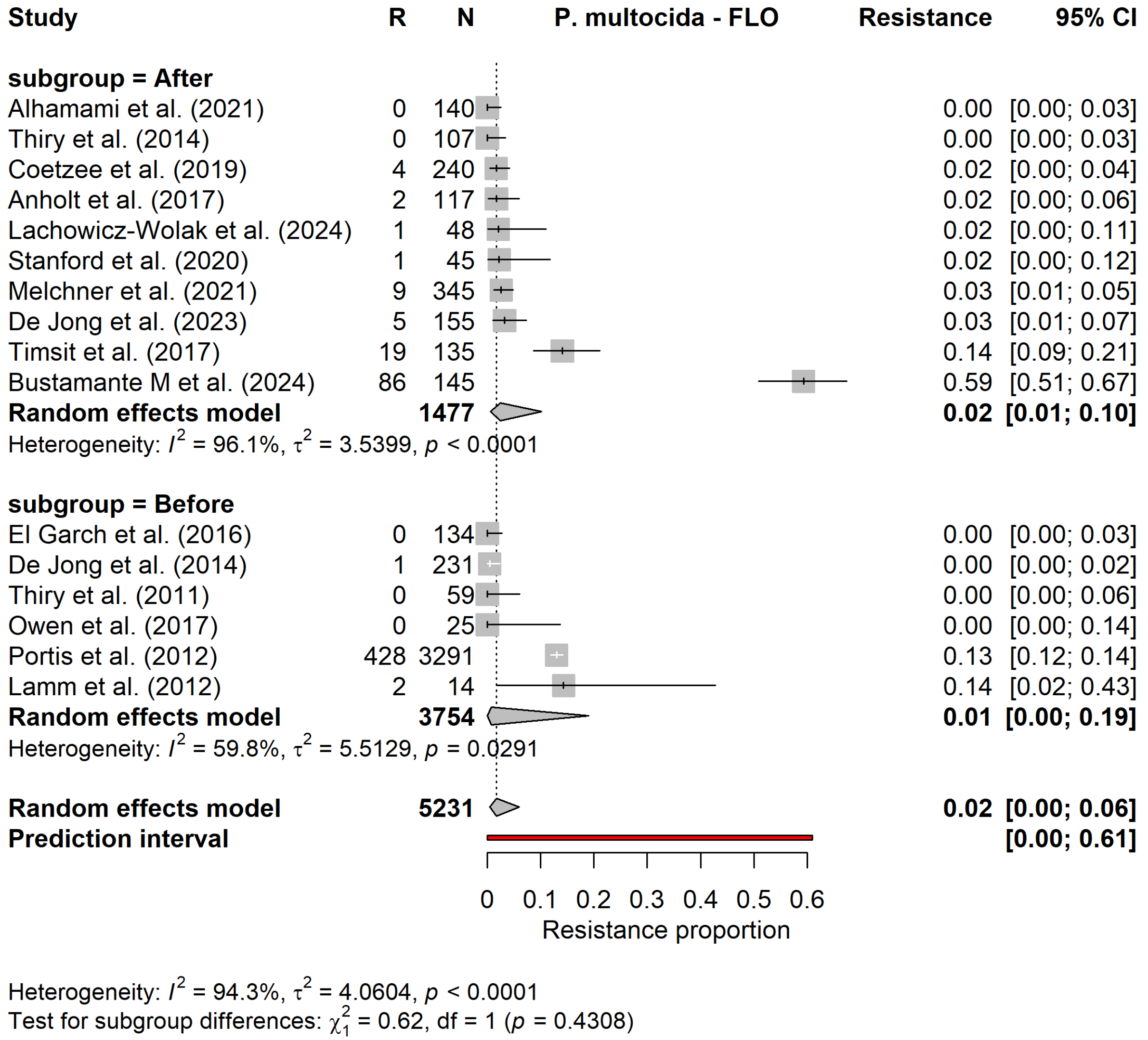


Supplementary figure S7.24. Forest plot for the subgroup meta−analysis of florfenicol resistance in Pasteurella multocida, stratified by study period.

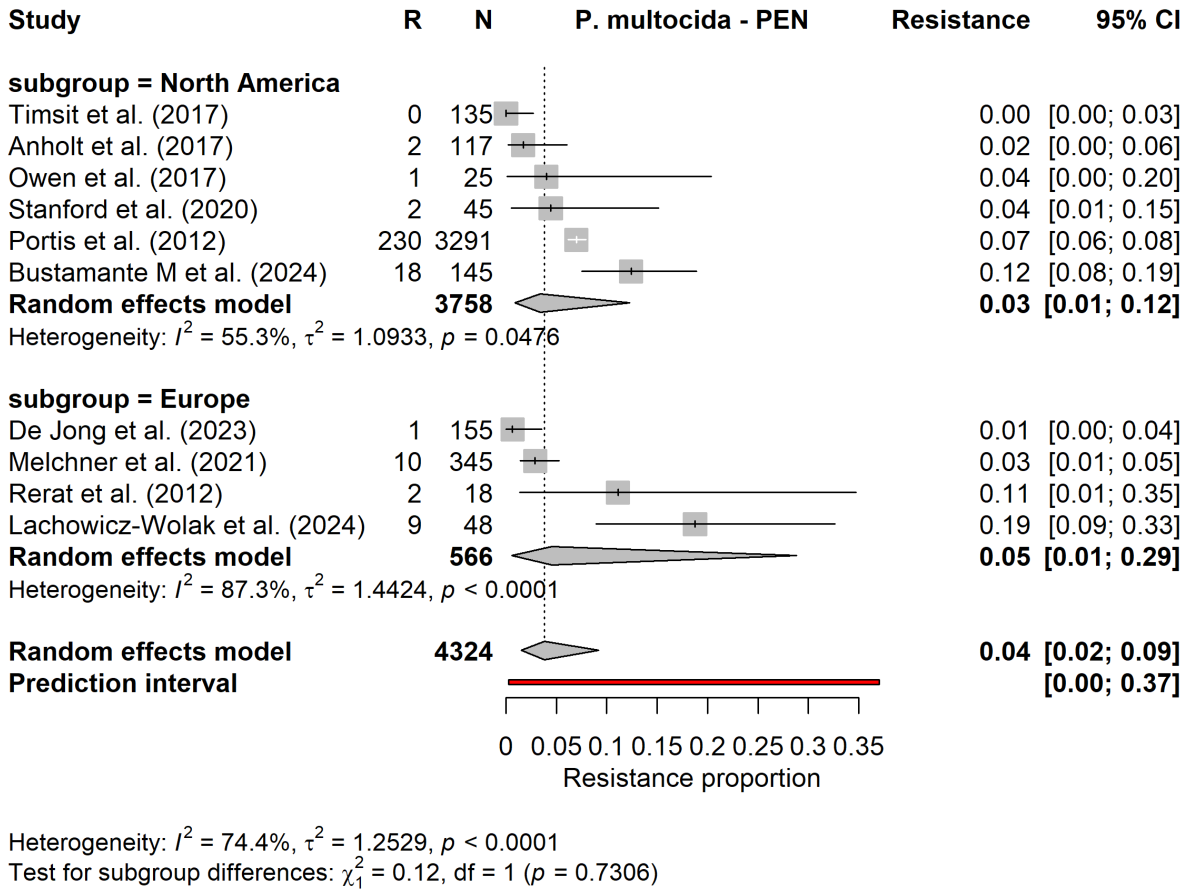


Supplementary figure S7.25. Forest plot for the subgroup meta−analysis of penicillin resistance in Pasteurella multocida, stratified by continent.

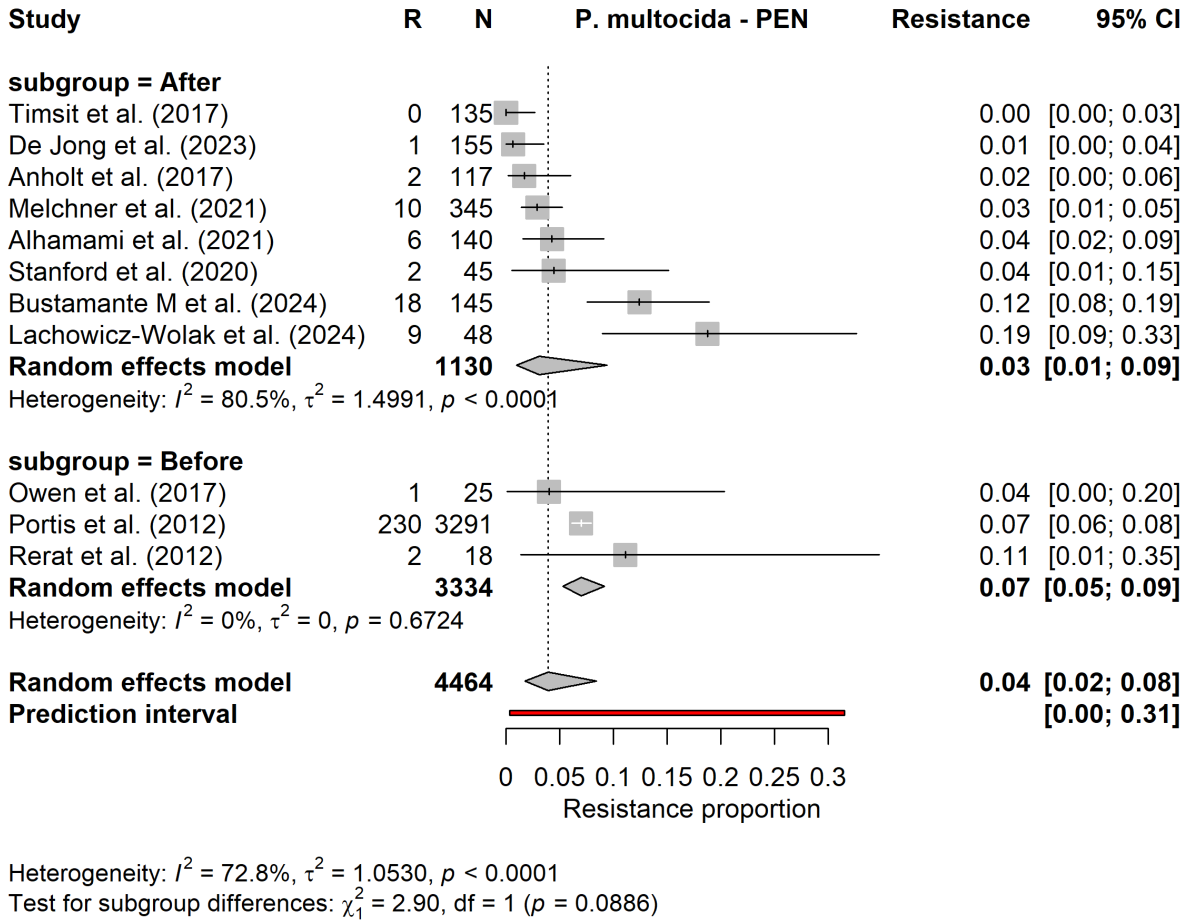


Supplementary figure S7.26. Forest plot for the subgroup meta−analysis of penicillin resistance in Pasteurella multocida, stratified by study period.

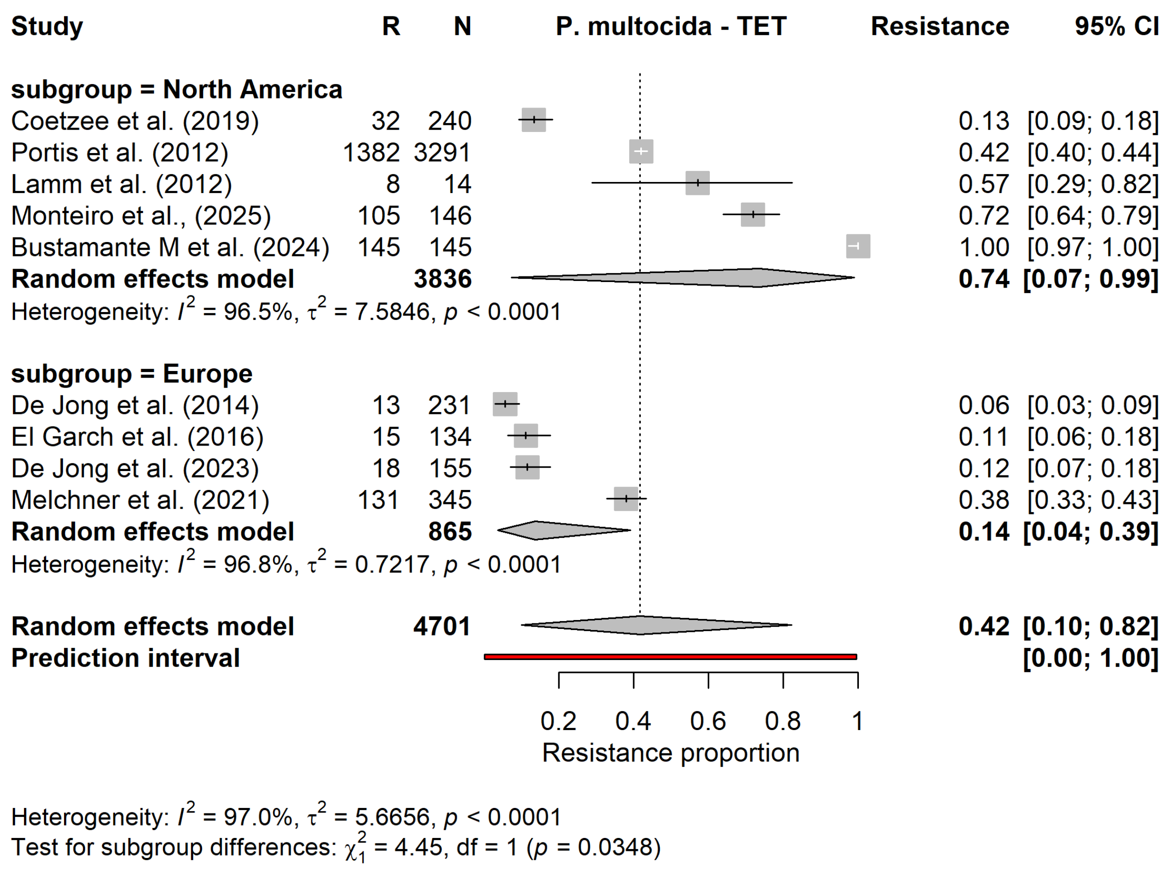


Supplementary figure S7.27. Forest plot for the subgroup meta−analysis of tetracycline resistance in Pasteurella multocida, stratified by continent.

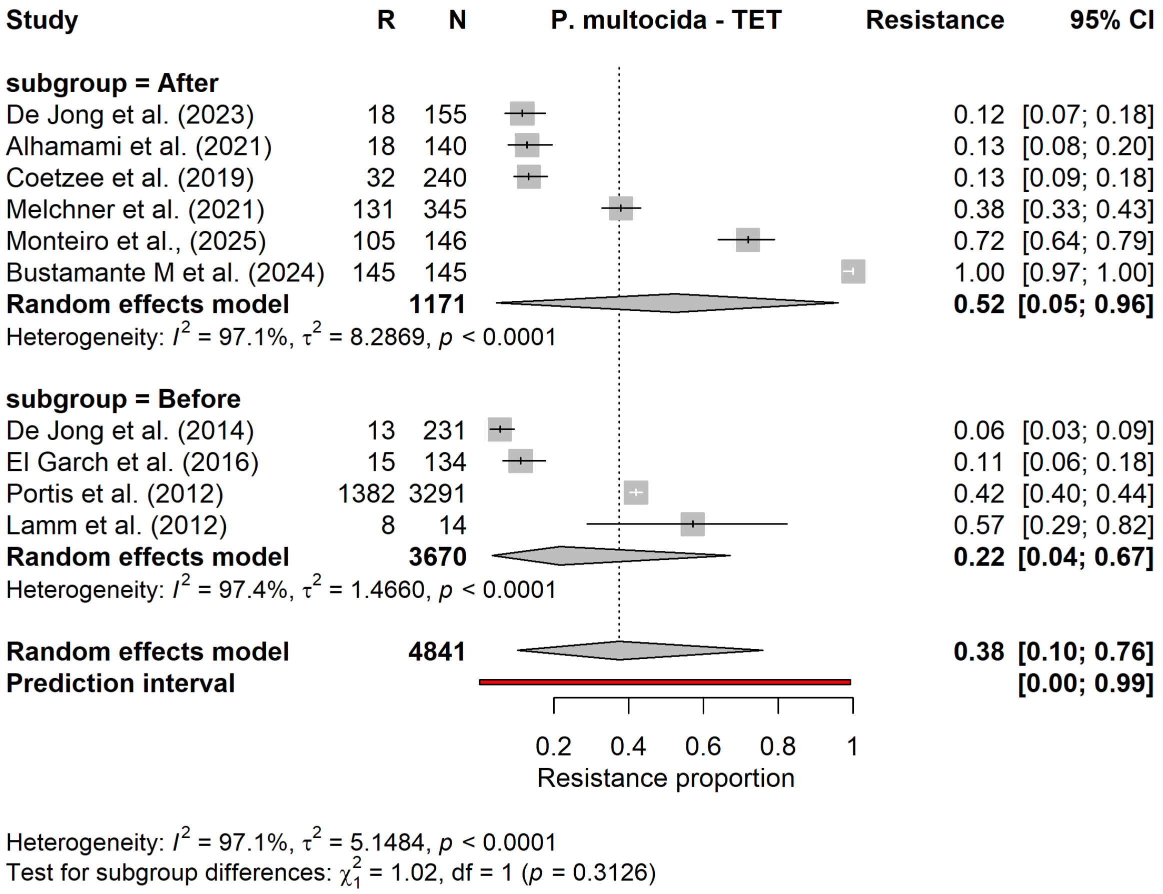


Supplementary figure S7.28. Forest plot for the subgroup meta−analysis of tetracycline resistance in Pasteurella multocida, stratified by study period.

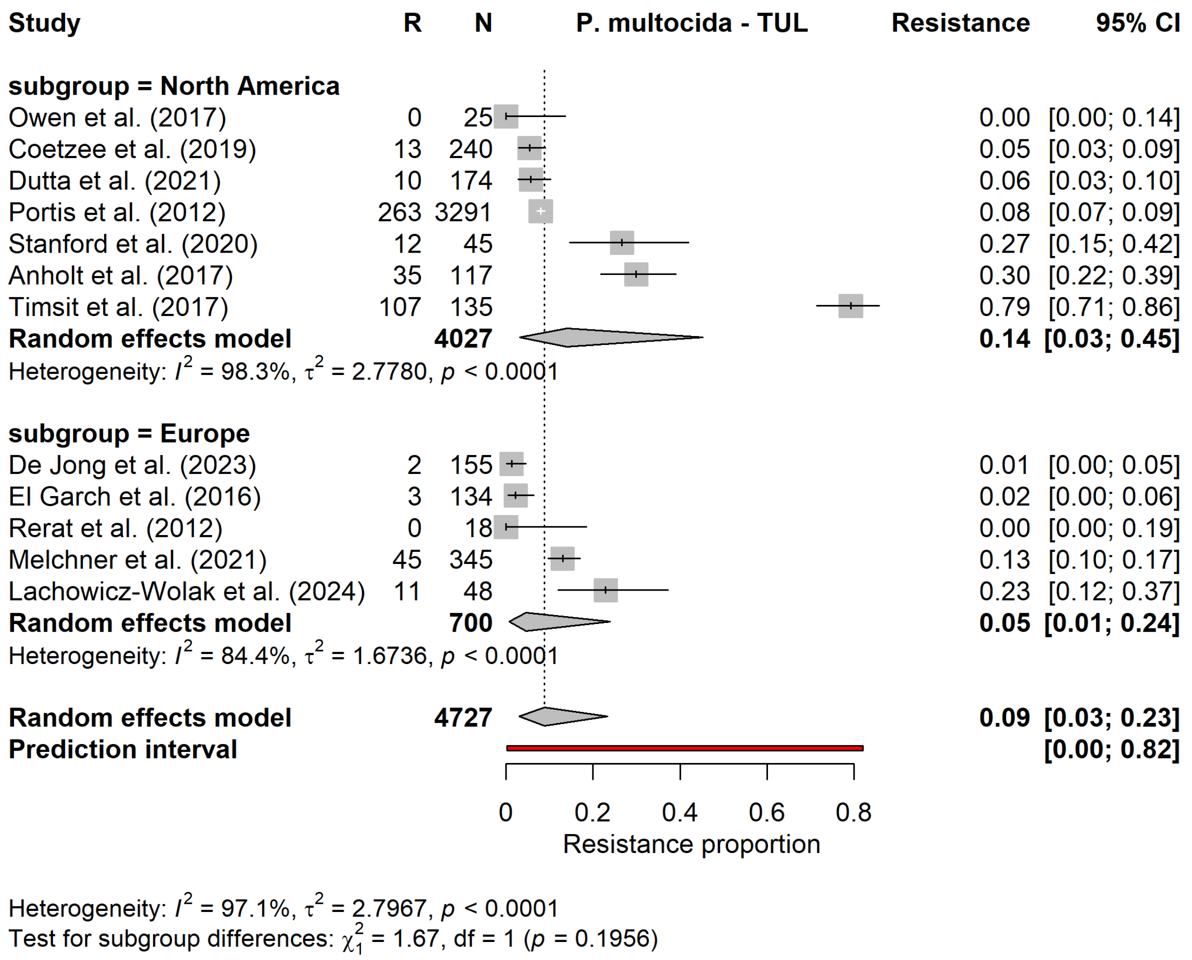


Supplementary figure S7.29. Forest plot for the subgroup meta−analysis of tulathromycin resistance in Pasteurella multocida, stratified by continent.

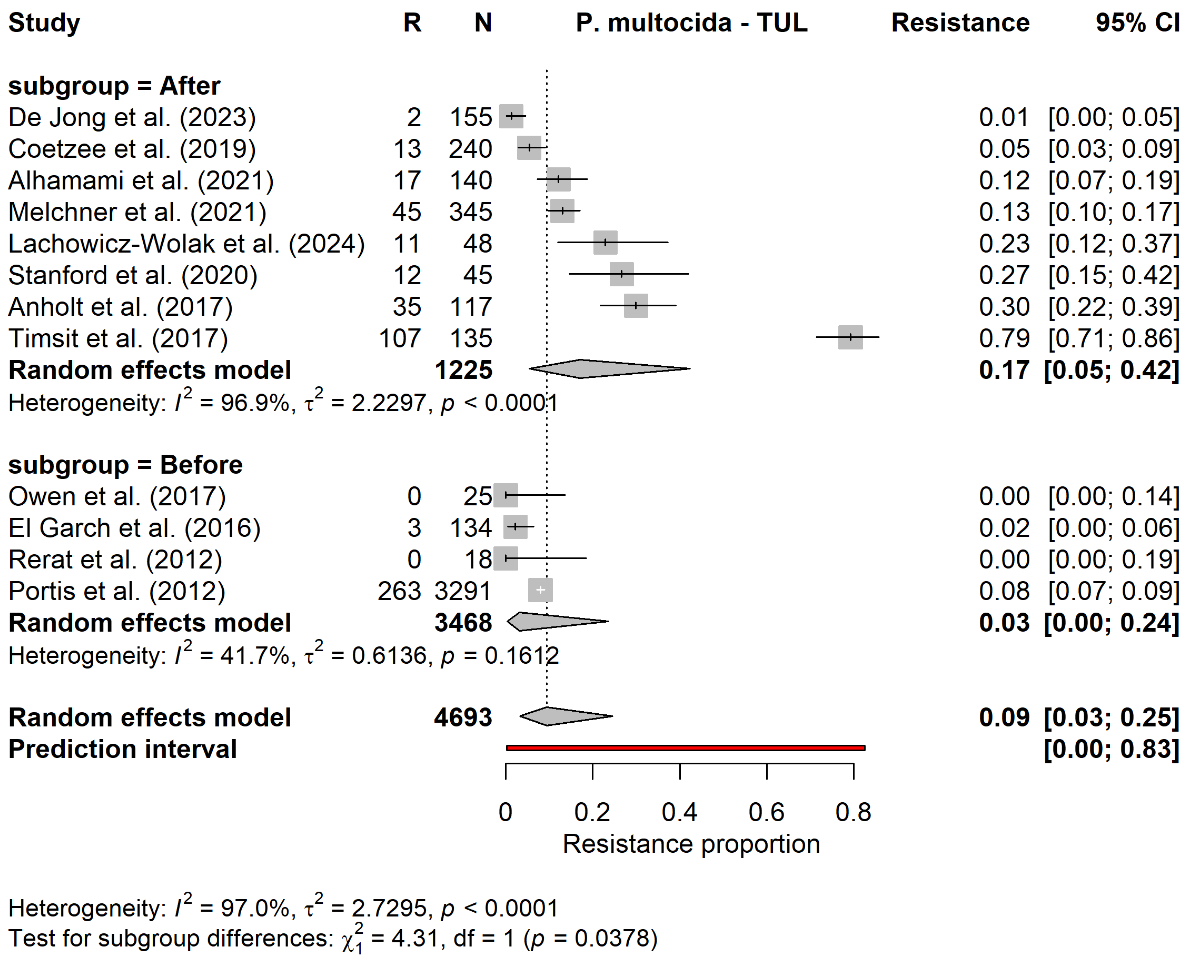


Supplementary figure S7.30. Forest plot for the subgroup meta−analysis of tulathromycin resistance in Pasteurella multocida, stratified by study period.
